# Interpretable phenotype decoding from multi-condition sequencing data with ALPINE

**DOI:** 10.1101/2025.02.15.638471

**Authors:** Wei-Hao Lee, Lechuan Li, Ruth Dannenfelser, Vicky Yao

## Abstract

As sequencing techniques advance in precision, affordability, and diversity, an abundance of heterogeneous sequencing data has become available, encompassing a wide range of phenotypic features and biological perturbations. Unfortunately, increased resolution comes with a cost of increased complexity of the biological search space, even at the individual study level, as perturbations are now often examined across many dimensions simultaneously, including different: donor phenotypes, anatomical regions and cell types, and time points. Furthermore, broad integration across studies promise unique opportunity to explore the molecular underpinnings of distinct healthy and disease states, larger than the original scope of the individual study. To fully realize the promise of both individual higher resolution studies and large cross-study integrations we need a robust methodology that can disentangle the influence of technical and non-relevant phenotypic factors, isolating relevant condition-specific signals from shared biological information while also providing interpretable insights into the genetic effects of these conditions. Current methods typically excel in only one of these areas. To address this gap, we developed ALPINE, a supervised non-negative matrix factorization (NMF) framework that effectively separates both technical and non-technical factors while simultaneously offering direct interpretability of condition-associated genes. Through simulations across 4 different scenarios, we demonstrate that ALPINE outperforms existing methods in both isolating the effect of different phenotypic conditions and prioritizing condition-associated genes. Furthermore, ALPINE has favorable performance in batch effect removal compared with state-of-the-art integration methods. When applied to real-world case studies, we showcase how ALPINE can be used to extract insights into the biological mechanisms that underlie differences between phenotypic conditions.

## Introduction

Biological complexity is incredibly multidimensional. Complex diseases impact different tissues and cell types in unique ways (Melms et al. 2021; Zeng et al. 2022; Kamath et al. 2022), and biological factors such as patient sex can also influence disease response (Ober et al. 2008; Belonwu et al. 2022; Z Huang et al. 2021). The rise of single cell technologies has fueled excitement for atlas-scale initiatives that capture cell-level diversity across numerous variables, but has also highlighted the challenges of data harmonization and interpretability (Mereu et al. 2020; Rozenblatt-Rosen et al. 2021).

The immediate recognition of a need for harmonization methods focused on removing unwanted technical variation (e.g., single cell platform, experimental laboratory) has driven substantial method development efforts (HTN Tran et al. 2020; Luecken et al. 2022), which typically aim to project cells from different datasets or samples into an integrated space (Eisenstein 2020; Argelaguet et al. 2021). However, during this integration process, the focus on aligning cell types can come at the cost of treating inter-individual variation as a batch effect that is removed (Luecken et al. 2022). Thus, while these approaches can be effective at aligning cell types to capture consensus signals across populations, they risk obscuring important, nested condition signals, such as sex- or tissue-specific differences with respect to disease state.

Recent realization that data harmonization can potentially come at the sacrifice of some interpretability has led to method development efforts to build disentangled representations of scRNA-seq data by explicitly modeling the batch effects and biological conditions during the integration process (Qian et al. 2022; Weinberger et al. 2023; Z Zhang et al. 2024; Piran et al. 2024; R Liu et al. 2024; Zhao et al. 2024). This modeling approach aims to enable the identification of condition-associated genes as well as more principled batch effect removal, though this task becomes more challenging when there are multiple groups of batches and conditions (as opposed to multiple levels within a single condition). For example, scINSIGHT (Qian et al. 2022) uses matrix factorization to facilitate interpretable representation decomposition, while contrastiveVI (Weinberger et al. 2023) uses a variational autoencoder framework to capture differences between conditions. However, these methods are inherently limited to handling different variables within a single condition. When there are multiple biological conditions, they need to be represented as a concatenation of all combinations of variables (e.g., tissue+sex), which makes it challenging to isolate and interpret individual conditions, let alone comparisons between conditions. Meanwhile, though methods such as scDisInFact (Z Zhang et al. 2024) and scDisco (R Liu et al. 2024) can explicitly model each biological condition separately, they use variational autoencoders to identify cell embeddings, which limits the ability to link genes with specific conditions, only providing a notion of which genes are generally associated with a broad category. Another deep generative method, biolord (Piran et al. 2024), uses latent optimization for disentanglement, but there is even less support for the ability to link genes with conditions, requiring the user to essentially reconstitute predicted counts and perform differential gene expression. In contrast, scParser (Zhao et al. 2024) adopts a matrix factorization framework combined with sparse representation learning to model multiple conditions, but because labels are directly encoded, it makes the unrealistic assumption that all cells with the same condition share the same effect.

Here, we introduce ALPINE (Adaptive Layering of Phenotypic and Integrative Noise Extraction), a novel, flexible approach designed to address the complexities of multi-condition and multi-batch scenarios with improved interpretability (Figure 1). ALPINE builds on the typically unsupervised non-negative matrix factorization (NMF) framework to incorporate supervised, label-guided decomposition of biological conditions and/or technical batches. The use of supervision sets it apart from previous NMF-based integration methods such as LIGER (Welch et al. 2019), and the novel joint supervised-unsupervised representation enables improved identifiability when modeling multiple conditions in a way that methods like scParser (Zhao et al. 2024) does not. This process enables users to directly extract meaningful condition-associated genes, remove batch-associated signatures, and use the unguided components to build a low-dimensional embedding of any remaining variation. In single cell datasets, the unguided components will typically capture the condition- and batch-agnostic cell type variation. The decomposed condition-associated signatures can be simultaneously analyzed for gene associations as well as cell type associations, including between conditions, which can enable efficient exploration of otherwise unwieldy datasets.

**Figure 1.**
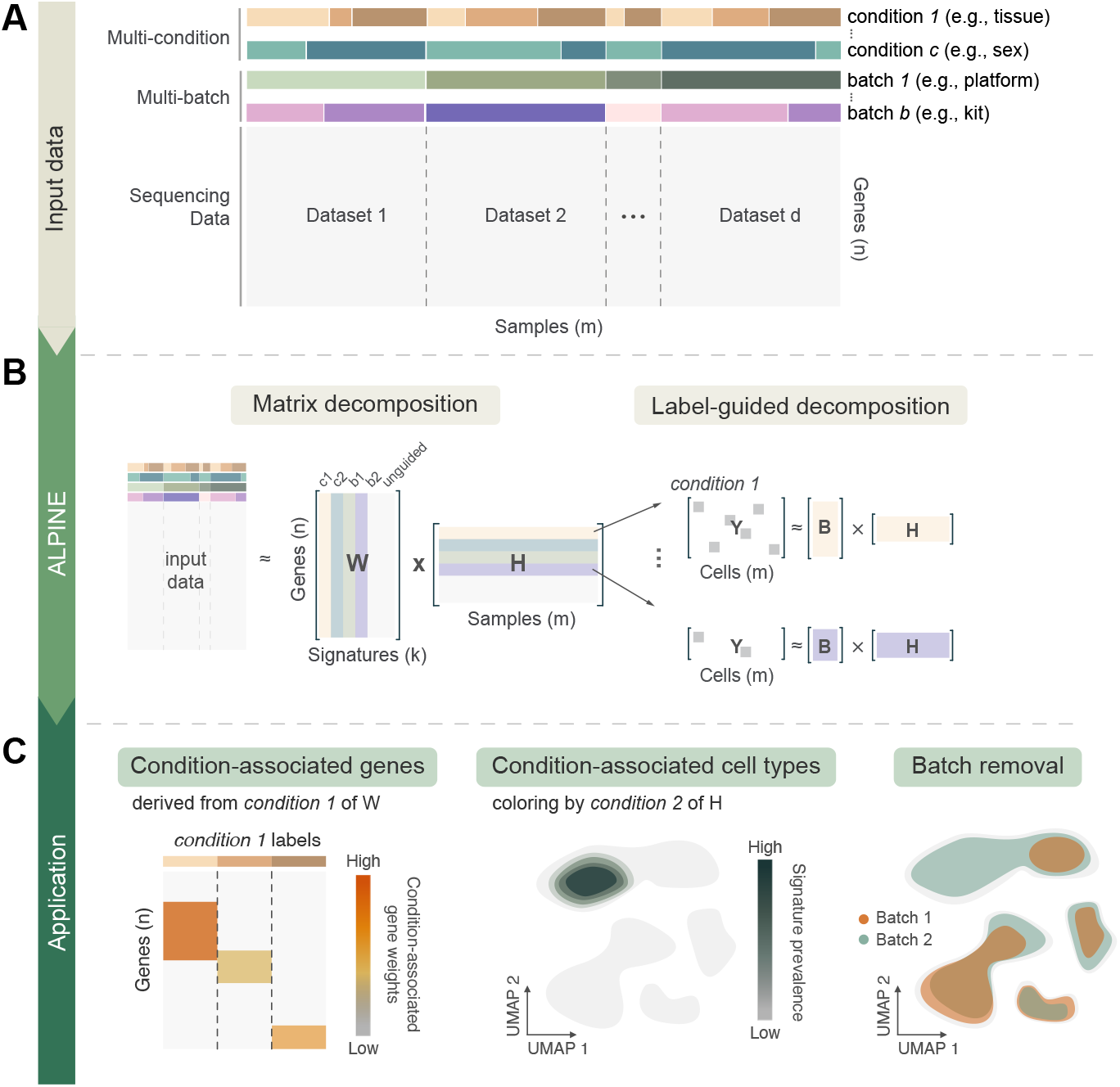
Overview of ALPINE for disentangling the effects of conditions and batches. (A) Example of a dataset with multiple condition and batch effects, creating challenges in cross-study analyses. (B) ALPINE’s workflow incorporates an NMF-based approach with two main components: classic matrix decomposition and label-guided decomposition. The label-guided decomposition enables identification of condition- or batch-associated components. (C) ALPINE extends beyond standard single-cell analyses (e.g., clustering and cell embedding) by uncovering condition-associated genes and cells and addressing batch effects in a principled way, facilitating deeper insights across complex datasets.

## Results

### Overview of the ALPINE framework

ALPINE extends traditional NMF by decomposing scRNA-seq expression data (Figure 1A) into interpretable components that reflect both known covariates and shared biological variation. Specifically, the method separates the data into “guided” components (directly linked to provided labels such as batch or phenotypic condition), as well as an “unguided” component that captures shared residual variation (Figure 1B, Methods). This dual strategy enables ALPINE to not only reconstruct the original data accurately, but also to generate condition-specific gene signatures that are readily interpretable and remove batch effects present in the data (Figure 1C). Guided variables in ALPINE function analogously to categorical predictors in a regression model. Multiple categories are supported, and ALPINE assigns interpretable components to each level. This flexibility allows the framework to capture diverse experimental covariates, such as sex, tissue, or condition, without being restricted to binary labels. To tackle the computational challenges associated with large-scale single-cell datasets, ALPINE employs a mini-batch training strategy to improve computational efficiency and enhance model generalizability (Methods). This approach leverages subsets of cells to update the cell embedding matrix, reducing memory usage and mitigating overfitting. This strategy is particularly valuable when processing large-scale scRNA-seq datasets. Overall, ALPINE’s design provides a powerful framework for disentangling complex biological and technical signals, delivering both high-quality integration and direct interpretability of condition-specific effects.

### ALPINE can disentangle condition information from shared cell information

To test ALPINE’s ability to disentangle multiple biological conditions and batch effects, we simulated datasets of 40,000 cells with one technical batch effect and two binary phenotypic conditions (stimulation: control vs. stimulated; severity: healthy vs. severe), similar to Z Zhang et al. 2024. We then implemented four distinct perturbation scenarios (naive, overlap, two-patterns, cell-specific), which were each generated and evaluated separately (Figure 2A, Supplemental Figure 1, and Methods). The different simulations captured varying effect sizes from the condition and batch effects (Supplemental Figure 2), including scenarios where the batch effect can exceed a weaker condition effect. We compared ALPINE’s performance against the two disentanglement methods we could successfully run (scDisInFact and scParser), as well as two baseline comparisons (raw counts directly or principal components as input). In addition to using ALPINE’s cell embeddings, we can also reconstitute cell counts using only the unguided portions of ALPINE’s *W* and *H* matrices, which are essentially a batch-removed, integrated dataset that can be used for downstream single-cell analyses. We evaluate ALPINE’s reconstructed counts in these simulations for their ability to support accurate cell type clustering and will revisit them with further evaluations in the batch effect removal section below. We note that scDisInFact provides functionality to reconstruct batch-removed counts while scParser does not, so we compare to both scDisInFact embeddings and counts, but embeddings only for scParser.

**Figure 2.**
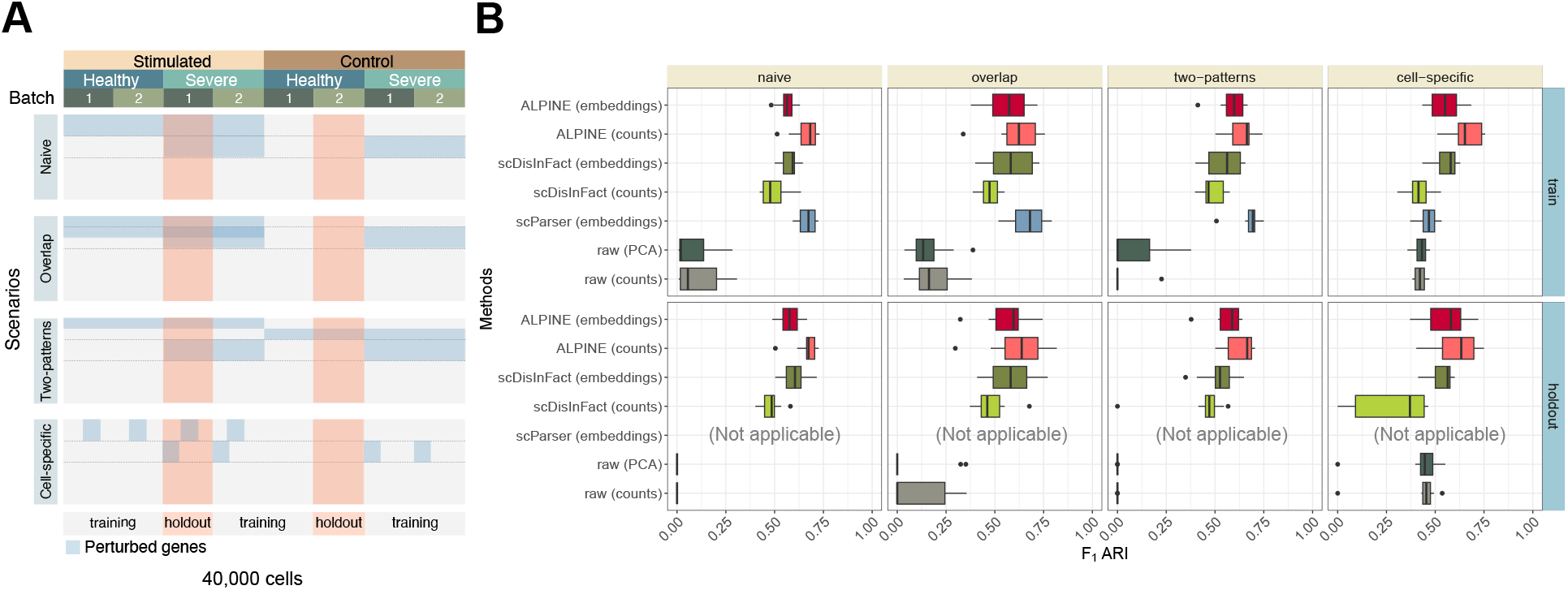
Systematic benchmarking for multi-batch and multi-condition disentanglement using different simulated scenarios. **(A)** Perturbation arrangement of count matrices in the simulated datasets. For each scenario, 10 count matrices are generated, corresponding to batch and 2 condition types (simulation and severity). The four scenarios include: adding signals to one label of each condition with independent perturbations (Naive); shared gene perturbations between conditions (Overlap); signals added to both labels of each condition (two-patterns); and cell-specific signals (Cell-specific). **(B)** Box plots compare the F1 ARI performance based on k-means clustering (with the known number of cell types) of ALPINE (embedding and counts) with two existing methods (scParser and scDisInFact) and two baseline approaches (raw, which uses the confounded counts including both batch and covariate effects directly, and raw (PCA), which uses the top 50 PCs of the raw counts) across training and holdout datasets. ALPINE (both embedding and counts) shows consistently strong performance, especially in more complex scenarios.

In the training data, we find that ALPINE demonstrates strong performance, especially in the more complex, realistic cell-specific scenario. Interestingly, ALPINE’s reconstructed counts often demonstrates even better performance than using the embeddings directly. Specifically, in the naive, overlap, and two-patterns scenarios, where we introduce fold changes to all of the cell types for a given set of genes, the ALPINE reconstructed counts show generally better performance than the scDisInFact embeddings, significantly better performance than the scDisInFact counts, and comparable performance with the scParser embeddings (Figure 2B, Supplemental Tables 1, 2). In the cell-specific scenario, where only a subset of cell types are subjected to the condition effect (while all cell types are affected by the batch effect), the ALPINE embeddings show comparable performance with scDisInFact embeddings and significantly outperform scParser and scDisInFact reconstructed counts, and the ALPINE reconstructed counts significantly outperform all other methods (Figure 2B, Supplemental Tables 1, 2). We note that this cell-specific simulation is a better reflection of real data, as many diseases or treatments affect only a subset of cell types.

To evaluate generalizability, we also held out two sets of cells that constitute entirely unseen combinations of effects (Figure 2). For these comparisons, only scDisInFact and ALPINE can generalize existing trained models to new datasets. The ALPINE embeddings show comparable performance to scDisInFact embeddings and significantly better performance than the scDisInFact counts across all scenarios. Similar to our observations in the training data, the ALPINE reconstructed counts have further improved performance, generally outperforming both the scDisInFact embeddings and scDisInFact counts (Figure 2B, Supplemental Tables 1, 2).

We also evaluated all methods under simulated scenarios with larger amounts of extrinsic variation in gene expression for cells in each cell type, corresponding to higher within cell type variability (Methods). As expected, as the heterogeneity of each cell type increases, all methods suffer and eventually are unable to support effective cell type clustering. That said, the performance trends between methods across different levels of heterogeneity were typically consistent, where ALPINE, especially the reconstructed counts, generally outperforms other methods (Supplemental Figure 3).

In general, we consistently observe that scParser achieves reasonable clustering in simpler cases but begins to lose accuracy in cell-specific perturbations due to its assumption that all cells with the same condition label behave similarly. Only scDisInFact and ALPINE support the ability to reconstruct batch-removed counts, but the reconstructed counts derived from scDisInFact generally underperform in clustering, suggesting that its disentangled VAE architecture does not fully preserve fine-grained cell type distinctions. Both ALPINE’s embeddings and reconstructed counts perform well, with reconstructed counts often outperforming the embeddings, likely because the counts retain richer gene-level variation while filtering out noise.

### ALPINE efficiently identifies condition-associated genes and cells

Using the same four scenarios, we can further assess to what degree the actual condition-associated genes are prioritized by each method. Both ALPINE and scParser successfully identify the true perturbed genes in simpler tasks, such as in the naive, overlap, and two-patterns scenarios (Figure 3A). However, as the complexity of the task approaching more realistic conditions, as in the cell-specific scenario, the AUPRC for scParser begins to decline. This decrease is likely due to scParser’s strong implicit integration of all labels during the matrix factorization stage. In scenarios where not all cells are subject to the condition effect, this approach hampers scParser’s effectiveness. While scDisInFact is capable of detecting some associated genes, it does not capture perturbed genes as effectively as scParser or ALPINE. Importantly, these trends persist even under higher extrinsic variation where cell type clustering breaks down (Supplemental Figure 3). In this more challenging setting, disentanglement methods still retain the ability to detect condition-associated genes, and ALPINE shows the most robust and consistent performance, especially in the cell-specific scenario (Supplemental Figure 4).

**Figure 3.**
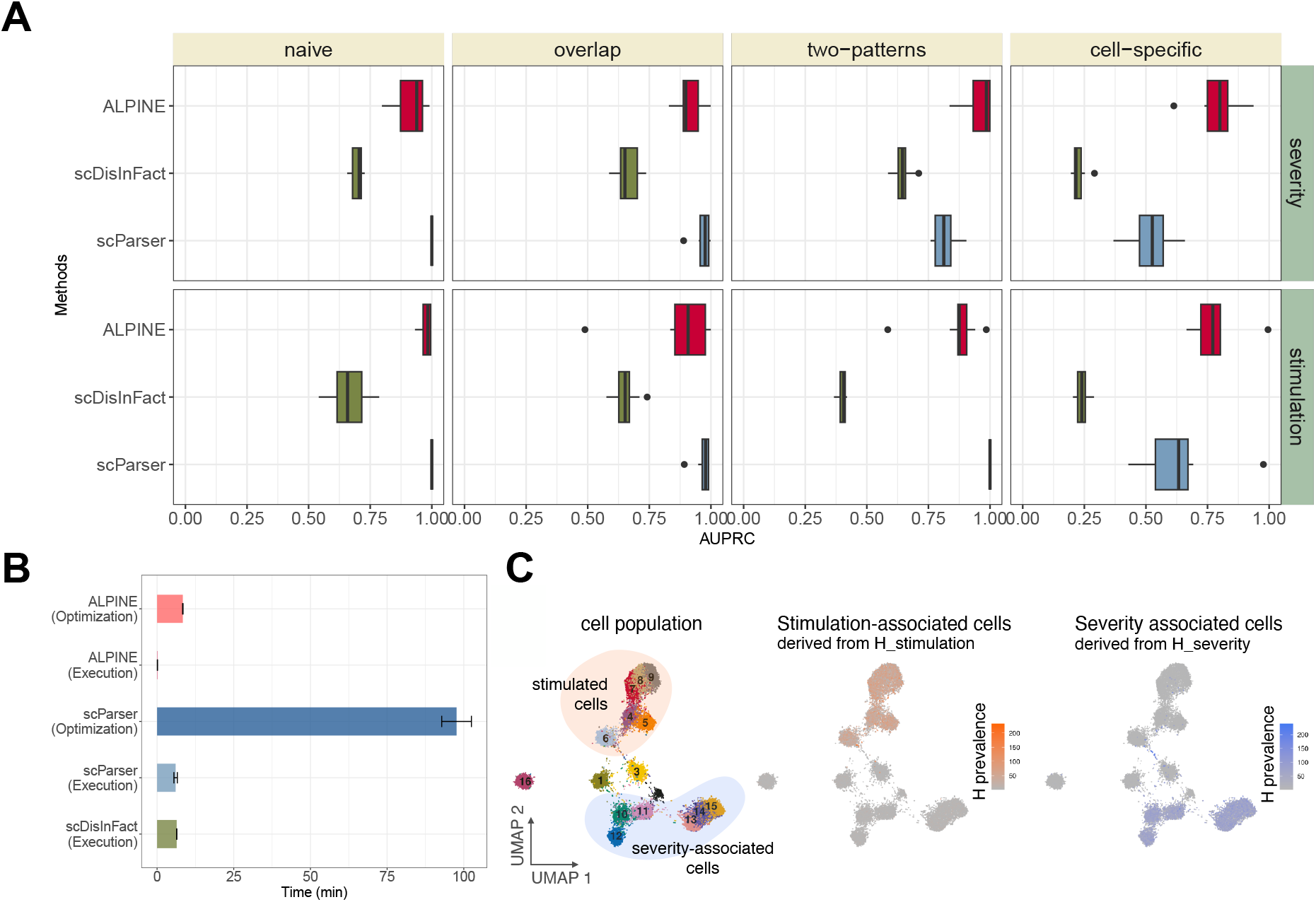
ALPINE can accurately extract condition-associated genes. **(A)** Boxplot of the AUPRC of ALPINE, scDisInfact, and scParser on condition-associated gene detection in four scenarios. **(B)** Algorithm run time (execution) for ALPINE, scParser, scDisInfact and hyperparameter searching (optimization) for ALPINE (which tests 100 hyperparameter sets) and scParser (which tests 5 hyperparameter sets). scDisInfact does not provide an optimization function. **(C)** UMAP plot of cell embeddings with batch and condition effects removed, colored by cell type. Highlighted are cells under two conditions with distinct gene perturbations. ALPINE’s condition embeddings capture condition-associated signatures, accurately identifying stimulation-associated (types 4-9) and severity-associated cell types (types 10-15).

In most single-cell analysis pipelines, differential expression (DE) analysis would be the standard approach for identifying condition-associated genes, so we also compared DE results with the disentanglement methods (Methods). In the simpler naive, overlap, and two-pattern scenarios (Supplemental Figure 5), we unsurprisingly find that it is important to do batch correction before downstream DE analysis, with non batch-corrected comparisons performing the worst. After batch correction, DE analysis can achieve comparable or often better performance than scDisInFact. ALPINE and scParser consistently outperform DE, with scParser matching DE only in one condition. In the more complex cell-specific setting, DE struggles when all cells are included (Supplemental Figure 6A) and the analysis and improves only slightly when restricted to only the perturbed cells (Supplemental Figure 6B). ALPINE outperforms all other approaches in this scenario.

We also note that ALPINE and scDisInFact have similar overall runtimes (Figure 3B), while scParser requires substantial time for its optimization procedure. Once trained, ALPINE has very efficient execution times. We do note that part of the runtime limitations of scParser could be due to the fact that it does not have a GPU implementation, but even directly comparing ALPINE’s CPU implementation, we see that scParser is still many orders of magnitude slower (Supplemental Figure 7).

ALPINE’s decomposition framework enables it to not only identify condition-associated genes, but simultaneously identify cell types associated with specific conditions. We first calculate 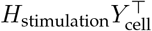 to ascertain the associations between signatures and labels, enabling us to determine which signatures are most relevant to each label. Subsequently, we can directly visualize the relevant component of *H*_stimulation_ to reveal which cell types are predominantly affected by the condition (Figure 3C). The low-dimensional representation visualizes the cell embeddings, with colors indicating prevalence scores derived from the condition-associated signatures. This color-coding reflects the cell types subject to which conditions. Such functionality, to our knowledge, is the first of its kind for methods that learn disentangled representations.

We also use these simulated scenarios to compare training ALPINE using mini-batch versus full-batch updates (Supplemental Figure 8). On the training set, both approaches yield similar F1 ARI scores. However, on unseen holdout sets, mini-batch training typically provides results in better performance, confirming its advantage in generalizing to new data (Supplemental Figure 8A). The only exception occurs in the cell-specific scenario, likely due to the fact that only a tiny proportion of cells are actually affected by the unique conditions, and there may be variable proportions of perturbed cells in each mini-batch. In addition, relative to full-batch training, the mini-batch training strategy typically results in more rapid convergence (Supplemental Figure 8B). Thus, we see that mini-batch training tends to enhance model robustness while reducing computational resource demands.

### ALPINE achieves state-of-the-art performance in batch effect removal and cell type clustering

After demonstrating ALPINE’s power and interpretability in simulation datasets, we further verify ALPINE’s effectiveness using real data. Following the setup in (HTN Tran et al. 2020), we use three datasets: (1) aligned cell types from two batches. The data is from 15,476 human peripheral blood mononuclear cells, and they are from two different sequencing platforms (10x 3’ and 10x 5’) (GX Zheng et al. 2017); (2) aligned cell types with multiple batches. The data consists of five studies of human pancreatic cells from four different technologies with a total of 14,767 cells (Baron et al. 2016; Segerstolpe et al. 2016; Muraro et al. 2016; YJ Wang et al. 2016; Xin et al. 2016); (3) non-identical cell types, with mouse retina data from two studies, including 62,851 cells (Shekhar et al. 2016; Macosko et al. 2015). Besides scParser and scDisInFact, we include seven methods (ComBat, Harmony, Liger, MNN, Seurat, Scanorama, and scVI) that are designed to perform batch removal and cell type clustering, even though they do not extract interpretable condition features. As a further baseline, we also include using the PCA using the top 20 PCs to assess how severe the original batch effects are in the input dataset.

In all three scenarios, the ALPINE embedding shows better or comparable performance in batch removal and cell type clustering compared to state-of-the-art batch removal methods (Figure 4, Supplemental Figure 9). The other two condition-disentanglement methods, scDisInFact and scParser, generally perform poorly. We suspect that the poor performance of scParser stems from its assumption that all cells in the same batch undergo identical transformations and its poor scalability, which required us to limit its optimization time to 12 hours. For scDisInFact, it is possible that the lack of clear stopping criteria and hyperparameter optimization, as well as the requirement that a condition label be given even when there is no additional condition information, may be related to its suboptimal performance. The ALPINE embedding consistently has the best ARI scores and comparable NMI scores (scenario 1: ALPINE embedding 0.798 vs best: 0.820; scenario 2: ALPINE embedding 0.769 vs best: 0.816; scenario 3: ALPINE embedding is the best) to the next best method. The discrepancies between ARI and NMI here are primarily a result of cluster over-splitting using Leiden, with NMI at times being higher even when ARI is lower. ALPINE’s reconstructed counts achieve the highest-performing ARI and NMI in all three scenarios, except ARI for scenario 3 (ALPINE counts 0.410 vs best: 0.411). Comparing cell type ASW and batch ASW scores, both ALPINE embedding and reconstructed counts also demonstrate strong performance compared to existing batch correction methods (Supplemental Figure 9).

**Figure 4.**
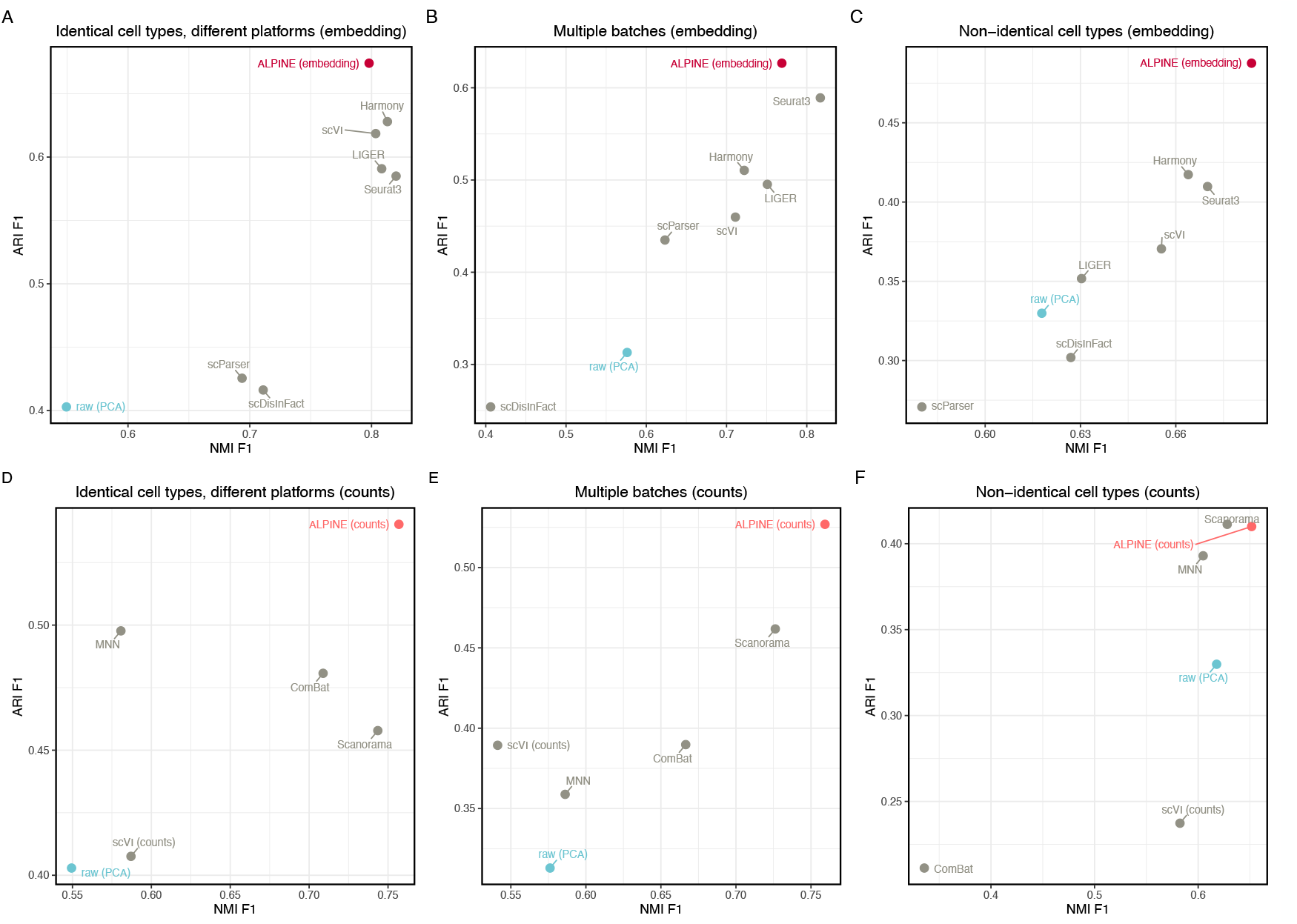
Comparative study of ALPINE against other methods for batch effect removal and cell type clustering. Adjusted Rand Index (ARI) and Normalized Mutual Information (NMI), of ALPINE versus scDisInFact (Z Zhang et al. 2024), scParser (Zhao et al. 2024), Seurat3 (Stuart et al. 2019), Harmony (Korsunsky et al. 2019), LIGER (J Liu et al. 2020), scVI (Lopez et al. 2018), Scanorama (Hie et al. 2019), MNN (Haghverdi et al. 2018), and ComBat (Johnson et al. 2007), in batch effect removal using three real datasets based on cell type clustering with Leiden (using default resolution=1). Methods producing low-dimensional embeddings (scDisInFact, scParser, Seurat3, Harmony, LIGER, scVI) are compared with ALPINE embeddings in **(A)**-**(C)**, while methods reconstructing counts (Scanorama, MNN, ComBat, and scVI (counts)) are compared with ALPINE-reconstructed counts in **(D)**-**(F). (A)&(D)**. Human peripheral blood monouclear cell datasets with two batches and matched cell types. **(B)&(E)**. Pancreatic cells dataset with five batches and matched cell types. **(C)&(F)**. Mouse retina data with two batches and non-identical cell types.

To directly assess how each method’s corrected counts accurately recover underlying expression patterns, we extract ground-truth counts (unaffected by batch or covariates) from our earlier simulated scenarios. We note that 3 of these methods (ComBat, MNN, and Scanorama) are not designed to generalize to unseen datasets, so we can only evaluate them on training dataset performance. In the training set, ComBat and MNN have the best performance, closely followed by ALPINE, and ALPINE achieves the lowest error in reconstructing counts for both the validation and holdout sets (Supplemental Figure 10).

### ALPINE achieves robust disentanglement and cross-species integration to enable condition analyses

We next apply a more end-to-end analysis of ALPINE on two large-scale real datasets. The first, a brain cancer dataset, provides a benchmark for disentangling patient, sex, and cancer-type covariates. The second, an adipose tissue dataset combining human and mouse samples across assays and tissues, demonstrates ALPINE’s ability to generalize to cross-species integration.

The brain cancer dataset (Abdelfattah et al. 2022) consists of cells from 18 patients spanning both sexes and different cancer types (glioblastoma, recurrent glioblastoma, astrocytoma, and oligodendroglioma). All three methods demonstrate reasonably strong performance in eliminating batch and covariate effects (Supplemental Table 3), but scParser exhibits poorer cell type clustering performance (ARI: 0.171) compared to scDisInFact (ARI: 0.352), the ALPINE embeddings (ARI: 0.413), and the ALPINE reconstructed counts (ARI: 0.359, Supplemental Table 4). For comparisons of condition-associated genes, we note that ALPINE and scParser can identify genes with individual labels within the covariate, while scDisInFact can only identify genes associated with the broader condition. This dataset highlights this difference, as ALPINE and scParser can provide gene weightings associated with each of the four cancer types, whereas scDisInFact can only provide a single list of genes associated with the overall cancer type covariate. When comparing the resulting condition-associated genes (Supplemental Figure 11), we find that there are typically a small set of genes that are identified by all 3 methods and more genes that are distinct for each method. The largest intersections are typically between ALPINE and scParser, with typically fewer between either method with scDisInFact. This discrepancy may be due to structural differences in the methods, as both ALPINE and scParser use matrix factorization, but it could also be a direct result of the fact that the genes identified by scDisInFact are, by design, a mixture of genes associated with the individual underlying labels.

We further demonstrate ALPINE’s versatility through an additional case study by applying it to a white adipose tissue dataset containing diverse batches (assays: Drop-seq, 10x 3’ v3) and conditions (organisms: Homo sapiens, Mus musculus; sex: male, female; depot tissues: subcutaneous adipose tissue, periovarian fat pad, omental fat pad, inguinal fat pad, and epididymal fat pad). The original study was only able to analyze each species separately, but here, we demonstrate that ALPINE is capable of identifying unifying cell embeddings across species, and the interpretability of its components further enables a more comprehensive comparison of cellular characteristics across species. Before further downstream analysis, we find that applying ALPINE directly with associated conditions effectively removes batch effects (Figure 5A), yielding well-mixed cell populations across species, with the notable exception of adipocytes, which remain distinct.

**Figure 5.**
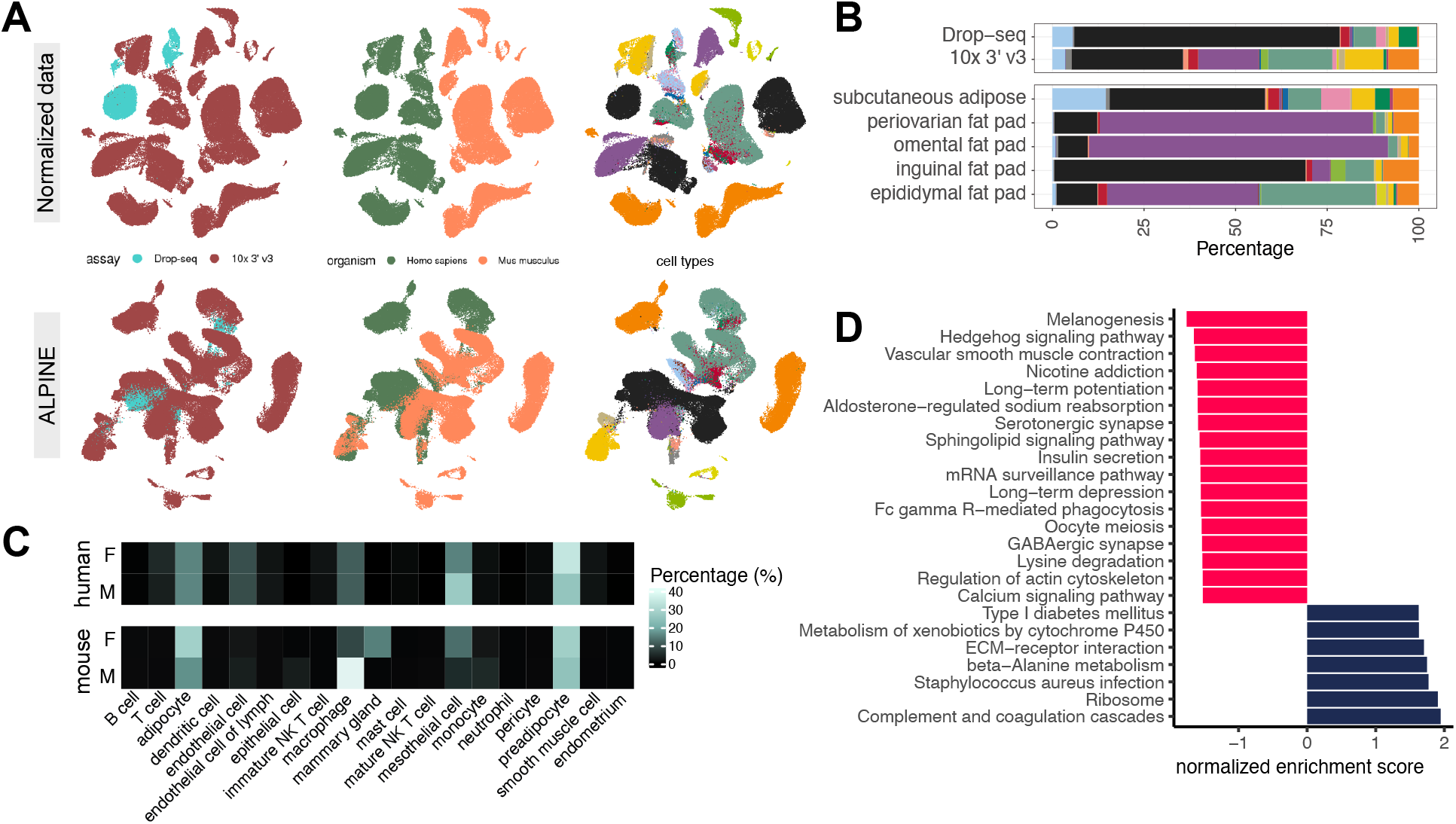
Cross-species analysis in adipose tissue with multi-condition analysis. **(A)** The original dataset displays multiple batch and condition effects, with colors indicating various categories. Post-ALPINE application, batch effects from sequencing technologies are removed, resulting in aligned clusters of similar cell types across species, although adipocytes remain distinct due to biological differences. **(B)** Condition embeddings reveal associations between cell types and assay/tissue labels, accurately capturing expected cell types. Notably, Drop-seq fails to identify fragile adipocytes, which are detectable only via sNuc-Seq. Tissue embeddings further represent accurate cell-type abundances across conditions. The stacked bar plot is color-coded by cell types, consistent with (A). **(C)** Analysis of sex embeddings, separated by species, indicates that human cell compositions are more similar between males and females, while mouse samples exhibit significant differences, underscoring the identified sex-related variations. **(D)** The bar plots display normalized enrichment scores for KEGG terms from the GSEA analysis, comparing human and mouse adipocytes using gene data from *W*_organism_. Positive values (navy bars) indicate higher enrichment in humans, while negative values (red bars) denote higher scores in mice.

Interestingly, we find that ALPINE’s cell embeddings accurately reflect a low abundance of adipocytes in Drop-seq versus a higher representation in sNuc-Seq, recapitulating the established platform-specific difficulty of capturing adipocytes (Figure 5B). We also find distinct cell composition variations across fat depots, notably a higher macrophage proportion in the male mouse epididymal fat pad, which prompted further exploration of sex-related differences. Within the sex-specific embeddings, ALPINE correctly captured increased macrophages in male mice, as well as mammary gland epithelial cells unique to female mice (Figure 5C). In human samples, we see a more even distribution of immune cell types compared to mice, highlighting ALPINE’s ability to capture detailed, biologically relevant signals across species.

An important unanswered question is what biological factors drive the distinct differences between mouse and human adipocytes. In the original study, this comparison was infeasible because the mouse and human cells were not embedded in a shared space. We calculated the difference between human and mouse adipocyte gene signatures and applied GSEA to reveal functional distinctions (Figure 5D), identifying several adipose-related signals, including melanogenesis (Randhawa et al. 2009), which has been reported in human adipose tissue, where various melanogenesis-related genes contribute to the presence of melanin. Additionally, we find the Hedgehog signaling pathway (Fontaine et al. 2008), known for its dysregulation during adipocyte differentiation. Other enriched terms, such as the sphingolipid signaling pathway (Łukaszuk et al. 2024), further demonstrate the biological changes associated with adipose tissue. These findings underscore the utility of our methods in uncovering the underlying biological functions linked to specific conditions.

We also compare ALPINE’s findings with those of scDisInFact on the same dataset (Supplemental Figure 12). Unfortunately, we were unable to obtain results for this dataset from scParser (see Methods). Initially, we analyze the integration measures for cells, covariates, and batch, observing comparable outcomes between ALPINE and scDisInFact except for ARI_cell_ (Supplemental Tables 5 and 6). scDisInFact’s conditional variational autoencoder-based model limits its ability to derive covariate-associated weights for each sample, so we are unable to construct a covariate-associated cell embedding space to enable the composition analysis described above.

We find that a further limitation of scDisInFact for interpretability is an inability to associate gene scores within individual labels. For the cross-species analysis, this means that we can only obtain a single organism-associated gene list, as opposed to distinct human- and mouse-associated genes. Comparing the individual genes identified (see Methods), ALPINE identifies 383 and 226 significantly associated genes for mouse and human, respectively, while scDisInFact finds 112 general organism-associated genes. There are only 5 genes found in common between scDisInFact’s genes with ALPINE (all with the human-associated signatures). Interestingly, among these are four mitochondrial genes (MT-CO2, MT-CO3, MT-ATP6, MT-ND3) that are the top-ranked genes by both methods. Although no cross-species comparison has been conducted on these genes, one study indicates that these mitochondrial genes play a crucial role in cold-induced mitochondrial biogenesis in white adipose tissue (Ito et al. 2024). Another study also identified MT-CO2, MT-ND3, and MT-ATP6 as being correlated with BMI in their meta-analysis, potentially linking them to participants in the study with high BMI (Kraja et al. 2019). While this shared identification is exciting, unfortunately, the organism-associated gene scores from scDisInFact do not yield significant GO enrichment using GSEA (FDR ≤ 0.1) for further functional comparisons with ALPINE. In general, these results illustrate that ALPINE not only achieves effective cross-species integration, but also delivers enhanced interpretability by explicitly delineating condition-specific signals.

### Discussion

In this study, we demonstrate that ALPINE is capable of simultaneously addressing multiple challenges in single cell data analysis, including disentangling complex batch and condition effects and finding condition-associated effects on genes, cells, and inter-condition interactions. Compared to existing disentanglement methods, ALPINE provides significant advantages in integration performance, interpretability, scalability.

Interestingly, across both simulations and real datasets, our analyses reveal a tradeoff among existing disentanglement methods. Some approaches excel at cell clustering but are limited in resolving condition-associated genes, while others capture gene-level associations at the cost of clustering robustness. Furthermore, we observe that beyond practical limitations such as runtime and usability, scParser can struggle with cell-type specific perturbations and does not provide the ability to generate reconstructed counts. scDisInFact provides that feature, but shows uneven performance between their embeddings and reconstructed counts. It also can only identify covariate-level rather than label-specific gene associations, limiting interpretability. Meanwhile, ALPINE uniquely balances these tasks and performs well across both of its outputs, even under heterogeneous perturbations or high extrinsic variability. Both its embeddings and reconstructed counts support accurate clustering, and its embeddings recover label-specific gene signatures that enable more precise biological interpretation. The consistency in ALPINE stems from its unique joint supervised-unsupervised design, which combines supervised guidance with interpretable factorization to both preserve cell-level structure and condition-specific gene signals.

While ALPINE demonstrates strengths in disentangling batch and condition effects and capturing both general and cell-specific signals, it may face limitations in scenarios with complex, non-additive, or non-linear interactions among conditions, as these may not be fully captured by its underlying NMF framework. Additionally, handling highly variable cell populations or very subtle condition effects may reduce interpretability. Future work could focus on extending ALPINE’s framework to capture non-linear interactions. Furthermore, it is also possible for biological variables of interest to exhibit more complex, interesting structure, such as continuous values or nested hierarchical structure. ALPINE can be naturally extended by considering different loss terms for different types of guided variables to handle that additional complexity. One practical limitation of ALPINE is that it currently relies on users to identify and combine fully confounded variables during preprocessing since their separate effects are not statistically identifiable. Future versions could incorporate model or feature selection strategies, which would enable the framework to automatically identify the most informative combinations of covariates. Finally, there are also opportunities to further refine ALPINE’s built-in hyperparameter optimization procedure, for example by incorporating adaptive strategies for clustering, which could improve robustness across diverse datasets.

In summary, ALPINE’s flexible and interpretable framework enables it to meet a wide range of analytical needs, providing robust performance across datasets with varying batch and condition complexities. As a scalable and versatile tool, ALPINE has significant potential for advancing condition-specific studies in diverse single-cell applications, offering researchers a powerful method to uncover subtle biological patterns and interrelationships within complex, heterogeneous data.

## Methods

### Overview

ALPINE aims to decompose a heterogeneous single cell transcriptomics dataset into interpretable low dimensional components. Specifically, a given single cell sample is often associated with different technical batch effects and phenotypic conditions, which we term ‘guided variables.’ ALPINE jointly optimizes for the decomposition of an input dataset into guided and unguided components and includes a built-in hyperparameter tuning framework to efficiently select hyperparameters. A key principle of the design of ALPINE is that there are no constraints on the relationship between guided variables, but the unguided components should capture shared biological signals across conditions and not signals that are represented in the guided variables. As illustrated in Supplemental Figure 13, the guided embeddings each align with their corresponding covariates, while the unguided embedding captures cell-type variation not attributable to batch or condition effects.

### Label-guided matrix decomposition

Given a log-transformed single-cell RNA expression matrix *X* ∈ ℝ ^*m×n*^, where *m* is the number of genes and *n* is the number of cells, ALPINE decomposes *X* into a non-negative gene feature matrix *W* ∈ ℝ^*m×k*^ and a cell embedding matrix *H* ∈ ℝ^*k×n*^, where *k* represents the total number of latent components.

For a given guided variable, we define the corresponding submatrices as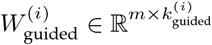 and 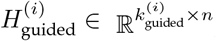, ensuring that the total number of components satisfies: 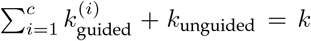, where *c* denotes the number of user-specified guided variables.

Each component is uniquely associated with either a guided variable or an unguided shared variation. In scenarios where certain guided variables are entirely confounded with each other (e.g., each condition is exclusively sequenced in a distinct batch), the corresponding guided variables become mathematically collinear (analogous to including perfectly correlated predictors in a regression model). Practically, users can identify such cases during data exploration by inspecting the overlap between covariates (e.g., crosstabulation of labels) or by checking for perfect correlations in the design matrix. Domain knowledge of the study design is helpful here, as collinearity often arises from experimental choices such as sequencing each condition in a separate batch. Under such situations, the model cannot uniquely partition the variation between the different guided variables, leading to an identifiability problem. To mitigate this, we recommend combining the confounded guided variables into a single composite variable (e.g., merging separate “female” and “healthy” labels into a combined “female + healthy” label) that captures their joint effect. This approach allows ALPINE to extract the shared biological signal while reflecting limits imposed by the experimental design.

Overall, the matrices *W* and *H* are formed by concatenating their respective submatrices:

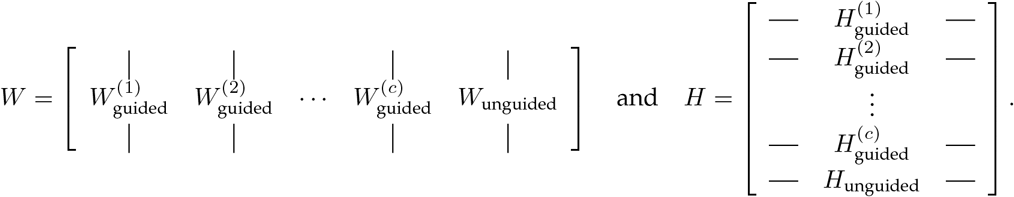

To effectively disentangle the signals of different guided variables and shared cell information, ALPINE uses a binary guided indicator matrix, 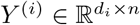 for each guided variable *i*, where *d*_*i*_ is the number of classes for that variable. The primary goal is to ensure that each cell embedding 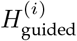 encapsulates all relevant information for 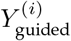. To achieve this, 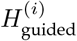 is used to reconstruct *Y* ^(*i*)^ through multiplication with a learned transformation matrix 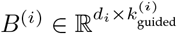. The training objective function is thus a combination between the unsupervised reconstruction loss and supervised prediction loss:

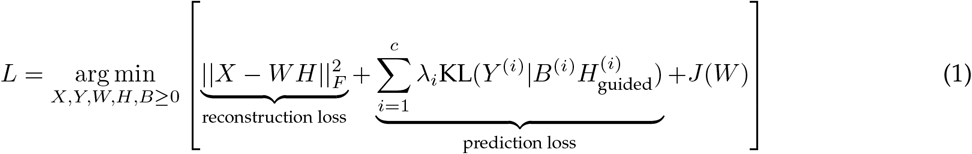

Here, || *·* ||_*F*_ represents the Frobenius norm that quantifies the reconstruction error between the matrix *X* and its approximation *WH*, and *λ*_*i*_ is a hyperparameter that balances reconstruction and prediction loss. KL represents the generalized Kullback-Leibler divergence (Lee and Seung 2000), which is particularly suitable for fitting the model to the binary condition matrix *Y* ^(*i*))^, as it quantifies the discrepancy between the predicted probabilities and the actual binary outcomes. We also include a regularization term, *J* (*W*), to encourage the model to learn more unique and generalizable signatures by incorporating elastic net regularization and orthogonality in the *W* matrix (Lin and Boutros 2020).

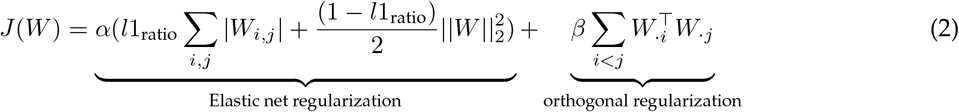

where hyperparameter *α* represents the weight for the Elastic net regularization and *l*1_ratio_ is the weight balancing LASSO and Ridge regularization. The final term, weighted by hyperparameter *β*, computes the sum of the product of two signatures, where lower values indicate lower similarity, promoting orthogonality.

The reconstruction loss optimizes for preservation of key biological information while the supervised prediction loss helps disentangle signals from different guided variables into the designated components. For a given guided variable, examining the sub-matrix 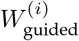 shows the contribution of individual genes to a low-dimension representation of the variable, while 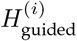 highlights the relative prevalence of the corresponding gene signatures in each cell. The remaining components *W*_unguided_ and *H*_unguided_ capture clean biological information that is consistent across different batches or conditions, which can be used for clustering cells or assigning cell types.

We derive the multiplicative updates for *W, B*, and *H* based on Equation (1) (full derivation details in Supplemental Methods):

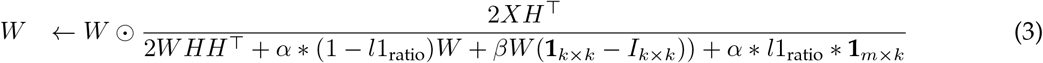

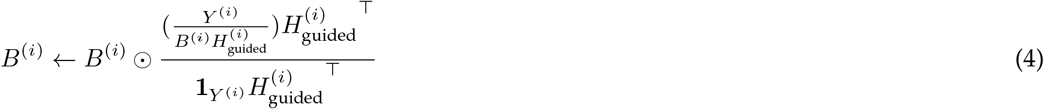

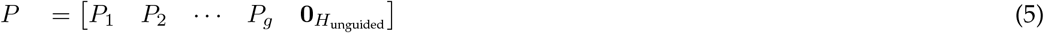

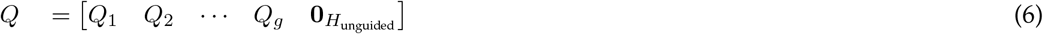

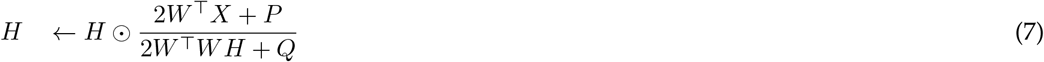

In the multiplicative update of *W*, **1** and *I* denote the ones matrix and the diagonal matrix, respectively. Here, 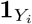 represents a matrix of ones with the same shape as *Y* ^(*i*)^ and 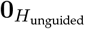 represents a matrix of zeros with the same shape as *H*_unguided_. The supervised learning updates *P*_*i*_ and *Q*_*i*_ are formulated as 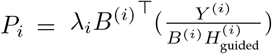 and 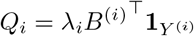, where *P* and *Q* represent combinations of guiding adjustments derived from the prediction loss associated with the guided variables. Because there are no labels for the unguided components, we use zeros to fill the *P* and *Q* submatrices. During the updating process, *P* and *Q* help regularize *H* with guided label information, pushing the components to be specific to each guided variable. The final output of ALPINE are the final lower-dimensional representations *W, H*, and *B* that compartmentalize shared biological information and different phenotypic conditions.

### Generalization to unseen datasets

Given an unseen dataset, ALPINE can use pre-trained *W* and *B* matrices from a previous integration to generalize to new single-cell data. For the new dataset, we use the new counts (*X*_new_) and iteratively update the new dataset cell embedding *H*_new_ based on Equation 7 (with *P* and *Q* being matrices of 0s). Note that while a corresponding indicator matrix (*Y*_new_) can optionally be used to update *B*^(*i*)^, *P*, and *Q* using Equations 4-6, we do not recommend doing so to avoid potential overfitting.

### Mini-batch training

A common limitation of existing matrix factorization approaches for single-cell analysis is the extensive computational resources required. As single-cell datasets grow, processing the entire expression matrix becomes increasingly time-consuming and memory-intensive. To address this, ALPINE implements minibatch training, which updates only a subset of cells at a time, thereby conserving training resources. We choose to deploy the mini-batch strategy only on the cells space and not the feature space, since by the nature of scRNA-seq data, the number of features (i.e., genes ∼20,000) is fixed and capturing gene-gene relationships is critical, whereas the number of cells can be extremely large (ranging from tens of thousands to millions). For reference, using full-batch training on a dataset with 1 million cells would use more than 40 GB of memory, making it infeasible for many GPUs, while mini-batch training can be easily adapted for available hardware.

ALPINE supports two sampling strategies: (i) random sampling, which shuffles the cells and ensures every cell is visited once per epoch; and (ii) weighted sampling, which addresses imbalances in guided variables.

In weighted sampling, for each cell, ALPINE calculates a weight based on the overall class representation (concatenating all guided variables), specifically weight 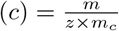, where *m* is the total number of cells in the dataset, *z* is the total number of class groups, and *m*_*c*_ is the number of cells annotated to class *c*. Within each mini-batch, cells are then randomly sampled with replacement based on these weights. This increases the representation of rare labels during training, though it does not guarantee that every cell is seen in each epoch. *W, H*, and *B* are updated following Equations (3-7), but for *H*, in each step, only the subset of cells that correspond to the selected batch is updated. By updating the cell embedding matrix *H* using only a subset of cells in each mini-batch, we reduce the computational burden and memory footprint, also mitigating overfitting and reducing the potential bias introduced by label imbalance in the guided variables. This sampling strategy is analogous to stochastic gradient descent, where updating parameters using subsets of data helps avoid local minima and improves convergence speed.

### Hyperparameter selection

The main hyperparameters of ALPINE include the number of components for both guided 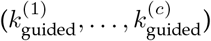 and unguided (*k*_unguided_) variables, along with *λ*_*i*_, *α, β*, and *l*1_ratio_ from Equations (1-2). An additional hyperparameter is the number of epochs, or iterations of multiplicative updates, we apply to optimize ALPINE. To enhance user experience and reduce manual tuning, we designed an optimization procedure to select hyperparameters, where only ranges of values need to be provided for each hyperparameter (we have default ranges as described below). For tuning the number of components, only the range for the total number of components, *k*, needs to be provided, and the optimization process automatically allocates components between the guided and unguided parts.

Internally, given *c* representing the number of user-specified guided variables, we consider the total number of components, *k*, as being distributed among *c* +1”parts,” i.e., the *c* guided variables and 1 unguided part. To tune for the specific number of components in each part, every component, including the unguided one, is assigned a proportion: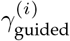 for guided and *γ*_unguided_ for unguided, with the constraint that 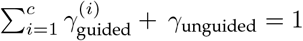. Furthermore, to ensure a sufficient number of unguided components even as the number of user-provided guided variables increase, we enforce 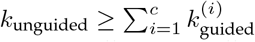, or equivalently, 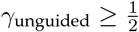.

The optimization algorithm dynamically tunes the component allocation following these constraints, making the process more efficient and user-friendly.

Another optional hyperparameter is the maximum number of iterations that the multiplicative updates in ALPINE should run. If not specified, ALPINE performs a warm-up run with up to 200 iterations and determines the maximum number of epochs automatically by selecting the elbow of a polynomial curve fit on the reconstruction loss curve using the Kneed package (Satopaa et al. 2011). Selecting the elbow on the polynomial curve rather than the direct reconstruction values helps with robustness of elbow detection, since it smooths potential variability from mini-batching. We use reconstruction loss for this procedure because the prediction losses for batch or condition labels typically plateau earlier than the reconstruction loss, reflecting the lower-dimensional nature of these tasks compared to full data reconstruction. As a result, monitoring prediction loss alone risks premature stopping before sufficient reconstruction occurs (example in Supplemental Figure 14).

Since ALPINE is designed to ensure that the unguided components should capture shared biological signals across conditions distinct from those specific to the guided variables, to optimize the model’s hyperparameters, we assess whether *H*_unguided_ has successfully excluded residual guided information. Specifically, clusters (*C*_unguided_) are derived from the *H*_unguided_ matrix using the Leiden algorithm (using the igraph implementation and default resolution of 1). We then compute both the adjusted rand index (ARI) and homogenenity score (HS) between these clusters and the guided variable labels, averaging the scores across all guided variables. Since the goal is to minimize the influence of guided labels on *H*_unguided_, we seek to minimize both ARI and HS. Minimizing ARI ensures that the clustering of *H*_unguided_ does not reflect the guided labels, while minimizing the HS ensures that clusters are well-mixed with respect to these labels. Together, the loss function being optimized is:

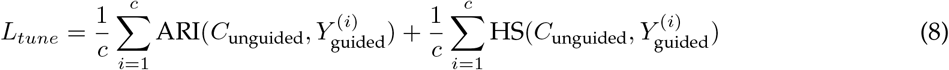

These hyperparameters are tuned with Bayesian optimization using Tree of Parzen Estimators (Bergstra et al. 2013) from their corresponding search spaces with 100 calls for the set of hyperparameters that minimizes *L*_*tune*_. We use [10, 100] as our search space for *k*, the total number of components. *λ*_*i*_ can depend on the complexity of the corresponding guiding variable *i*, so we use a large range [1, 10,000]. The default ranges for *α, β*, and *l*1_ratio_ are [0, 100], [0, 1], and [0, 1], respectively.

To gain a better understanding of ALPINE’s sensitivity to different hyperparameters, we explore performance variability across each of the major hyperparameters using three real datasets. Using the optimized hyperparameter set from our optimizer, we vary one hyperparameter at a time while keeping the others fixed. We find that performance the optimizer consistently finds strong sets of hyperparameters, and the hyperparameters that have the largest effect on performance are the total number of components, the guided (batch) component ratio, as well as the contribution of prediction loss (*λ*) (Supplemental Figure 15). The optimal hyperparameter values does vary across different datasets, highlighting the benefit of our adaptive optimizer.

To further enhance the generalizability of ALPINE, we added a cross-validation option to the optimization process. The training data is randomly split into k folds. For each fold, we calculate *L*_*tune*_ as described previously, selecting the hyperparameter set with the smallest average *L*_*tune*_. This cross-validation step helps ALPINE generalize better to unseen data. We used 3-fold cross-validation for the simulation dataset. For the benchmark and adipose tissue datasets, we did not apply cross-validation, as those tasks do not require applying trained ALPINE models to unseen data.

### Data simulation for conditional gene detection and batch removal

To demonstrate that ALPINE effectively separates diverse covariate influences, including both phenotypic conditions and batch effects, we used the Symsim tool (X Zhang et al. 2019) to simulate 40,000 cells with 500 genes generated from a phylogenetic tree with 16 distinct cell types as ground truth, including a simulated “rare” cell type with ∼200 cells and all other cell types having similar sizes (∼1400-1600 cells). Batch effects were introduced using Symsim with default parameters (batch effect size=1). We also introduced two condition covariates: “stimulation” (comprising control and stimulation) and “severity” (encompassing healthy and severe states), yielding four condition combinations (Figure 2A). We randomly assigned each cell a “stimulation” and “severity” label, giving each covariate a 50-50 chance for their respective labels.

Perturbations were simulated by upregulating 100 randomly selected genes per condition (200 total per scenario) to an average two-fold increase in expression, with slight variability around the fold-change. We considered 4 perturbation scenarios: naive, overlap, two-patterns, and cell-specific. In the naive scenario, perturbations to gene expression are applied uniformly on the randomly selected genes. The overlap scenario models cases where both conditions influence a subset of the same genes. In the two-patterns scenario, the stimulated and control conditions perturb different genes, while the severe condition perturbs the same genes. Finally, the cell-specific scenario is the most complex, as it involves selective perturbing of cell types, thus simulating condition effects within only specific cell populations.

We also varied the extrinsic variability of cell expression for the naive and cell-specific scenarios, which equates to changing the *σ* hyperparameter in Symsim. Under the hood, Symsim uses a simulated phylogenetic tree to simulate single cell expression. *σ* controls the standard deviation of the phylogenetic tree position for each cell. Higher *σ* leads to greater heterogeneity, which can obfuscate cell type differences. The default value for *σ* is 0.4, and we have extended the values to 0.6, 0.8, and 1.0.

### Identifying genes associated with specific covariates and affected cell types

Recomposing the various decomposed matrices allows for interpretable analysis of resulting associations. To systematically extract genes associated with covariates, we consider a general covariate with corresponding label matrix 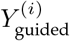 and signature matrix 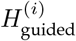. We quantify the association between latent signatures and covariate-related labels by computing

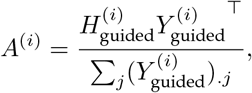

where the denominator normalizes by the number of samples associated with each label, and *j* indexes the labels. The resulting matrix *A*^(*i*)^ captures the strength of association between signatures and covariate labels.

We can further utilize the association matrix *A* to compute the gene signatures associated with specific labels. By multiplying *W* with *A*^(*i*)^, where the result has dimensions corresponding to genes and labels, we obtain our covariate-associated gene signatures matrix. Each column in this matrix represents a signature associated with a particular label.

Additionally, the 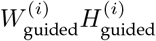 matrix offers insights into the relationships between genes and corresponding cells, enabling users to explore the effects of various conditions on cells. Alternatively, users may investigate the 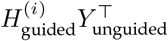 matrix to uncover associations between guided variables and labels, providing a comprehensive understanding of condition-associated signatures and their impact across cell populations.

### Performance evaluation metrics for model assessment and identifying condition-associated genes

To provide objective comparisons of the performance of different tools, we use the adjusted rand index (ARI), normalized mutual information (NMI), and Average Silhouette Width (ASW) to evaluate how well cells are blended across different conditions. Ideally, for cell clustering, ARI and NMI scores close to 1 indicate that each cluster contains only one type of cell, reflecting pure clusters. However, for assessing guided variables, the goal is to have cells from different conditions and batches be well-blended, so to calculate a single performance metric, we report 1 − ARI_guided_, where a value closer to 1 represents better mixing of conditions or batches. We use the transformation from scIB (Luecken et al. 2022) to process the ASW score, mapping both batch and cell type ASW to a 0-1 range, with higher values indicating better performance. To prevent majority cell types from dominating the results, the final batch ASW score is weighted by number of cells in a cell type.

For easier representation when there are situations with multiple batches and conditions, we adopted the F1 score equation from (HTN Tran et al. 2020) to evaluate the performance for removing batch effects while retaining cell type information:

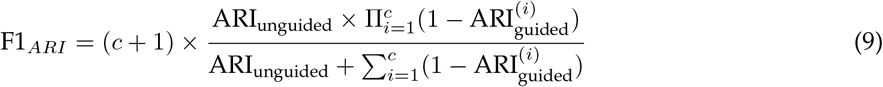

where *c* is the number of guided variables, which includes both batch and condition variables. The F1 score for the normalized mutual information (NMI) follows the same formula as the ARI, with ARI values simply replaced by their corresponding NMI values.

Both ARI and NMI require evaluation on clustering results. To address this, we employed two strategies: (1) k-means clustering by specifying the known number of cell types and then using the k-means results to compute ARI and NMI with scIB (Luecken et al. 2022), or (2) Leiden clustering with a default resolution of 1. For the comparisons using counts rather than embeddings, we clustered the counts directly and then used the same metrics for evaluations. The clustering approach used for each analysis is specified in the corresponding legend.

To evaluate each tool’s ability to identify true condition-associated genes, we calculated the Area Under the Precision-Recall Curve (AUPRC), comparing each tool’s top-weighted genes against ground truth perturbations. For scDisInFact, we directly used the weights provided by the tool. In order to run scParser on the simulation results without crashing, we encoded stimulation and severity into a single composite label (e.g., control + severe). To score recovery of genes for a given condition label (e.g., ‘severe’), we computed the AUPRC between the relevant set of composite signature’s gene weights and the ground-truth perturbed gene set for that level, and reported the highest AUPRC across signatures as scParser’s performance, which is essentially an upper-bound estimate.

### Benchmarking existing condition disentangling methods

We attempted to compare ALPINE to existing condition disentangling methods: biolord (Piran et al. 2024), scDisco (R Liu et al. 2024), scDisInFact (Z Zhang et al. 2024), scINSIGHT (Qian et al. 2022), and scParser (Zhao et al. 2024). Unfortunately, we were unable to successfully instantiate a biolord model for training and downstream analysis. We also repeatedly encountered NaNs when attempting to run scDisco so we were unable to obtain comparable results. Meanwhile, scINSIGHT required extensive training time (>24 hours) even on small datasets. As such, these 3 methods were excluded from downstream comparisons. We also note that though scParser is conceptually designed to handle multiple batch and covariate inputs, we encountered an index out of bounds issue from their C++ implementation when providing inputs in that way; we did discover that we could combine multiple condition labels into a single covariate to run scParser successfully and thus used it in this way. We adhered to the recommended hyperparameter settings for scDisInFact and used the optimization function in scParser to identify optimal parameters. To obtain reconstructed counts using scDisInFact, we found that the predicted counts functionality requires batch and covariate labels to be provided. We observed that when using the default values of *None* for both batch and covariate, the predicted counts performed very poorly, but using “control” and “healthy” as the reference labels (which theoretically do not include perturbation effects) consistently performed better. Thus, we used “control” and “healthy” as the given labels, and *None* for the batch label for all comparisons of scDisInFact’s reconstructed counts.

### Comparisons against differential gene expression analysis

To minimize variability in single-cell gene expression, we constructed pseudo-bulk replicates using the decoupler package (Badia-i-Mompel et al. 2022). For each cell type, we aggregated counts from randomly sampled cells to generate 10 pseudo-bulk replicates, each representing the sum of gene expression across a subset of cells. We compared using either ComBat (Johnson et al. 2007) or Scanorama (Hie et al. 2019) for batch correction (as well as not using batch correction at all). We then compared using either Wilcoxon rank-sum test or t-test for differential gene expression analysis in each of the simulated datasets. Note that for differential gene expression analysis, we can only compare groups of pseudo-bulk samples within cell types, so we get one set of gene weights per cell type (versus disentanglement methods, which provide gene weights per condition / label). AUPRC for identifying known perturbed genes was calculated using the log fold-change.

### Benchmarking existing batch removal tools

We compare ALPINE to 7 existing single cell batch removal methods, Seurat (Stuart et al. 2019), Harmony (Korsunsky et al. 2019), LIGER (J Liu et al. 2020), scVI (Lopez et al. 2018), Scanorama (Hie et al. 2019), MNN (Haghverdi et al. 2018), ComBat (Johnson et al. 2007) and 2 condition disentangling methods, scParser (Zhao et al. 2024) and scDisInFact (Z Zhang et al. 2024), for batch effect removal tasks. Using Scanpy (Wolf et al. 2018), we filter out genes present in fewer than 5 cells and cells expressing fewer than 300 genes in each dataset. For models assuming a zero-inflated negative binomial distribution, such as scDisInFact and scVI, we provide raw counts as input. For Seurat, Harmony, LIGER, and scParser, we provide log-transformed data as required. For scDisInFact, which requires an additional condition beyond batch for execution, we create a dummy condition by assigning the same label to all cells.

### Data preprocessing and analysis for the brain cancer dataset

The brain cancer dataset (Abdelfattah et al. 2022) consists of 201,986 single cells from 18 patients, including glioma, immune, and stromal cells. We obtained the preprocessed dataset from the Single Cell Portal (Tarhan et al. 2023) and selected the top 2,000 highly variable genes, with patient identity as the batch variable. We randomly subsampled 50,000 cells stratified by patient for benchmarking. In addition to patient, this dataset includes covariates for sex and cancer type, and we included all 3 as guided covariates to ALPINE and scParser. scDisInFact requires distinguishing between batch versus covariates of interest, so we designated patient as batch and sex and cancer type as covariates of interest. We evaluated cell clustering using the same metrics as in the simulations; beyond ARI and NMI, we also calculated 1-ARI/1-NMI to evaluate batch and covariate mixing. To identify genes associated with each provided label, we extracted all nonzero gene weights from each method and standardized them as z-scores. To compare the overlap of top-associated genes from each method using upset plots (Conway et al. 2017), we used a 1.96 z-score threshold (corresponding to p-value=0.05 in a two-sided test or p-value=0.025 in a one-wided test). This process is relatively straightforward for ALPINE, which provides non-negative gene weights. The gene weights that scParser can be positive or negative, so we opted to create two separate lists to preserve the sign in downstream comparisons: one for positive gene scores where z ≥ 1.96, and another for negative gene scores where z ≤ −1.96. scDisInFact already provides functionality to consider both positive and negative values to transform the raw weights into non-negative scores, but as we have already previously mentioned, it can only identify covariate-level associations and not genes associated with each label. Thus, we used the same covariate-associated genes for comparison for the label-associated genes from ALPINE and scParser. For example, when comparing male- and female-associated gene lists, scDisInFact uses the sex-associated genes that they find for both.

### Data preprocessing and analysis for the adipose tissue dataset

The adipose tissue dataset (Emont et al. 2022) includes both human and mouse data, sequenced on two platforms, single-cell RNA-seq and single-nucleus RNA-seq, with 166,149 human cells and 197,721 mouse cells initially collected. For analysis, we subsampled 50,000 cells from each species, totaling 100,000 cells (27.48% of the dataset). To combine the two species’ single-cell data, we used orthologous gene mapping, which yielded 15,417 shared genes. The combined data was then normalized and log-transformed to unify library sizes across species. We use the provided batch (i.e., organism) and biological condition variables (i.e., assay, sex, depot tissues) as guided variables to ALPINE for signal decomposition. When attempting to compare with scParser, the package would crash whenever we would try running it with the full set of labels (one batch and three covariates), so we were unable to include these comparisons. In comparing our results with those of scDisInFact, we used the same guided labels and designated the assay as the primary batch, while treating the remaining variables as covariates of interest for training the scDisInFact model. To extract important genes associated with labels from scDisInFact, we first transformed the gene scores into z-scores, then calculated the corresponding p-values, followed by False Discovery Rate (FDR) correction. We applied the same procedure to the gene signatures identified by ALPINE and used an FDR threshold of ≤ 0.05?to select significant genes.

### Software implementation

ALPINE is built on top of the Scanpy ecosystem, so all data structures and matrix operations are native Scanpy AnnData objects. We also provide utility functions to export the guided and unguided embeddings (and associated gene weights), as well as reconstructed counts into Seurat-compatible formats, enabling seamless downstream analysis in either Scanpy or Seurat workflows.

### Software availability

Our implementation of ALPINE package is available on Github at https://github.com/ylaboratory/ ALPINE, with additional analysis code and tutorials available at https://github.com/ylaboratory/ ALPINE-analysis, both released under the BSD 3-clause license for open source use. We also include a snapshot of the repositories as Supplemental Code.

## Competing interest statement

The authors declare no competing interests.

## Acknowledgments

The authors would like to thank members of the ylaboratory for helpful discussions.

## Funding

This work was supported by the Cancer Prevention & Research Institute of Texas (CPRIT RR190065) and the AI2Health research cluster within the Ken Kennedy Institute at Rice University. VY is a CPRIT Scholar in Cancer Research.

## Author contributions

WHL, LL, and VY conceived the project and developed the method. WHL and LL implemented the method and benchmarked it. WHL, LL, and RD analyzed results. VY supervised the study. All authors wrote and reviewed the manuscript.

## Notes

### Competing Interest Statement

The authors have declared no competing interest.

### Summary of Updates

Figure 2 revised, additional Supplemental material, additional results, clarifications in methods

