## Supplemental Materials for "Interpretable phenotype decoding from multi-condition sequencing data with ALPINE"

### Contents

**Supplemental Methods: Derivation details for the ALPINE framework** 1

**Supplemental Figures** 6

**Supplemental Tables** 21

### Supplemental Methods: Derivation details for the ALPINE framework

To briefly recap, we represent a log-transformed single-cell RNA expression dataset as $X \in \mathbb{R}^{m\times n}$ , where $m$ is the number of genes and $n$ is the number of cells. ALPINE decomposes $X$ into a non-negative gene feature matrix $W \in \mathbb{R}^{m\times k}$ and a cell embedding matrix $H \in \mathbb{R}^{k\times n}$, where $k$ is the total number of latent components. To incorporate label information and guide the decomposition, a subset of the gene signatures in $W$ are supervised by user-supplied covariates. We define the corresponding submatrices as $W^{(i)}_{\text{guided}} \in \mathbb{R}^{m\times k_i}$ and $H^{(i)}_{\text{guided}} \in \mathbb{R}^{k_i\times n}$, such that the total number of components is distributed as $\sum_{i=1}^{c} k_i + k_{\text{unguided}} = k$, where $c$ represents the number of covariates specified by the user. The unguided signatures correspond to components that are not supervised by any label information. These components preserve cell-level variations unaffected by batch or covariate influence. The matrices $W$ and $H$ can be expressed as stacked forms of the guided and unguided signatures:

$$W = \begin{bmatrix} | & | & & | & | \\ W^{(1)}_{\text{guided}} & W^{(2)}_{\text{guided}} & \cdots & W^{(c)}_{\text{guided}} & W_{\text{unguided}} \\ | & | & & | & | \end{bmatrix} \quad \text{and} \quad H = \begin{bmatrix} - & H^{(1)}_{\text{guided}} & - \\ - & H^{(2)}_{\text{guided}} & - \\ & \vdots & \\ - & H^{(c)}_{\text{guided}} & - \\ - & H_{\text{unguided}} & - \end{bmatrix}.$$

To incorporate guided covariate information, we transform each covariate into a one-hot encoded matrix, denoted as $Y^{(i)} \in \mathbb{R}^{d_i\times n}$, where $d_i$ represents the number of unique labels in the covariate. We then introduce a weight matrix $B^{(i)} \in \mathbb{R}^{d_i\times k}$, which models the relationship between the signatures and labels. The goal is to approximate $B^{(i)}H^{(i)}_{\text{guided}}$ such that it aligns with the one-hot encoded matrix $Y^{(i)}$. Thus, the full optimization objective for the ALPINE model is formulated as:

$$\underset{W,H\geq 0}{\arg\min} \left[ ||X - WH||_F^2 + \sum_{i=1}^{c} \lambda_i D_{KL}(Y^{(i)}||B^{(i)}H^{(i)}_{\text{guided}}) + J(W) \right],$$

where the first term, $||X - WH||_F^2$, represents the reconstruction error, ensuring that the decomposition $WH$ closely approximates the original matrix $X$. The second term, which we also call the prediction term, $\sum_{c=1}^{c} \lambda_i D_{KL}(Y^{(i)}||B^{(i)}H^{(i)}_{\text{guided}})$, uses the generalized Kullback-Leibler (KL) divergence [1] to enforce alignment between the guided submatrices $H^{(i)}_{\text{guided}}$ and their corresponding label matrices $Y^{(i)}$:

$$D_{KL}(Y^{(i)}||B^{(i)}H^{(i)}_{guided}) = \sum_m \sum_n (Y_{mn} \log(\frac{Y_{mn}}{(B^{(i)}H^{(i)}_{guided})|_{mn}}) - Y_{mn} + (B^{(i)}H^{(i)}_{guided})|_{mn})$$

As an aside, we note that generalized KL divergence $D_{KL}(A||B)$ may be a slightly confusing name for this measure as $Y$ and $BH$ are not probability distributions, but we are following the naming convention in [1], which seemed to have opted for this name because the measure reduces to the standard KL-divergence when $\sum_{ij} A_{ij} = \sum_{ij} B_{ij} = 1$.

The last term, $J(W)$, is a regularization term for the matrix $W$, which controls the complexity of the learned gene features, which we discuss in more detail below. This formulation balances the need for a good reconstruction of the expression matrix $X$ while ensuring that the guided components align with the provided label information. By using KL divergence instead of the traditional Frobenius norm in the prediction loss term, we refine the approach to better handle probabilistic label alignment, which is more appropriate when dealing with categorical data.

To derive the multiplicative update rules, we first compute the partial derivatives of the objective function with respect to each matrix. For each guided submatrix $H^{(i)}_{\text{guided}}$, the partial derivative of the reconstruction term is given by:

$$\frac{\partial ||X - WH||_F^2}{\partial H^{(i)}_{\text{guided}}} = -2W^{(i)\top} X + 2W^{(i)\top} W^{(i)} H^{(i)}_{\text{guided}},$$

where $W^{(i)}$ corresponds to the factors associated with $H^{(i)}_{\text{guided}}$.

The partial derivative of the prediction term is:

$$\frac{\partial D_{KL}(Y^{(i)} || B^{(i)} H^{(i)}_{\text{guided}})}{\partial H^{(i)}_{\text{guided}}} = -B^{(i)\top} \frac{Y^{(i)}}{B^{(i)} H^{(i)}_{\text{guided}}} + B^{(i)\top} \mathbf{1}.$$

By combining these two terms, we use the multiplicative update method by Lee and Seung [1] to derive the update rule for $H^{(i)}_{\text{guided}}$:

$$H^{(i)}_{\text{guided}} \leftarrow H^{(i)}_{\text{guided}} \odot \frac{2W^{(i)\top} X + \lambda^{(i)} B^{(i)\top} \frac{Y^{(i)}}{B^{(i)} H^{(i)}_{\text{guided}}}}{2W^{(i)\top} W^{(i)} H^{(i)}_{\text{guided}} + \lambda^{(i)} B^{(i)\top} \mathbf{1}}.$$

For the unguided submatrix $H_{\text{unguided}}$, only the reconstruction term contributes, which leads to the following update rule:

$$H_{\text{unguided}} \leftarrow H_{\text{unguided}} \odot \frac{2W^\top X}{2W^\top W H_{\text{unguided}}}.$$

Since the guided signatures $H^{(i)}_{\text{guided}}$ and unguided signatures $H_{\text{unguided}}$ are independent, we can combine them into a complete $H$ matrix by merging the corresponding numerator and denominator terms. The multiplicative update rule for the entire $H$ matrix is thus:

$$H \leftarrow H \odot \frac{2W^\top X + P}{2W^\top W H + Q},$$

where

$$P = \begin{bmatrix} P_1 & P_2 & \cdots & P_g & \mathbf{0}_{H_{\text{unguided}}} \end{bmatrix}, \quad \text{and} \quad P_i = \lambda_i B^{(i)}{}^\top \left( \frac{Y^{(i)}}{B^{(i)} H^{(i)}_{\text{guided}}} \right)$$

and

$$Q = \begin{bmatrix} Q_1 & Q_2 & \cdots & Q_g & \mathbf{0}_{H_{\text{unguided}}} \end{bmatrix}, \quad \text{where} \quad Q_i = \lambda_i B^{(i)}{}^\top \mathbf{1}_{Y^{(i)}}.$$

In this formulation, $P$ represents the vertically stacked numerator terms from all guided components, and $Q$ represents the vertically stacked denominator terms. Since the unguided part is not regularized by the labels, the corresponding submatrices in $P$ and $Q$ are assigned zero values, ensuring that the shapes of $P$ and $Q$ match the structure of $H$.

Before introducing the multiplicative update for $W$, we define the regularization term $J(W)$, which incorporates LASSO, Ridge, and orthogonality constraints to enhance the uniqueness and generalizability of gene signatures which refers from the Lin & Boutros's matrix regularization part [2]. This regularization is expressed as:

$$J(W) = \underbrace{\alpha(l1_{\text{ratio}} \sum_{i,j} |W_{i,j}| + \frac{(1 - l1_{\text{ratio}})}{2} ||W||_2^2) +}_{\text{Elastic net regularization}} \quad \underbrace{\beta \sum_{i<j} W_{\cdot i}^\top W_{\cdot j}}_{\text{orthogonal regularization}}$$

where $\alpha$ controls the strength of Elastic net regularization, and $l1_{\text{ratio}}$ balances LASSO and Ridge penalties. The orthogonality term, weighted by $\beta$, reduces similarity between signatures to promote diversity. The partial derivatives of the regularization equation for each term with respect to W are as follows:

$$\frac{\partial}{\partial W} \sum_{i,j} |W_{i,j}| = \frac{\partial}{\partial W} tr(WE) = E^\top$$

$$\frac{\partial}{\partial W} \frac{1}{2} \|W\|_2^2 = \frac{\partial}{\partial W} \frac{1}{2} tr(WW^\top) = W$$

$$\frac{\partial}{\partial W} \sum_{i<j} W_{i\cdot} W_{\cdot j} = \frac{\partial}{\partial W} \frac{1}{2} tr(W(E - I)W^\top) = W(E - I)$$

where $E$ is the ones matrix with the appropriate dimension and $I$ is the identity matrix.

The gradient of $||X - WH||_F^2$ with respect to $W$ is:

$$\frac{\partial ||X - WH||_F^2}{\partial W} = -2XH^\top + 2WHH^\top.$$

Incorporating regularization, the multiplicative update for $W$ is thus:

$$W \leftarrow W \odot \frac{2XH^\top}{2WHH^\top + \alpha(1 - l1_{\text{ratio}})W + \beta W(\mathbf{1}_{k\times k} - I_{k\times k})) + \alpha l1_{\text{ratio}} \mathbf{1}_{m\times k}}.$$

Here, $H$ includes both guided and unguided submatrices, ensuring a structured decomposition.

For each $B^{(i)}$ associated with the $i$-th guided submatrix $H_{\text{guided}}^{(i)}$, the partial derivative of the KL divergence is:

$$\frac{\partial D_{KL}(Y^{(i)} || B^{(i)} H_{\text{guided}}^{(i)})}{\partial b_{i,j}} = -\sum_n \left( \frac{x_{i,n} h_{j,n}}{B^{(i)} H_{\text{guided}}^{(i)}|_{i,n}} \right) + \sum_n h_{j,n}.$$

In matrix form, this simplifies to:

$$\frac{\partial D_{KL}(Y^{(i)} || B^{(i)} H_{\text{guided}}^{(i)})}{\partial B^{(i)}} = -\frac{Y^{(i)}}{B^{(i)} H_{\text{guided}}^{(i)}} {H_{\text{guided}}^{(i)}}^\top + \mathbf{1} {H_{\text{guided}}^{(i)}}^\top.$$

Using the multiplicative update rule, the update for $B^{(i)}$ is:

$$B^{(i)} \leftarrow B^{(i)} \odot \frac{\frac{Y^{(i)}}{B^{(i)} H_{\text{guided}}^{(i)}} H_{\text{guided}}^{(i)}{}^{\top}}{\mathbf{1} H_{\text{guided}}^{(i)}{}^{\top}}.$$

This update ensures that $B^{(i)}$ is adjusted iteratively to better fit the data while maintaining non-negativity.

### Supplemental Figures

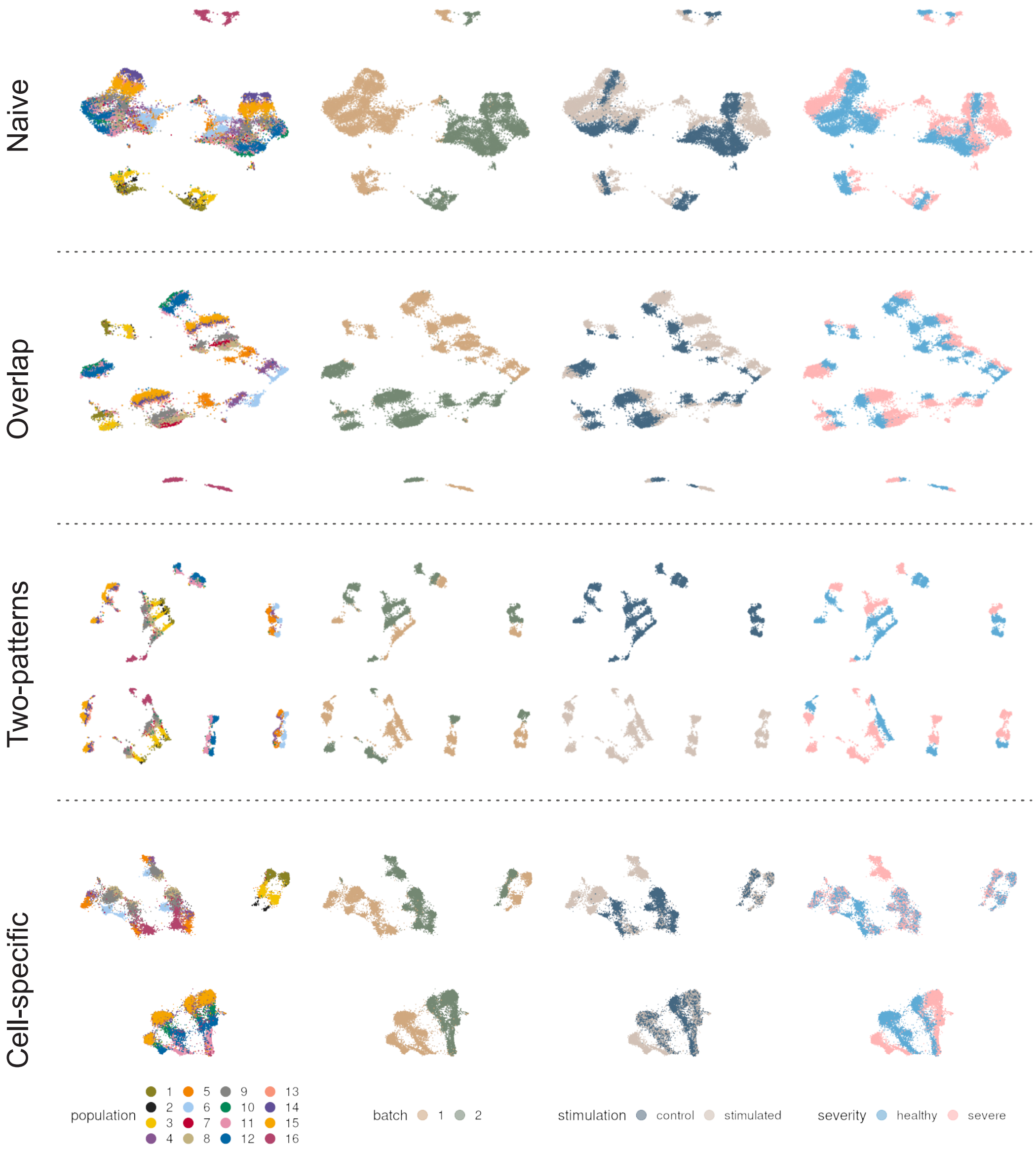


**Supplemental Figure 1. UMAP visualization of the simulation datasets.** In the simulation datasets, each scenario includes 10 different simulations. We visualize the first single-cell sequencing dataset from each scenario. Different colors represent the population (cell types), batches, stimulation, and severity.

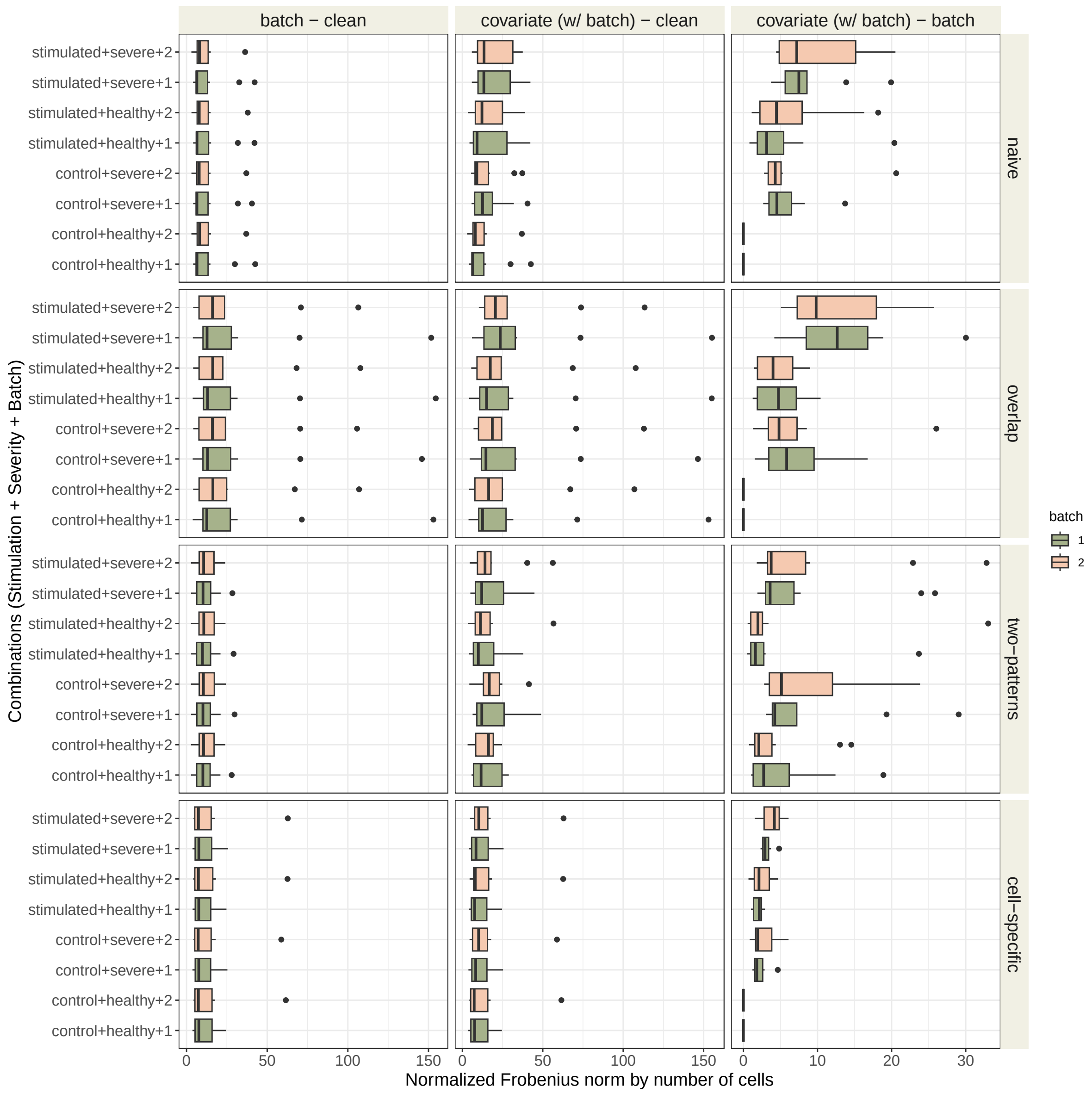


**Supplemental Figure 2. Effect sizes associated with batch effects and covariate influences.** The box plots compare the effect size for three conditions: a ’clean’ matrix, a matrix with added batch effects (’batch’), and a matrix with both covariate and batch effects (’covariate (w/ batch)’). The effect size is calculated as the Frobenius norm, which is normalized by the cell count to correct for the varying number of cells across the different label combinations shown on the y-axis. The box plots are colored by the batch.

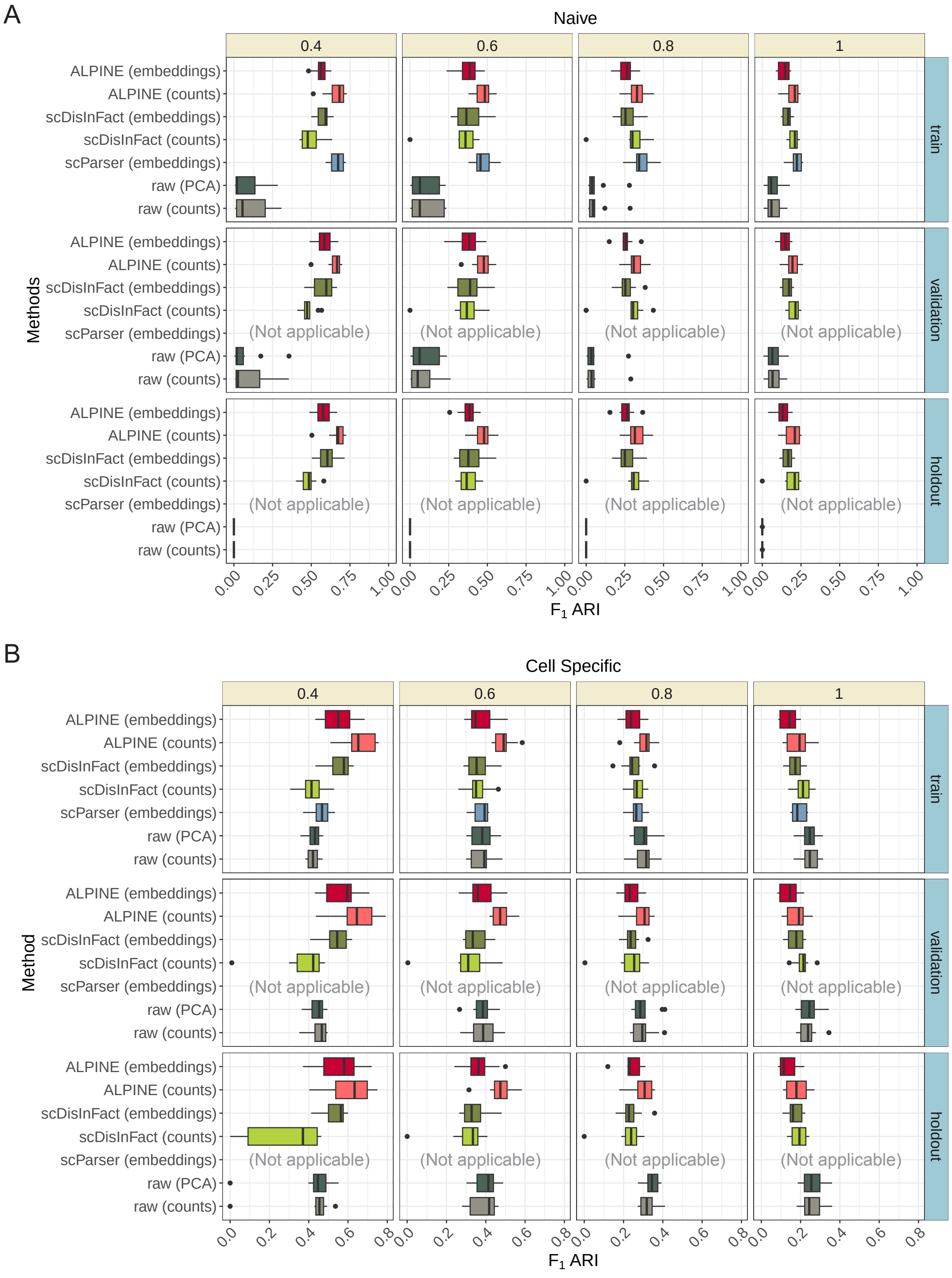


**Supplemental Figure 3. Cell clustering performance with varying extrinsic variability.** F1 ARI based on k-means clustering (with the known number of cell types) for training, validation, and holdout sets are shown for the **(A)** naive and **(B)** cell-specific scenarios across different values of $\sigma$, a hyperparameter in Symsim that controls the within cell-type variability of gene expression. ALPINE (embedding and reconstructed counts) are compared against two existing disentanglement methods (scDisInFact and scParser), as well as two baseline approaches (raw, which uses the confounded counts including both batch and covariate effects directly, and raw (PCA), which uses the top 50 PCs of the raw counts). Note that scParser does not provide functionatly to generate reconstructed counts and cannot be applied to new datasets so only results for the embeddings are available.

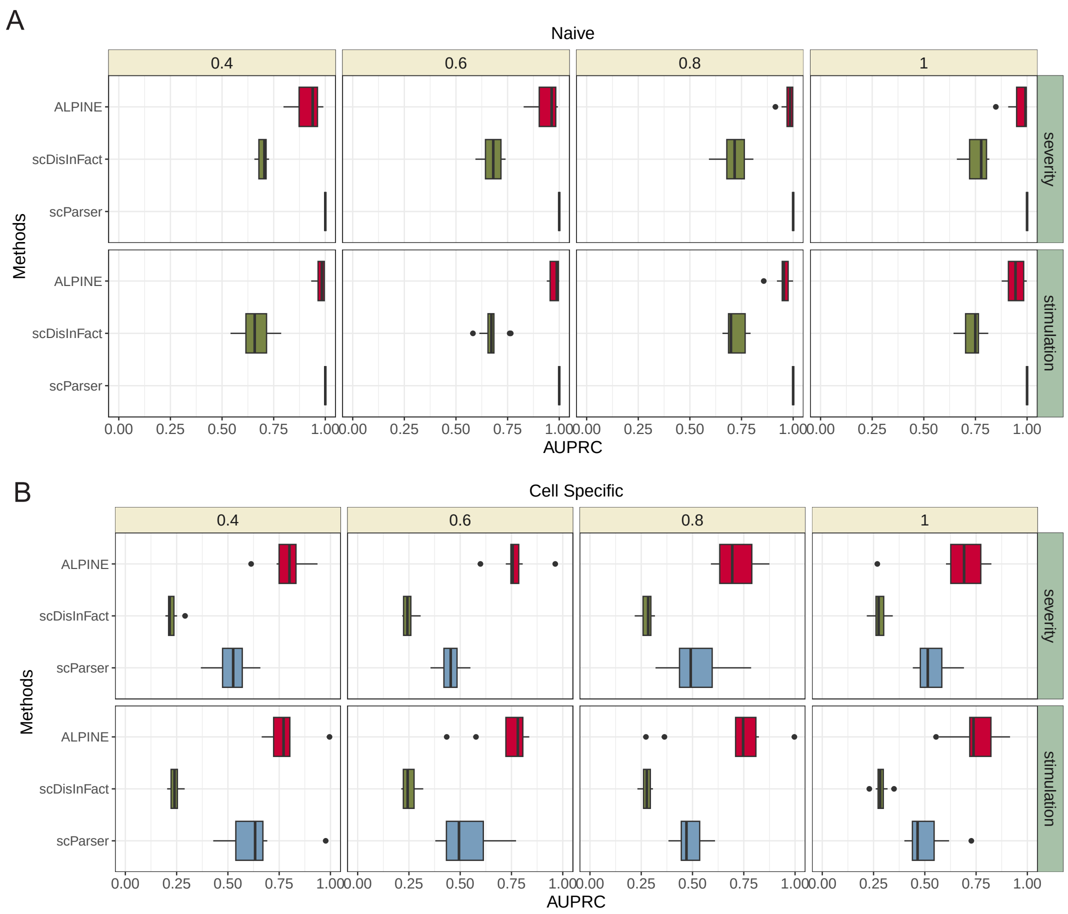


**Supplemental Figure 4. Performance comparisons evaluating ability to identify condition-associated genes at different levels of extrinsic variability.** AUPRC of ALPINE, scDisInFact, and scParser for the **(A)** naive and **(B)** cell-specific perturbation scenarios using different $\sigma$ values (0.4, 0.6, 0.8, 1.0), a hyperparameter in Symsim that controls the within cell-type variability of gene expression.

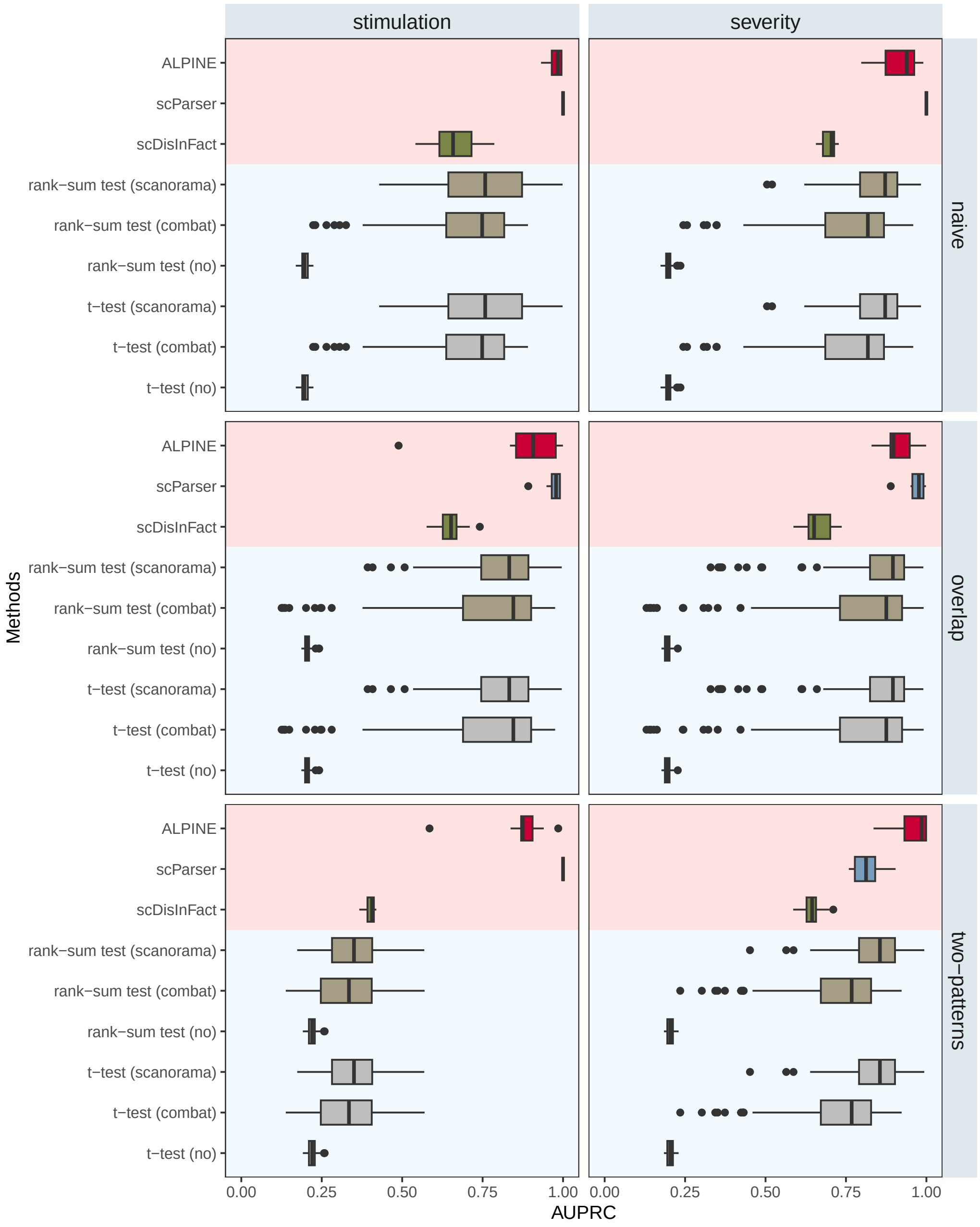


**Supplemental Figure 5. Comparison of differential expression (DE) analysis and disentanglement methods for identifying condition-associated genes in the naive, overlap, and two-patterns simulation scenarios.** Disentanglement methods (ALPINE, scParser, scDisInFact; red background) directly provide condition-associated gene weights. DE analysis was performed on pseudo-bulk cells using Wilcoxon rank-sum or t-tests, with batch correction by Scanorama, ComBat, or no correction (blue background).

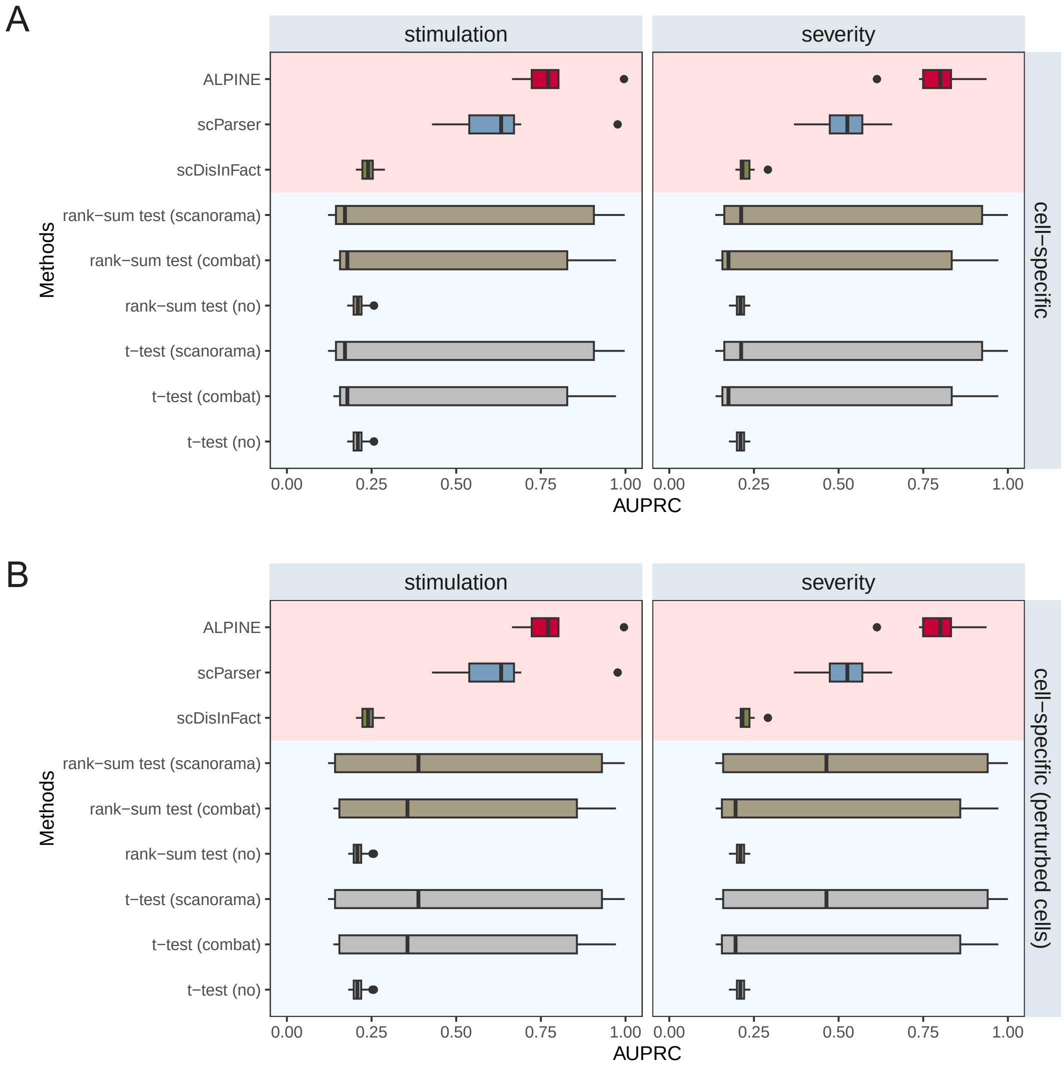


**Supplemental Figure 6. Comparison of differential expression (DE) analysis and disentanglement methods for identifying condition-associated genes in the cell-specific simulation scenario.** Disentanglement methods (ALPINE, scParser, scDisInFact; red background) directly provide condition-associated gene weights. DE analysis was performed on pseudo-bulk cells using Wilcoxon rank-sum or t-tests, with batch correction by Scanorama, ComBat, or no correction (blue background). ALPINE, scParser, and scDisInFact show the same results in both A and B, which show the DE results calculated with **(A)** all cell types, including those without perturbation, or **(B)** using only the perturbed cells. Note that in a non-simulation scenario, it is unlikely that a user would know exactly which cells are perturbed. However, we include B also for comparison purposes.

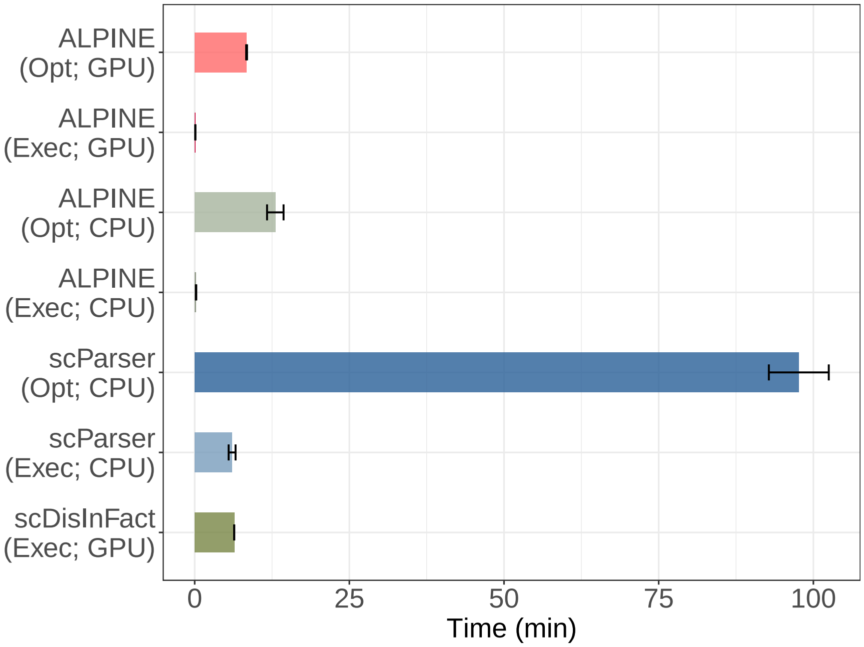


**Supplemental Figure 7. Time performance on the simulation datasets.** The bar plot illustrates the average time performance across different approaches. It is noteworthy that both ALPINE (which tested 100 hyperparameter sets) and scParser (which tested 5 hyperparameter sets) possess the optimization (Opt) process and execution (Exec) with optimal settings. Due to the absence of optimization code in scDisInFact, we utilized only the default parameters. Each bar includes 10 replicates for each scenario, resulting in a total of 40 measurements.

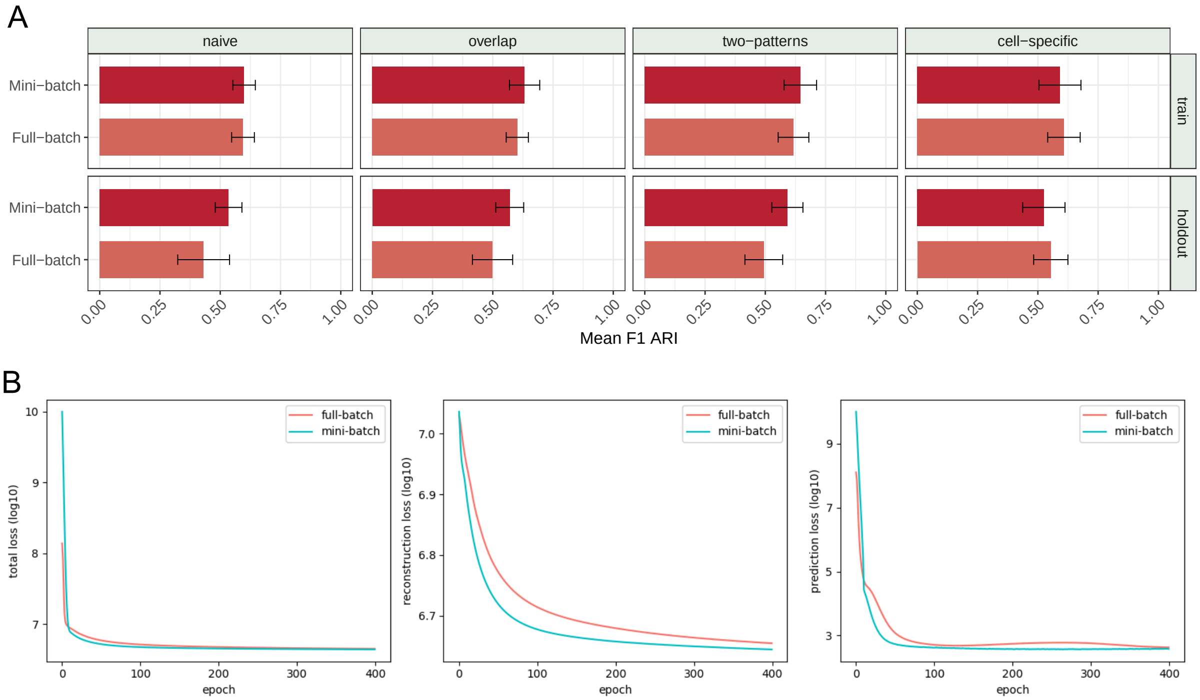


**Supplemental Figure 8. Comparison between using a mini-batch training strategy versus full dataset under different simulation scenarios. (A)** Average F1 ARI scores (considering cell cluster and batch integration) and standard errors across four simulated datasets using either mini-batch training or the entire dataset for both training and holdout sets. We see that the mini-batch training strategy typically results in slightly improved generalizability. **(B)** Line plots visualizing the progression of different loss terms (total is the sum of reconstruction and prediction losses) as the number of epochs increases, with colors indicating whether the models were trained using mini-batch or full-batch optimization. We observe that mini-batch training often converges quicker than full-batch training.

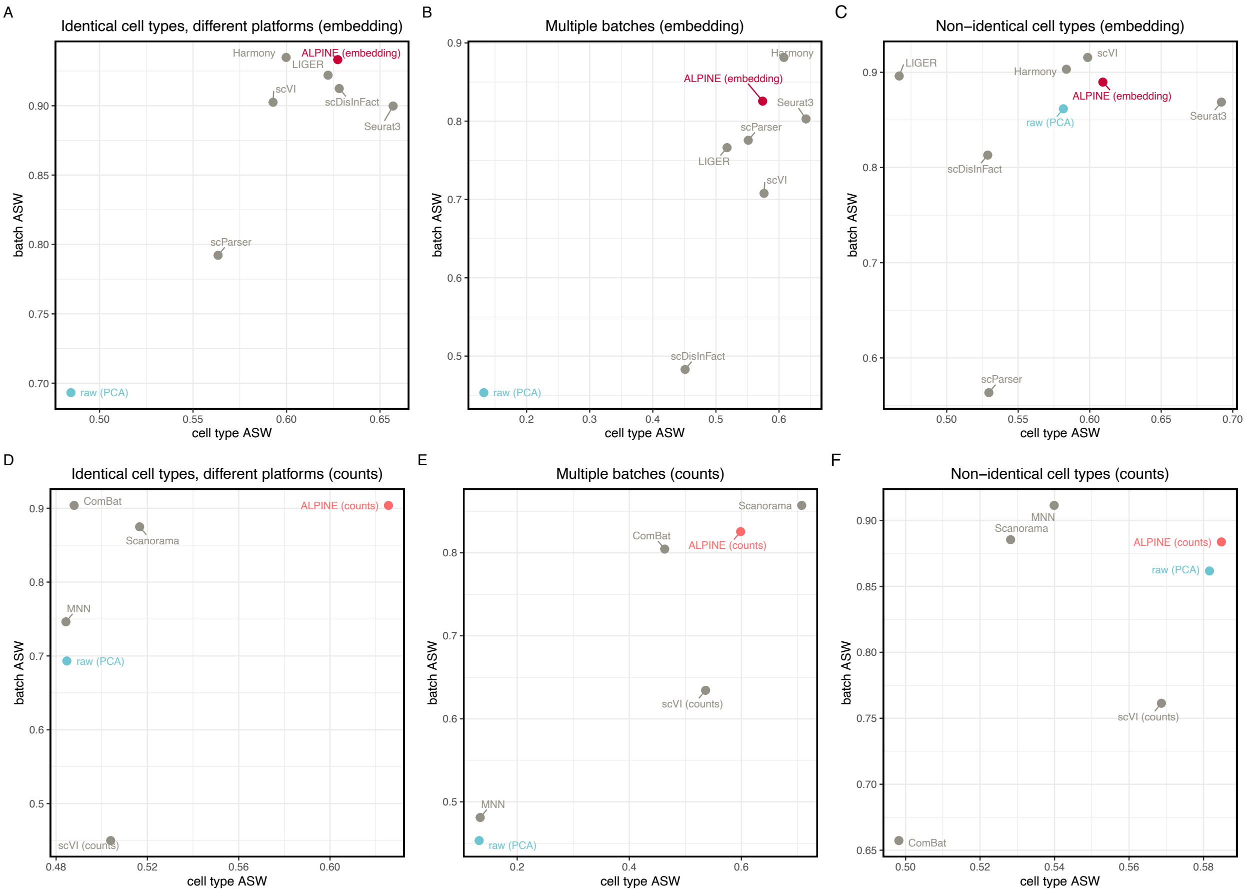


**Supplemental Figure 9. ASW score comparisons of ALPINE against other methods for batch effect removal and cell type annotation.** ASW of ALPINE versus scDisInfact, scParser, Seurat3, Harmony, Liger, scVI, Scanoramam MNN, and ComBat, in batch effect removal using three real datasets based on cell type clustering with Leiden (using default resolution=1). Methods producing low-dimensional embeddings (scDisInfact, scParser, Seurat3, Harmony, Liger, scVI) are compared with ALPINE embeddings in **(A)-(C)**, while methods reconstructing counts (Scanorama MNN, ComBat, and scVI (counts)) are compared with ALPINE-reconstructed counts in **(D)-(F)**. **(A)&(D)**. Human peripheral blood monouclear cell datasets with two batches and matched cell types. **(B)&(E)**. Pancreatic cells dataset with five batches and matched cell types. **(C)&(F)**. Mouse retina data with two batches and non-identical cell types.

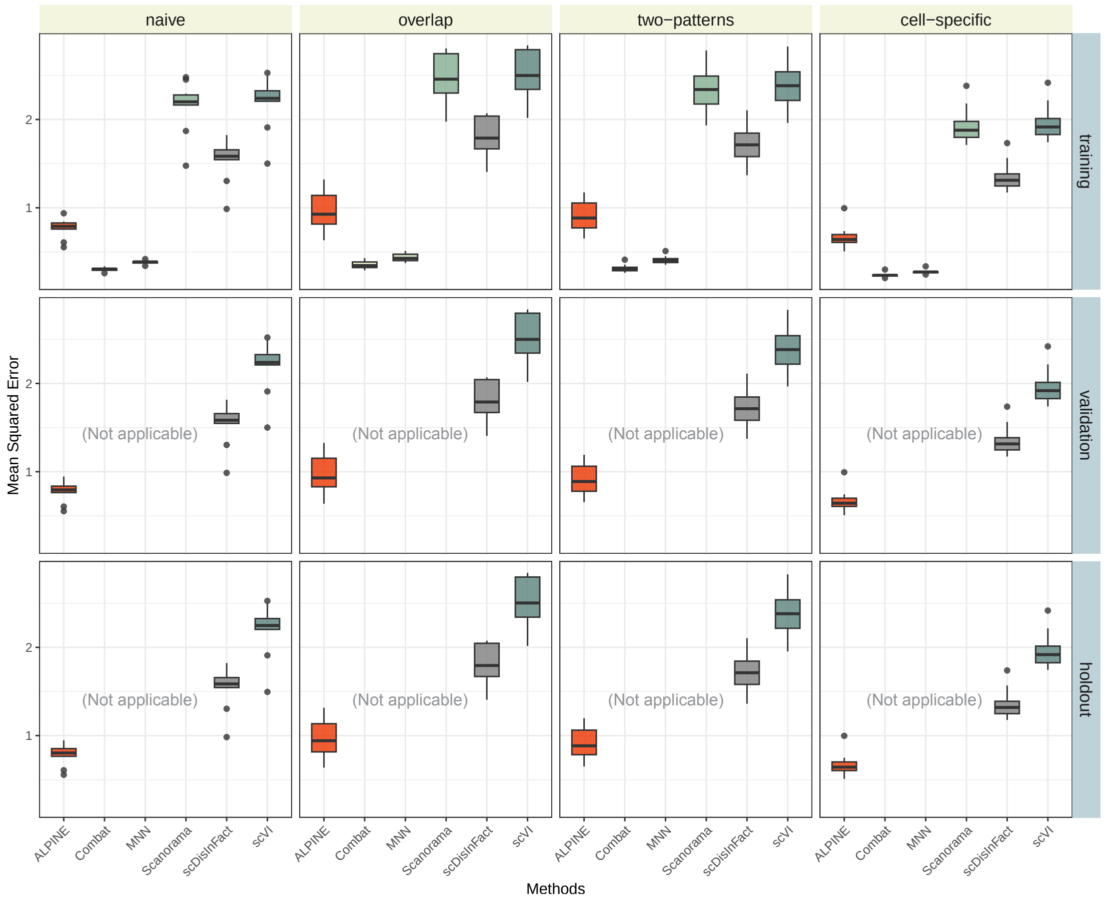


**Supplemental Figure 10. Comparison of reconstructed counts after batch removal in the simulation dataset.** The reconstructed counts from each method are compared against clean counts prior to adding batch and covariate effects in each simulation scenario. The box plots illustrate the mean squared errors (MSE) for each method across the four different scenarios (lower MSE indicates that the resulting counts are more similar to original counts). Since Combat, MNN, and Scanorama were specifically designed for batch removal, they lack the capability to apply learned information to eliminate batch effects from unseen datasets, validation, and holdout sets, which is why they are labeled as "Not applicable."

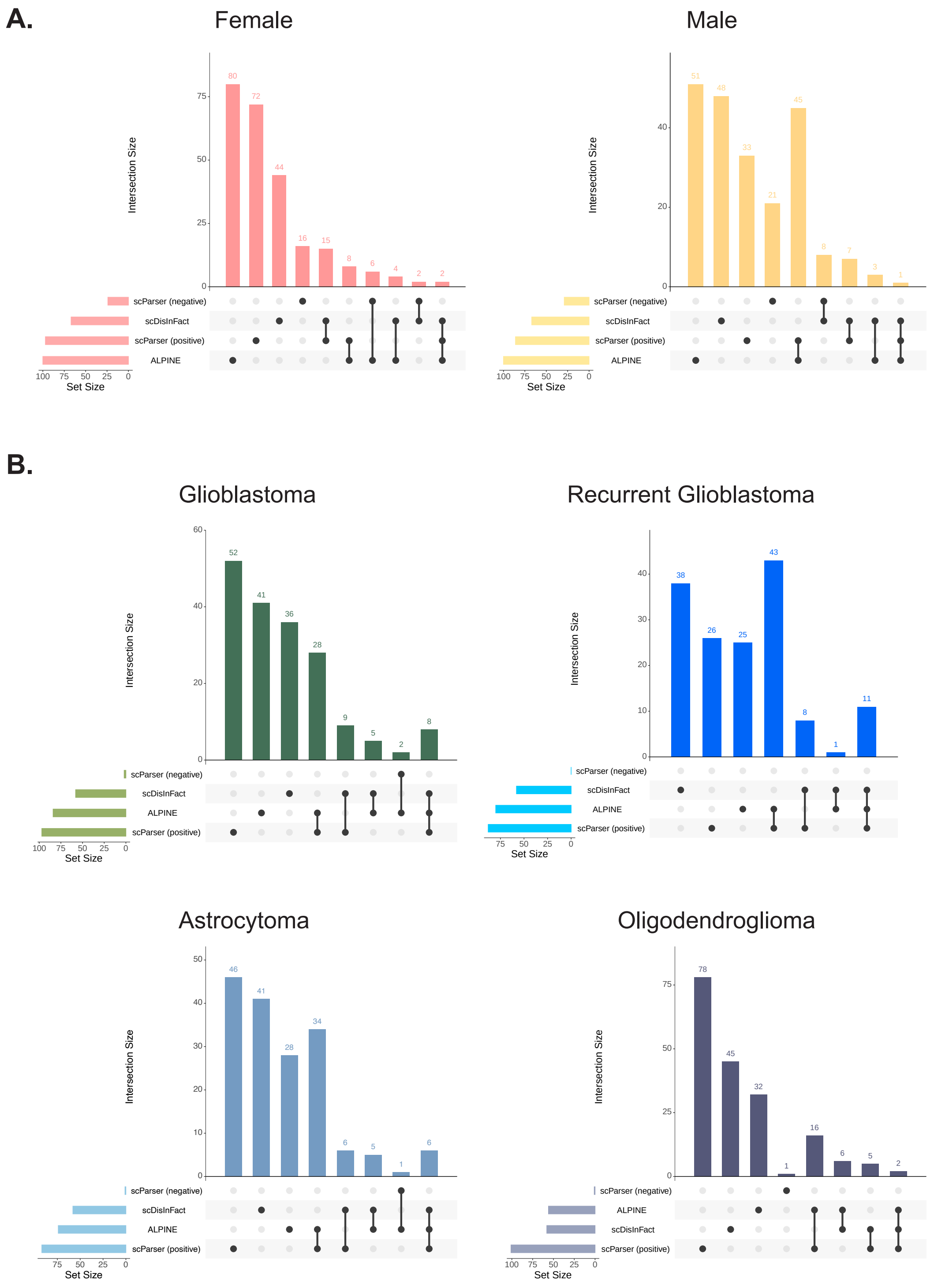


**Supplemental Figure 11. Analyzing similarity of identified condition-associated genes in the brain cancer dataset. (A)** Sex-associated genes. **(B)** Different cancer type-associated genes. Note that the scDisInFact genes within each covariate are the same, as there is no way to identify distinct label-associated genes (see Methods for further details).

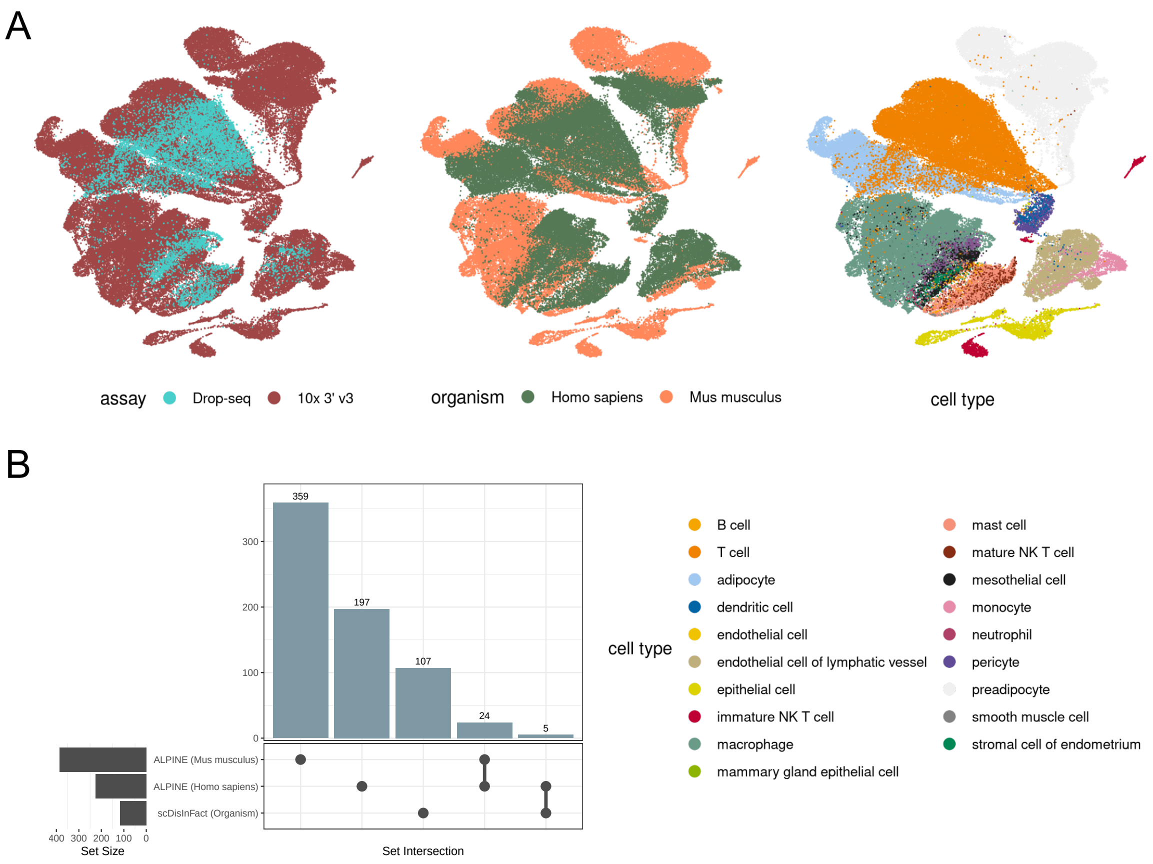


**Supplemental Figure 12. Comparing scDisInFact and ALPINE for the adipose tissue case study. (A)** The UMAP depicts the cell embeddings post-integration using scDisInFact, colored according to associated covariates, assay, organism, and cell type, with complete cell annotations displayed below. **(B)** The upset plot provides insight into the overlap of significant label-associated genes identified by ALPINE and scDisInFact. Due to ALPINE's design, covariate signatures can be directly associated with their corresponding labels, enhancing interpretability in the context of multi-label covariates, such as species (organism). In contrast, scDisInFact generates a single set of gene scores for the entire organism covariate, lacking label-specific resolution.

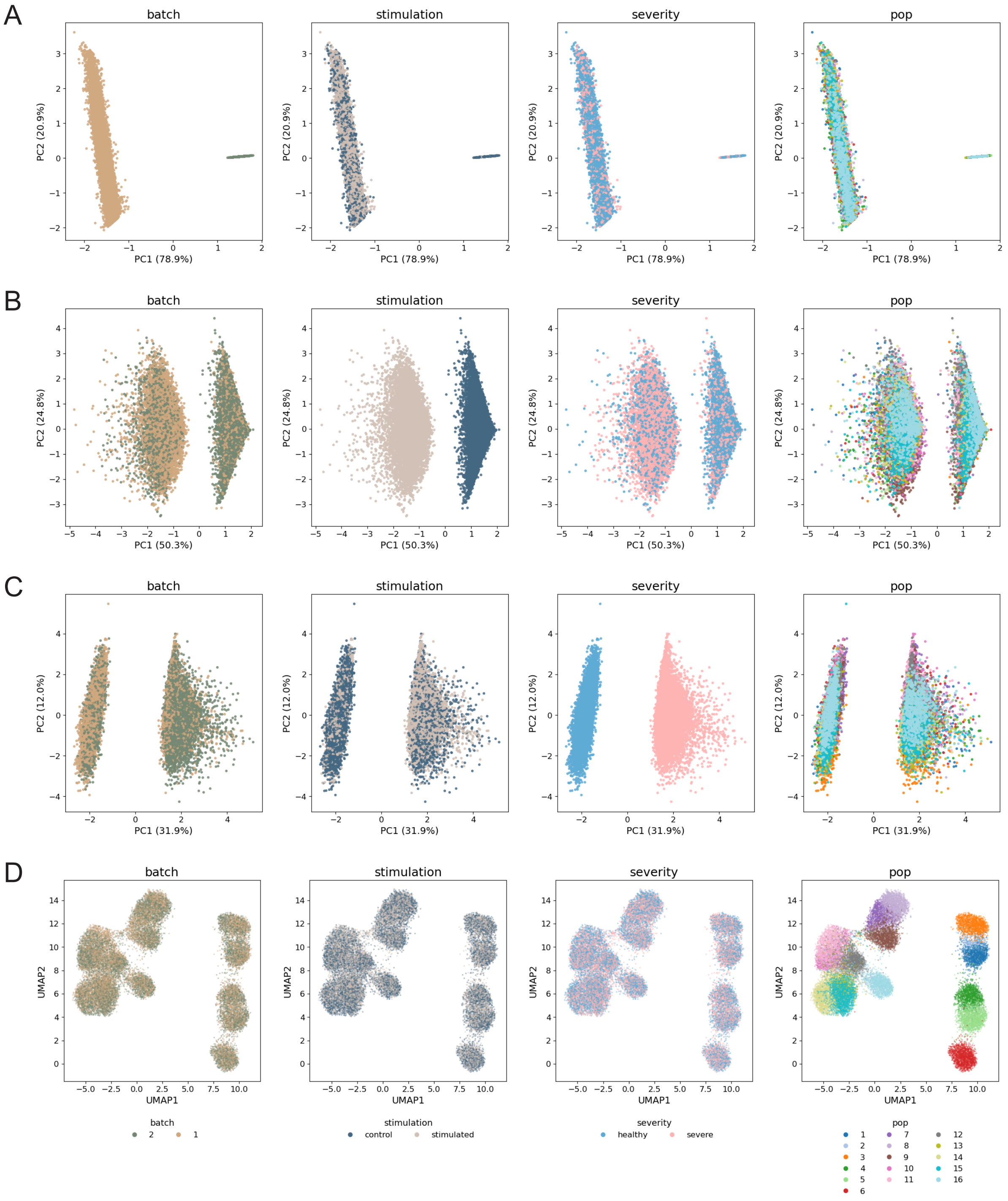


**Supplemental Figure 13. Visualization of guided and unguided embeddings based on an example training dataset from the naive simulation scenario.** The guided embeddings are visualized using PCA: **(A)** batch embeddings, **(B)** stimulation embeddings, and **(C)** severity embeddings. **(D)** UMAP plot representing the unguided embeddings. The different colors used in the plots highlight labels corresponding to distinct covariates.

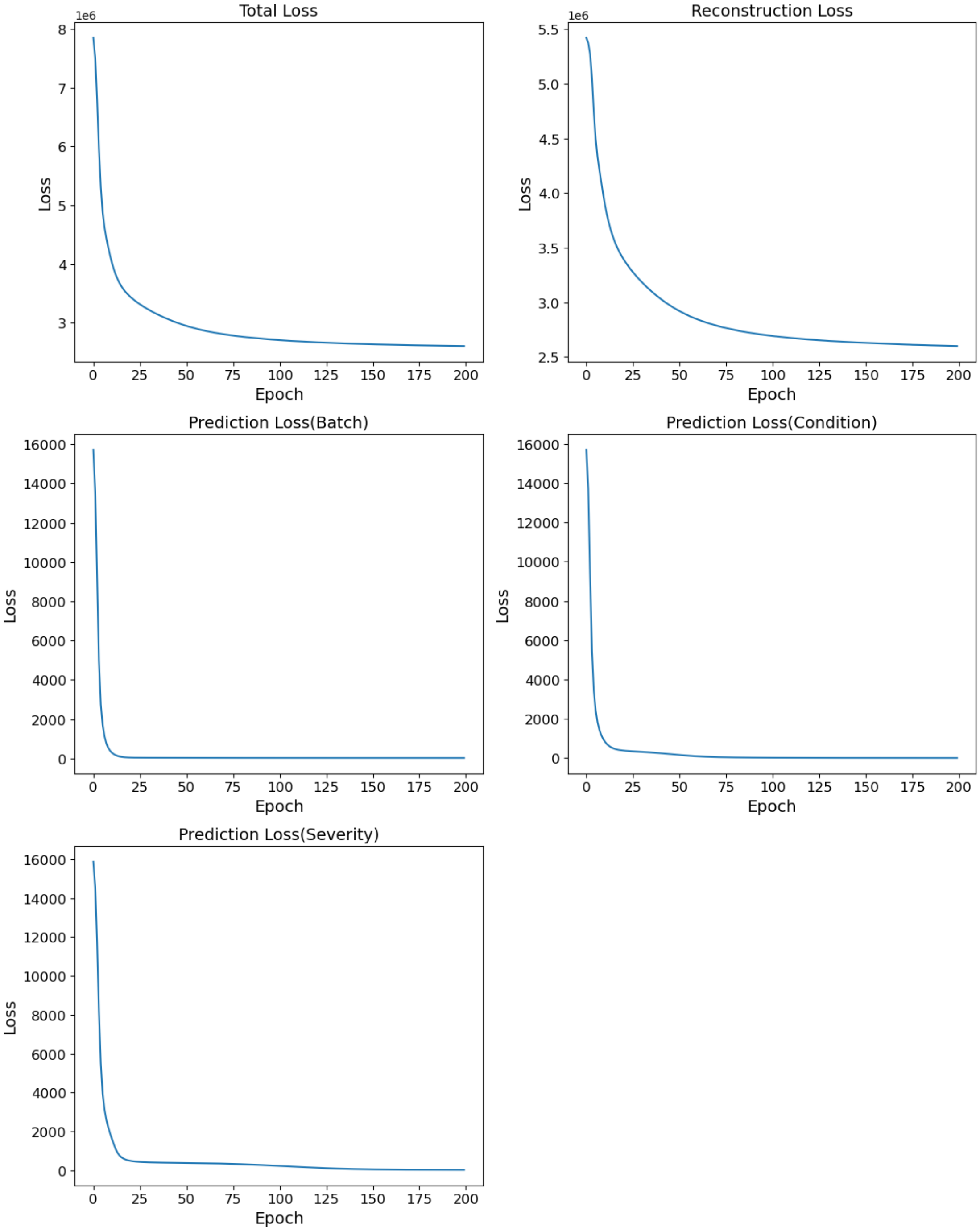


**Supplemental Figure 14. Training loss curves generated from an example run of ALPINE using the naive simulation dataset.** Illustrative example of training loss curves, including the total loss, reconstruction loss, and individual prediction losses for each covariate.

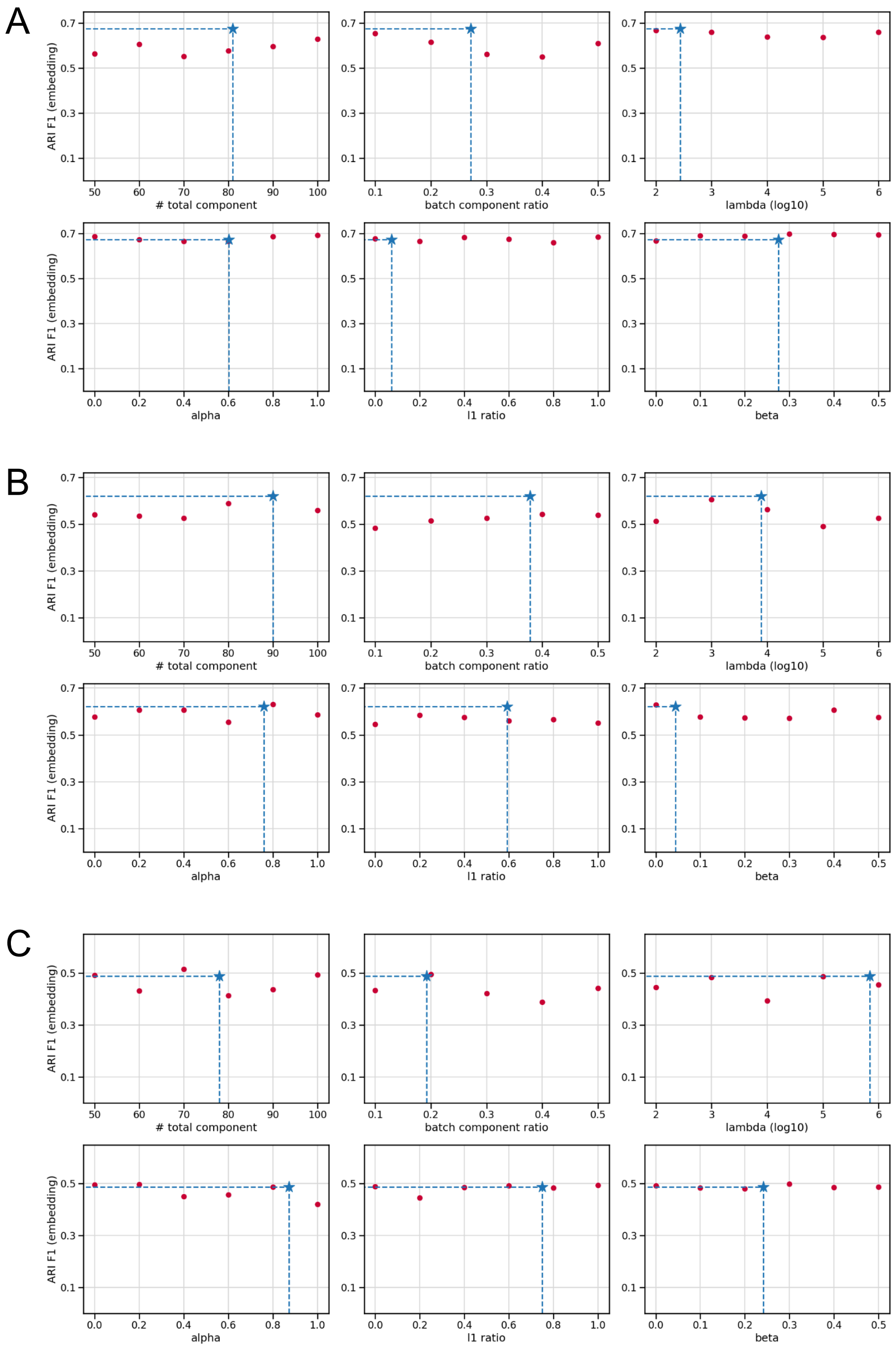


**Supplemental Figure 15. ALPINE's performance is robust to different hyperparameter choices.** In each subfigure, ARI F1 using ALPINE with alternative hyperparameters is shown. The blue star represents ALPINE with the optimized hyperparameter from a 50-epoch process, while each red dot shows the performance when varying a single hyperparameter, while keeping others fixed. **(A)**. Human peripheral blood monouclear cell datasets with two batches and matched cell types. **(B)**. Pancreatic cells dataset with five batches and matched cell types. **(C)**. Mouse retina data with two batches and non-identical cell types.

### Supplemental Tables

**Supplemental Table 1. Relative performance of ALPINE (embeddings and counts) to existing methods and baselines in the training dataset.** All test statistics and p-values are based on one-sided Wilcoxon rank sum test comparisons of $F_1ARI$.

| Set | Scenario | ALPINE variant | Comparison method | Statistic | p-value |
|---|---|---|---|---|---|
| train | naive | ALPINE (embeddings) | scDisInFact (embeddings) | 36.00 | 8.60e-01 |
| train | naive | ALPINE (embeddings) | scDisInFact (counts) | 82.00 | 7.34e-03 |
| train | naive | ALPINE (embeddings) | scParser (embeddings) | 4.00 | 1.00e+00 |
| train | naive | ALPINE (embeddings) | raw (counts) | 100.00 | 5.41e-06 |
| train | naive | ALPINE (embeddings) | raw (PCA) | 100.00 | 5.41e-06 |
| train | naive | ALPINE (counts) | scDisInFact (embeddings) | 84.00 | 4.47e-03 |
| train | naive | ALPINE (counts) | scDisInFact (counts) | 94.00 | 1.62e-04 |
| train | naive | ALPINE (counts) | scParser (embeddings) | 49.00 | 5.44e-01 |
| train | naive | ALPINE (counts) | raw (counts) | 100.00 | 5.41e-06 |
| train | naive | ALPINE (counts) | raw (PCA) | 100.00 | 5.41e-06 |
| train | overlap | ALPINE (embeddings) | scDisInFact (embeddings) | 45.00 | 6.58e-01 |
| train | overlap | ALPINE (embeddings) | scDisInFact (counts) | 78.00 | 1.77e-02 |
| train | overlap | ALPINE (embeddings) | scParser (embeddings) | 21.00 | 9.88e-01 |
| train | overlap | ALPINE (embeddings) | raw (counts) | 99.00 | 1.08e-05 |
| train | overlap | ALPINE (embeddings) | raw (PCA) | 99.00 | 1.08e-05 |
| train | overlap | ALPINE (counts) | scDisInFact (embeddings) | 57.00 | 3.15e-01 |
| train | overlap | ALPINE (counts) | scDisInFact (counts) | 88.00 | 1.44e-03 |
| train | overlap | ALPINE (counts) | scParser (embeddings) | 34.00 | 8.91e-01 |
| train | overlap | ALPINE (counts) | raw (counts) | 98.00 | 2.17e-05 |
| train | overlap | ALPINE (counts) | raw (PCA) | 99.00 | 1.08e-05 |
| train | two-patterns | ALPINE (embeddings) | scDisInFact (embeddings) | 64.00 | 1.57e-01 |
| train | two-patterns | ALPINE (embeddings) | scDisInFact (counts) | 85.00 | 3.42e-03 |
| train | two-patterns | ALPINE (embeddings) | scParser (embeddings) | 10.00 | 9.99e-01 |
| train | two-patterns | ALPINE (embeddings) | raw (counts) | 100.00 | 8.15e-05 |
| train | two-patterns | ALPINE (embeddings) | raw (PCA) | 100.00 | 8.93e-05 |
| train | two-patterns | ALPINE (counts) | scDisInFact (embeddings) | 82.00 | 7.34e-03 |
| train | two-patterns | ALPINE (counts) | scDisInFact (counts) | 94.00 | 1.62e-04 |
| train | two-patterns | ALPINE (counts) | scParser (embeddings) | 32.00 | 9.17e-01 |
| train | two-patterns | ALPINE (counts) | raw (counts) | 100.00 | 8.15e-05 |
| train | two-patterns | ALPINE (counts) | raw (PCA) | 100.00 | 8.93e-05 |
| train | cell-specific | ALPINE (embeddings) | scDisInFact (embeddings) | 44.00 | 6.85e-01 |
| train | cell-specific | ALPINE (embeddings) | scDisInFact (counts) | 91.00 | 5.25e-04 |
| train | cell-specific | ALPINE (embeddings) | scParser (embeddings) | 80.00 | 1.16e-02 |
| train | cell-specific | ALPINE (embeddings) | raw (counts) | 94.00 | 1.62e-04 |
| train | cell-specific | ALPINE (embeddings) | raw (PCA) | 91.00 | 5.25e-04 |
| train | cell-specific | ALPINE (counts) | scDisInFact (embeddings) | 85.00 | 3.42e-03 |
| train | cell-specific | ALPINE (counts) | scDisInFact (counts) | 99.00 | 1.08e-05 |
| train | cell-specific | ALPINE (counts) | scParser (embeddings) | 99.00 | 1.08e-05 |
| train | cell-specific | ALPINE (counts) | raw (counts) | 100.00 | 5.41e-06 |
| train | cell-specific | ALPINE (counts) | raw (PCA) | 100.00 | 5.41e-06 |

**Supplemental Table 2. Relative performance of ALPINE (embeddings and counts) to existing methods and baselines in the holdout dataset.** All test statistics and p-values are based on one-sided Wilcoxon rank sum test comparisons of $F_1ARI$.

| Set | Scenario | ALPINE variant | Comparison method | Statistic | p-value |
|---|---|---|---|---|---|
| holdout | naive | ALPINE (embeddings) | scDisInFact (embeddings) | 41.00 | 7.59e-01 |
| holdout | naive | ALPINE (embeddings) | scDisInFact (counts) | 91.00 | 5.25e-04 |
| holdout | naive | ALPINE (embeddings) | raw (counts) | 100.00 | 3.19e-05 |
| holdout | naive | ALPINE (embeddings) | raw (PCA) | 100.00 | 3.19e-05 |
| holdout | naive | ALPINE (counts) | scDisInFact (embeddings) | 79.00 | 1.44e-02 |
| holdout | naive | ALPINE (counts) | scDisInFact (counts) | 98.00 | 2.17e-05 |
| holdout | naive | ALPINE (counts) | raw (counts) | 100.00 | 3.19e-05 |
| holdout | naive | ALPINE (counts) | raw (PCA) | 100.00 | 3.19e-05 |
| holdout | overlap | ALPINE (embeddings) | scDisInFact (embeddings) | 49.00 | 5.44e-01 |
| holdout | overlap | ALPINE (embeddings) | scDisInFact (counts) | 75.00 | 3.15e-02 |
| holdout | overlap | ALPINE (embeddings) | raw (counts) | 97.00 | 1.64e-04 |
| holdout | overlap | ALPINE (embeddings) | raw (PCA) | 98.00 | 1.04e-04 |
| holdout | overlap | ALPINE (counts) | scDisInFact (embeddings) | 62.00 | 1.97e-01 |
| holdout | overlap | ALPINE (counts) | scDisInFact (counts) | 82.00 | 7.34e-03 |
| holdout | overlap | ALPINE (counts) | raw (counts) | 97.00 | 1.64e-04 |
| holdout | overlap | ALPINE (counts) | raw (PCA) | 98.00 | 1.04e-04 |
| holdout | two-patterns | ALPINE (embeddings) | scDisInFact (embeddings) | 67.00 | 1.09e-01 |
| holdout | two-patterns | ALPINE (embeddings) | scDisInFact (counts) | 86.00 | 2.60e-03 |
| holdout | two-patterns | ALPINE (embeddings) | raw (counts) | 100.00 | 7.47e-05 |
| holdout | two-patterns | ALPINE (embeddings) | raw (PCA) | 100.00 | 7.47e-05 |
| holdout | two-patterns | ALPINE (counts) | scDisInFact (embeddings) | 83.00 | 5.75e-03 |
| holdout | two-patterns | ALPINE (counts) | scDisInFact (counts) | 95.00 | 1.03e-04 |
| holdout | two-patterns | ALPINE (counts) | raw (counts) | 100.00 | 7.47e-05 |
| holdout | two-patterns | ALPINE (counts) | raw (PCA) | 100.00 | 7.47e-05 |
| holdout | cell-specific | ALPINE (embeddings) | scDisInFact (embeddings) | 60.00 | 2.41e-01 |
| holdout | cell-specific | ALPINE (embeddings) | scDisInFact (counts) | 93.00 | 6.55e-04 |
| holdout | cell-specific | ALPINE (embeddings) | raw (counts) | 81.00 | 9.27e-03 |
| holdout | cell-specific | ALPINE (embeddings) | raw (PCA) | 81.00 | 9.27e-03 |
| holdout | cell-specific | ALPINE (counts) | scDisInFact (embeddings) | 75.00 | 3.15e-02 |
| holdout | cell-specific | ALPINE (counts) | scDisInFact (counts) | 96.00 | 2.90e-04 |
| holdout | cell-specific | ALPINE (counts) | raw (counts) | 88.00 | 1.44e-03 |
| holdout | cell-specific | ALPINE (counts) | raw (PCA) | 88.00 | 1.44e-03 |

**Supplemental Table 3. Evaluation of batch and covariate removal performance using ALPINE, scDisInFact, and scParser on the brain cancer dataset.** The $1 - \text{ARI}$ and $1 - \text{NMI}$ performance metrics are calculated based on k-means clustering (with the known number of cell types) of the unguided embeddings.

| | ALPINE | | ALPINE (counts) | | scDisInFact | | scParser | |
|---|---|---|---|---|---|---|---|---|
| **Category** | **1 - ARI** | **1 - NMI** | **1 - ARI** | **1 - NMI** | **1 - ARI** | **1 - NMI** | **1 - ARI** | **1 - NMI** |
| Patients | 0.885 | 0.724 | 0.878 | 0.742 | 0.923 | 0.849 | 0.917 | 0.705 |
| Sex | 0.979 | 0.941 | 0.972 | 0.964 | 1.007 | 0.995 | 0.979 | 0.911 |
| Types | 1.011 | 0.988 | 0.969 | 0.942 | 0.971 | 0.994 | 1.015 | 0.978 |

**Supplemental Table 4. Evaluation of cell type clustering performance using ALPINE, scDisInFact, and scParser on the brain cancer dataset.** The ARI and NMI performance metrics are calculated based on k-means clustering (with the known number of cell types) of the unguided embeddings.

| | ALPINE | | ALPINE (counts) | | scDisInFact | | scParser | |
|---|---|---|---|---|---|---|---|---|
| **Category** | **ARI** | **NMI** | **ARI** | **NMI** | **ARI** | **NMI** | **ARI** | **NMI** |
| Cell | 0.413 | 0.531 | 0.359 | 0.472 | 0.352 | 0.457 | 0.171 | 0.335 |

**Supplemental Table 5. Evaluation of batch and covariate removal performance using ALPINE and scDisInFact in the adipose tissue case study.** The $1 - \text{ARI}$ and $1 - \text{NMI}$ performance metrics are calculated based on Leiden clustering (with a default resolution of 1) of the unguided embeddings.

| | ALPINE | | scDisInFact | |
|---|---|---|---|---|
| **Category** | **1 - ARI** | **1 - NMI** | **1 - ARI** | **1 - NMI** |
| Assay | 0.991 | 0.949 | 0.998 | 0.799 |
| Organism | 0.949 | 0.845 | 0.972 | 0.861 |
| Sex | 0.988 | 0.958 | 0.986 | 0.940 |
| Tissue | 0.935 | 0.828 | 0.946 | 0.799 |

**Supplemental Table 6. Evaluation of cell type clustering performance of ALPINE and scDisInFact in the adipose tissue case study.** The ARI and NMI performance metrics are calculated based on Leiden clustering (with a default resolution of 1) of the unguided embeddings.

| | ALPINE | | scDisInFact | |
|---|---|---|---|---|
| **Category** | **ARI** | **NMI** | **ARI** | **NMI** |
| Cell | 0.329 | 0.606 | 0.232 | 0.622 |
