## Supplemental Code for "Interpretable phenotype decoding from multi-condition sequencing data with ALPINE": main.py

```
import gc
import torch
import warnings
import numpy as np
import pandas as pd
import anndata as ad
import scanpy as sc
import numpy.typing as npt

from kneed import KneeLocator
from tqdm import tqdm
from typing import Optional, Union, List, Dict
from copy import copy, deepcopy
from contextlib import nullcontext
from dataclasses import dataclass
from .utils.encoder import FeatureEncoders
from .utils.sampling import (
    generate_epoch_indices,
    get_batch_indices,
    get_num_batches,
    create_joint_labels_from_dummy_matrices,
)

# Define a type alias for 32-bit float arrays
Float32Array = npt.NDArray[np.float32]


@dataclass
class AlpineMatrices:
    X: torch.Tensor
    Ys: List[torch.Tensor]
    Ws: List[torch.Tensor]
    Hs: List[torch.Tensor]
    Bs: List[torch.Tensor]

    def to_numpy(self) -> Dict[str, Union[Float32Array, List[Float32Array]]]:
        return {
            "X": self.X.cpu().numpy().astype(np.float32),
            "Ys": [y.cpu().numpy().astype(np.float32) for y in self.Ys],
            "Ws": [w.cpu().numpy().astype(np.float32) for w in self.Ws],
            "Hs": [h.cpu().numpy().astype(np.float32) for h in self.Hs],
            "Bs": [b.cpu().numpy().astype(np.float32) for b in self.Bs],
        }


class ALPINE:
    def __init__(
        self,
        n_components: int,
        n_covariate_components: List[int],
        lam: List[float],
        orth_W: float = 0.0,
        alpha_W: float = 0.0,
        l1_ratio_W: float = 0.0,
        use_als: bool = False,
        scale_needed: bool = True,
        loss_type: str = "kl-divergence",
        device: str = "cuda",
        eps: float = 1e-6,
        random_state: int = 42,
    ):
        self.n_components: int = n_components
        self.n_covariate_components: List[int] = n_covariate_components
        self.lam: List[float] = lam
        self.orth_W: float = orth_W
        self.alpha_W: float = alpha_W
        self.l1_ratio_W: float = l1_ratio_W
        self.use_als: bool = use_als
        self.scale_needed: bool = scale_needed
        self.device: torch.device = torch.device(device)
        self.loss_type: str = loss_type
        self.eps: float = eps
        self.random_state: int = random_state

        # Validate initialization arguments
        self._validate_init_args()

        # other useful attributes
        self.n_all_components = self.n_covariate_components + [self.n_components]
        self.total_components = sum(self.n_all_components)

    def fit(
        self,
        adata: ad.AnnData,
        covariate_keys: List[str],
        batch_size: Optional[int] = None,
        max_iter: Optional[int] = None,
        sampling_method: str = "random",
        verbose: bool = False,
    ) -> "ALPINE":
        # validate fit arguments and save them to class attribute
        self._validate_fit_args(
            adata, covariate_keys, batch_size, max_iter, sampling_method, verbose
        )
        self.feature_names: List[str] = adata.var_names.tolist()
        self.n_features: int = adata.shape[1]
        self.covariate_keys: List[str] = covariate_keys
        self.sampling_method: str = sampling_method
        self.verbose: bool = verbose

        # copy the expression matrix
        # owing to ALPINE takes the sample of feature by samples
        # we do need to transport the matrix X
        X: Float32Array = copy(adata.X).astype(np.float32).T  # type: ignore
        n_sample = X.shape[1]

        # encode the covariates
        self.fe: FeatureEncoders = FeatureEncoders(covariate_keys)
        Y: List[Float32Array] = self.fe.fit_transform(adata.obs)  # type: ignore

        # if batch_size is none -> the batch_size = number of samples
        self.batch_size: int = batch_size if batch_size is not None else n_sample

        # if max_iter is not define, the ALPINE will run a warm-up run that to search the max_iter
        # based on the elbow point of the reconstruction error
        if max_iter is None:
            # initialize a warm-up AlpineMatrices
            m_warmup: AlpineMatrices = self._initialize_matrices(X, Y)
            self.max_iter: int = 200
            self._fit(m_warmup)
            self.max_iter = self._compute_best_iter(
                self.loss_history["reconstruction loss"].values
            )

            # release the memory used by the warm-up run
            del m_warmup
            gc.collect()
            if self.device.type == "cuda":
                torch.cuda.empty_cache()
        else:
            self.max_iter: int = max_iter

        # main training loop
        # initialize other matrices
        m: AlpineMatrices = self._initialize_matrices(X, Y)
        self._fit(m)

        # if scaling is needed
        if self.scale_needed:
            self._scale_matrices(m)

        # only when fit complete the function will generate matrices attributes
        self.matrices: Dict[str, Union[Float32Array, List[Float32Array]]] = m.to_numpy()

        # automatically save the embedding into anndata
        self.store_embeddings(adata)
        return self

    def transform(
        self,
        adata: ad.AnnData,
        n_iter: Optional[int] = None,
    ) -> None:
        # Check if the model has been trained
        if not hasattr(self, "matrices"):
            raise RuntimeError("Model is not trained yet. Please fit the model first.")

        if not isinstance(adata, ad.AnnData):
            raise TypeError("adata must be an AnnData object.")

        if not isinstance(n_iter, (int, type(None))) or (
            n_iter is not None and n_iter <= 0
        ):
            raise ValueError("n_iter must be a positive integer or None.")

        n_iter = n_iter if n_iter is not None else self.max_iter
        self._transform(adata, n_iter)

    def fit_transform(
        self,
        adata: ad.AnnData,
        covariate_keys: List[str],
        batch_size: Optional[int] = None,
        max_iter: Optional[int] = None,
        sampling_method: str = "random",
        verbose: bool = False,
    ) -> None:
        self.fit(
            adata,
            covariate_keys,
            batch_size=batch_size,
            max_iter=max_iter,
            sampling_method=sampling_method,
            verbose=verbose,
        ).transform(adata)

    def compute_loss(self, adata: ad.AnnData):
        if not hasattr(self, "matrices"):
            raise RuntimeError("Model is not trained yet. Please fit the model first.")

        if not isinstance(adata, ad.AnnData):
            raise TypeError("adata must be an AnnData object.")

        if "ALPINE_embedding" not in adata.obsm:
            raise ValueError(
                "ALPINE_embedding not found in adata.obsm. Please transform the data first."
            )

        def kl_divergence(y, y_hat):
            y_hat = np.clip(y_hat, a_min=self.eps, a_max=None)
            return np.sum(
                y * np.log(np.clip(y / y_hat, a_min=self.eps, a_max=None)) - y + y_hat
            )

        X = copy(adata.X).astype(np.float32).T  # type: ignore

        # get Ws and Hs
        Ws, Hs = [], []
        for _, covariate in enumerate(self.covariate_keys):
            Hs.append(copy(adata.obsm[covariate].T))
            Ws.append(copy(adata.varm[covariate]))
        Hs.append(copy(adata.obsm["ALPINE_embedding"].T))
        Ws.append(copy(adata.varm["ALPINE_weights"]))

        # compute the reconstruction loss
        W = np.concatenate(Ws, axis=1)
        H = np.concatenate(Hs, axis=0)
        recon_loss = np.linalg.norm(X - W @ H, ord="fro") ** 2

        # label matrices
        Ys = self.fe.transform(adata.obs)  # type: ignore
        Bs = self.matrices["Bs"]

        # compute the prediction loss
        if self.loss_type == "kl-divergence":
            pred_loss = [kl_divergence(Ys[i].T, Bs[i] @ Hs[i]) for i in range(len(Ys))]
        else:
            pred_loss = [
                np.linalg.norm(Ys[i].T - Bs[i] @ Hs[i], ord="fro") ** 2
                for i in range(len(Ys))
            ]

        total_loss = recon_loss + sum(
            [self.lam[i] * pl for i, pl in enumerate(pred_loss)]
        )
        return total_loss

    def get_decomposed_matrices(
        self,
    ) -> Dict[str, Union[Float32Array, Union[Float32Array, List[Float32Array]]]]:
        if not hasattr(self, "matrices"):
            raise RuntimeError("Model is not trained yet. Please fit the model first.")
        else:
            return self.matrices

    def get_covariate_gene_scores(
            self,
            adata: Optional[ad.AnnData] = None,
        ) -> Union[Dict[str, pd.DataFrame], None]:
        
        if not hasattr(self, "matrices"):
            raise RuntimeError("Model is not trained yet. Please fit the model first.")

        cov_gene_scores = {}
        for i, covariate in enumerate(self.covariate_keys):
            W = self.matrices["Ws"][i]
            H = self.matrices["Hs"][i]
            Y = self.matrices["Ys"][i]

            HY = H @ Y.T / Y.sum(axis=1)
            cond_genes = W @ HY

            colnames = self.fe.encoded_labels[covariate]
            cov_gene_scores[covariate] = pd.DataFrame(
                cond_genes, index=self.feature_names, columns=colnames
            )

        if adata is None:
            return cov_gene_scores
        else:
            for condition, df in cov_gene_scores.items():
                adata.varm[condition + "_gene_scores"] = df
            return None

    def get_normalized_expression(
        self, adata: ad.AnnData, library_size: Optional[float] = None
    ) -> None:
        # the get_normalized_expression will used the transformed ALPINE_embedding to recontruct the counts
        # therefore, need to ensure model is trained and data is transformed
        if not hasattr(self, "matrices"):
            raise RuntimeError("Model is not trained yet. Please fit the model first.")
        elif not isinstance(adata, ad.AnnData):
            raise TypeError("adata must be an AnnData object.")
        elif "ALPINE_embedding" not in adata.obsm:
            raise ValueError(
                "ALPINE_embedding not found in adata.obsm. Please transform the data first."
            )
        elif (library_size is not None) and (library_size <= 0):
            raise ValueError("library_size must be a positive float.")

        # Reconstruct the counts from the learned matrices
        W: Float32Array = self.matrices["Ws"][-1]
        H: Float32Array = adata.obsm["ALPINE_embedding"].T  # type: ignore
        X_normalized = np.dot(W, H).astype(np.float32).T

        # If library_size is not provided, use the default target_sum=None
        temp = ad.AnnData(X_normalized)
        sc.pp.normalize_total(temp, target_sum=library_size)

        # save the normalized_expression into the adata.layers
        adata.layers["normalized_expression"] = copy(temp.X)

    def store_embeddings(self, adata: ad.AnnData) -> None:
        # check whether the model has been trained and validate the input
        if not hasattr(self, "matrices"):
            raise RuntimeError("Model is not trained yet. Please fit the model first.")
        elif not isinstance(adata, ad.AnnData):
            raise TypeError("adata must be an AnnData object.")

        # save the unguided parts
        adata.obsm["ALPINE_embedding"] = copy(self.matrices["Hs"][-1].T)
        adata.varm["ALPINE_weights"] = copy(self.matrices["Ws"][-1])

        dummy_matrices = self.fe.transform(adata.obs)  # type: ignore

        # save the guided parts (covariate assocaited)
        for i, covariate in enumerate(self.covariate_keys):
            adata.obsm[covariate] = copy(self.matrices["Hs"][i].T)
            adata.obsm[f"{covariate}_dummy_matrix"]  = dummy_matrices[i]  # type: ignore
            adata.varm[covariate] = copy(self.matrices["Ws"][i])

    def _validate_init_args(self) -> None:
        # validate n_components
        if self.n_components <= 0:
            raise ValueError("n_components must be greater than 0.")

        # validate n_covariate_components
        if not isinstance(self.n_covariate_components, list):
            raise TypeError("n_covariate_components must be a list.")
        else:
            for n in self.n_covariate_components:
                if not isinstance(n, int) or n < 0:
                    raise ValueError(
                        "Each element in n_covariate_components must be a non-negative integer."
                    )

        # validate lam
        if not isinstance(self.lam, list):
            raise TypeError("lam must be in a list.")
        else:
            for lam in self.lam:
                if not isinstance(lam, float) or lam < 0:
                    raise ValueError(
                        "Each element in lam must be a non-negative float."
                    )

        # validate alpha_W
        if not isinstance(self.alpha_W, float) or self.alpha_W < 0:
            raise ValueError("alpha_W must be a non-negative float.")

        # validate orth_W
        if not isinstance(self.orth_W, float) or self.orth_W < 0:
            raise ValueError("orth_W must be a non-negative float.")

        # validate l1_ratio_W
        if (
            not isinstance(self.l1_ratio_W, float)
            or self.l1_ratio_W < 0
            or self.l1_ratio_W > 1
        ):
            raise ValueError("l1_ratio_W must be a float between 0 and 1.")

        # validate scale_needed
        if not isinstance(self.scale_needed, bool):
            raise TypeError("scale_needed must be a boolean.")

        # validate loss_type
        if not isinstance(self.loss_type, str):
            raise TypeError("loss_type must be a string.")
        else:
            valid_loss_types = ["kl-divergence", "frobenius"]
            if self.loss_type not in valid_loss_types:
                raise ValueError(f"loss_type must be one of {valid_loss_types}.")

        # validate eps
        if not isinstance(self.eps, float) or self.eps < 0:
            raise ValueError("eps must be a non-negative float.")

        # validate random_state
        if not isinstance(self.random_state, int) or self.random_state < 0:
            raise ValueError("random_state must be a non-negative integer.")

    def _validate_fit_args(
        self,
        adata,
        covariate_keys,
        batch_size,
        max_iter,
        sampling_method,
        verbose,
    ) -> None:
        if not isinstance(adata, ad.AnnData):
            raise TypeError("adata must be an AnnData object.")

        if not isinstance(adata.X, np.ndarray):
            raise TypeError("adata.X must be a numpy array.")
        elif adata.X.ndim != 2:
            raise ValueError("adata.X must be a 2D numpy array.")
        elif not np.all(adata.X >= 0):
            raise ValueError("All elements in adata.X must be non-negative.")

        if not isinstance(covariate_keys, list):
            raise TypeError("covariate_keys must be a list.")
        elif not len(covariate_keys) == len(self.n_covariate_components):
            raise ValueError(
                "Length of covariate_keys must match length of n_covariate_components."
            )
        else:
            for key in covariate_keys:
                if not isinstance(key, str):
                    raise TypeError("Each element in covariate_keys must be a string.")

                if key not in adata.obs.columns:
                    raise ValueError(f"Covariate key '{key}' not found in adata.obs.")
                if not adata.obs[key].dtype.kind == "O":
                    raise TypeError(
                        f"Covariate '{key}' in adata.obs must be a categorical or object type variable."
                    )

        if (
            batch_size is not None
            and not isinstance(batch_size, int)
            and batch_size > 0
        ):
            raise TypeError("batch_size must be a positive integer.")

        if max_iter is not None and not isinstance(max_iter, int) and max_iter > 0:
            raise TypeError("max_iter must be a positive integer.")

        if not isinstance(sampling_method, str):
            raise TypeError("sampling_method must be a string.")

        if not isinstance(verbose, bool):
            raise TypeError("verbose must be a boolean.")

    def _initialize_matrices(
        self, X_array: Float32Array, Y_list_array: List[Float32Array]
    ) -> AlpineMatrices:
        # set random seed for cpu and gpu
        torch.manual_seed(self.random_state)
        if self.device.type == "cuda":
            torch.cuda.manual_seed(self.random_state)

        # convert numpy arrays to torch tensors
        X: torch.Tensor = torch.tensor(X_array, dtype=torch.float32, device=self.device)
        Y: List[torch.Tensor] = [
            torch.tensor(y.T, dtype=torch.float32, device=self.device)
            for y in Y_list_array
        ]

        n_features, n_samples = X.shape

        # random initialize the Ws, Hs, and Bs
        Ws: List[torch.Tensor] = [
            torch.rand((n_features, k), dtype=torch.float32, device=self.device)
            for k in self.n_all_components
        ]
        Ws = [w.clamp(min=self.eps) for w in Ws]

        Hs: List[torch.Tensor] = [
            torch.rand((k, n_samples), dtype=torch.float32, device=self.device)
            for k in self.n_all_components
        ]
        Hs = [h.clamp(min=self.eps) for h in Hs]

        Bs: List[torch.Tensor] = [
            torch.rand((y.shape[0], k), dtype=torch.float32, device=self.device)
            for (y, k) in zip(Y, self.n_covariate_components)
        ]
        Bs = [b.clamp(min=self.eps) for b in Bs]

        return AlpineMatrices(X, Y, Ws, Hs, Bs)

    def _compute_orthogonal_matrix(self, size: int) -> torch.Tensor:
        # orthogonal matrix
        orth_mat: torch.Tensor = torch.ones(
            (size, size),
            dtype=torch.float32,
            device=self.device,
        )
        orth_mat -= torch.eye(size, dtype=torch.float32, device=self.device)
        orth_mat = self.orth_W * orth_mat

        return orth_mat

    def _fit(self, m: AlpineMatrices) -> None:
        loss_history: List[List[float]] = []

        # setting progress bar
        pbar = (
            tqdm(total=self.max_iter, desc="Iteration", ncols=100)
            if self.verbose
            else nullcontext()
        )

        joint_labels = create_joint_labels_from_dummy_matrices(m.Ys)

        with torch.no_grad():
            with pbar:
                for i_iter in range(self.max_iter):
                    # Generate indices for the entire epoch
                    epoch_indices = generate_epoch_indices(
                        joint_labels=joint_labels,
                        sampling_method=self.sampling_method,
                        device=self.device,
                    )

                    # calculate how many batches needed
                    num_batches = get_num_batches(len(epoch_indices), self.batch_size)

                    # iterate over all batches
                    for batch_num in range(num_batches):
                        batch_indices = get_batch_indices(
                            epoch_indices, batch_num, self.batch_size
                        )
                        if len(batch_indices) == 0:
                            break

                        # Precompute reused views
                        X_batch = m.X[:, batch_indices]
                        Ys_batch = [Y[:, batch_indices] for Y in m.Ys]

                        if self.use_als:
                            # ALS updates inlined
                            for idx in range(len(self.n_all_components)):
                                # === Update W[idx] ===
                                Hs_batch = [h[:, batch_indices] for h in m.Hs]
                                H_batch = Hs_batch[idx]
                                W = m.Ws[idx]
                                W_cat = torch.cat(m.Ws, dim=1)
                                H_cat_batch = torch.cat(Hs_batch, dim=0)

                                numerator = 2 * X_batch @ H_batch.T
                                eye_mat = torch.eye(
                                    W.shape[1], dtype=torch.float32, device=self.device
                                )
                                orth_mat = self._compute_orthogonal_matrix(W.shape[1])
                                denominator = (
                                    2 * W_cat @ H_cat_batch @ H_batch.T
                                    + (1 - self.l1_ratio_W) * self.alpha_W * W @ eye_mat
                                    + W @ orth_mat
                                )
                                denominator += self.l1_ratio_W * self.alpha_W * torch.ones_like(denominator)
                                denominator = torch.clamp(denominator, min=self.eps)
                                m.Ws[idx] *= numerator / denominator

                                # === Update B[idx] if covariate ===
                                if idx < len(self.n_covariate_components):
                                    Y_batch = Ys_batch[idx]
                                    B = m.Bs[idx]
                                    if self.loss_type == "kl-divergence":
                                        numerator = (
                                            self.lam[idx]
                                            * (Y_batch / torch.clamp(B @ H_batch, min=self.eps))
                                            @ H_batch.T
                                        )
                                        denominator = self.lam[idx] * torch.ones_like(Y_batch) @ H_batch.T
                                    else:
                                        numerator = 2 * Y_batch @ H_batch.T
                                        denominator = 2 * B @ H_batch @ H_batch.T
                                    denominator = torch.clamp(denominator, min=self.eps)
                                    m.Bs[idx] *= numerator / denominator

                                # === Update H[idx] ===
                                W = m.Ws[idx]
                                W_cat = torch.cat(m.Ws, dim=1)
                                unguided_num = 2 * W.T @ X_batch
                                unguided_den = 2 * W.T @ (W_cat @ H_cat_batch)

                                if idx < len(self.covariate_keys):
                                    Y_batch = Ys_batch[idx]
                                    B = m.Bs[idx]
                                    if self.loss_type == "kl-divergence":
                                        guided_num = (
                                            self.lam[idx]
                                            * B.T @ (Y_batch / torch.clamp(B @ H_batch, min=self.eps))
                                        )
                                        guided_den = self.lam[idx] * B.T @ torch.ones_like(Y_batch)
                                    else:
                                        guided_num = 2 * self.lam[idx] * B.T @ Y_batch
                                        guided_den = 2 * self.lam[idx] * B.T @ (B @ H_batch)
                                    numerator = unguided_num + guided_num
                                    denominator = unguided_den + guided_den
                                    denominator = torch.clamp(denominator, min=self.eps)
                                    m.Hs[idx][:, batch_indices] *= numerator / denominator
                                else:
                                    unguided_den = torch.clamp(unguided_den, min=self.eps)
                                    m.Hs[idx][:, batch_indices] *= unguided_num / unguided_den
                        else:
                            # Multiplicative updates (non-ALS), inlined
                            # === Update W ===
                            W_cat = torch.cat(m.Ws, dim=1)
                            Hs_batch = [h[:, batch_indices] for h in m.Hs]
                            H_cat_batch = torch.cat(Hs_batch, dim=0)

                            numerator = 2 * X_batch @ H_cat_batch.T
                            orth_mat = self._compute_orthogonal_matrix(W_cat.shape[1])
                            denominator = (
                                2 * W_cat @ H_cat_batch @ H_cat_batch.T
                                + (1 - self.l1_ratio_W) * self.alpha_W * W_cat
                                + W_cat @ orth_mat
                            )
                            denominator += self.l1_ratio_W * self.alpha_W * torch.ones_like(denominator)
                            denominator = torch.clamp(denominator, min=self.eps)
                            W_cat *= numerator / denominator

                            # split back into m.Ws
                            start = 0
                            for idx, w in enumerate(m.Ws):
                                end = start + w.shape[1]
                                m.Ws[idx] = W_cat[:, start:end]
                                start = end

                            # === Update Bs ===
                            for i in range(len(self.covariate_keys)):
                                Yb, Hb, B = Ys_batch[i], Hs_batch[i], m.Bs[i]
                                if self.loss_type == "kl-divergence":
                                    numerator = (
                                        self.lam[i]
                                        * (Yb / torch.clamp(B @ Hb, min=self.eps))
                                        @ Hb.T
                                    )
                                    denominator = self.lam[i] * torch.ones_like(Yb) @ Hb.T
                                else:
                                    numerator = 2 * Yb @ Hb.T
                                    denominator = 2 * B @ Hb @ Hb.T
                                denominator = torch.clamp(denominator, min=self.eps)
                                m.Bs[i] *= numerator / denominator

                            # === Update H ===
                            W_cat = torch.cat(m.Ws, dim=1)
                            numerator = torch.zeros_like(H_cat_batch)
                            denominator = torch.zeros_like(H_cat_batch)

                            # prediction part
                            start = 0
                            for i in range(len(self.covariate_keys)):
                                end = start + Hs_batch[i].shape[0]
                                if self.loss_type == "kl-divergence":
                                    guided_num = (
                                        self.lam[i]
                                        * m.Bs[i].T @ (Ys_batch[i] / torch.clamp(m.Bs[i] @ Hs_batch[i], min=self.eps))
                                    )
                                    guided_den = self.lam[i] * m.Bs[i].T @ torch.ones_like(Ys_batch[i])
                                else:
                                    guided_num = 2 * self.lam[i] * m.Bs[i].T @ Ys_batch[i]
                                    guided_den = 2 * self.lam[i] * m.Bs[i].T @ (m.Bs[i] @ Hs_batch[i])
                                numerator[start:end] = guided_num
                                denominator[start:end] = guided_den
                                start = end

                            # reconstruction part
                            numerator += 2 * W_cat.T @ X_batch
                            denominator += 2 * W_cat.T @ (W_cat @ H_cat_batch)
                            denominator = torch.clamp(denominator, min=self.eps)
                            H_cat_batch *= numerator / denominator

                            # split back into m.Hs
                            start = 0
                            for j, H in enumerate(Hs_batch):
                                end = start + H.shape[0]
                                m.Hs[j][:, batch_indices] = H_cat_batch[start:end]
                                start = end

                    # compute loss
                    loss = self._compute_loss(m)
                    loss_history.append(loss)

                    if self.verbose:
                        pbar.set_postfix({"objective loss": loss[0]})  # type: ignore
                        pbar.update(1)  # type: ignore

                colnames = ["total loss", "reconstruction loss"] + [
                    f"prediction loss({k})" for k in self.covariate_keys
                ]
                self.loss_history = pd.DataFrame(loss_history, columns=colnames)

    def _transform(self, adata: ad.AnnData, n_iter: int) -> None:
        # validate the input and convert to torch.Tensor
        X_array: Float32Array = copy(adata.X).astype(np.float32).T  # type: ignore
        if not np.all(X_array >= 0):
            raise ValueError("All elements in adata.X must be non-negative.")
        X: torch.Tensor = torch.tensor(X_array, dtype=torch.float32, device=self.device)
        n_sample: int = X.shape[1]

        # Initialize H_transformed
        H_transformed: torch.Tensor = torch.rand(
            (self.total_components, n_sample), dtype=torch.float32, device=self.device
        )
        Hs_transformed = [
            torch.zeros((k, n_sample), dtype=torch.float32, device=self.device)
            for k in self.n_all_components
        ]

        # convert np.ndarray to torch.Tensor
        W: torch.Tensor = torch.cat(
            [
                torch.tensor(w, dtype=torch.float32, device=self.device)
                for w in self.matrices["Ws"]
            ],
            dim=1,
        )

        # main transform loop (basic multiplicative update for H)
        for _ in range(n_iter):
            numerator = 2 * W.T @ X
            denominator = 2 * W.T @ (W @ H_transformed)
            denominator = torch.clamp(denominator, min=self.eps)
            H_transformed *= numerator / denominator

        # seperate entire H into individual part
        start_idx, end_idx = 0, 0
        for idx in range(len(Hs_transformed)):
            end_idx = start_idx + Hs_transformed[idx].shape[0]
            Hs_transformed[idx] = H_transformed[start_idx:end_idx]
            start_idx = end_idx

        # save the Hs_transformed_array into the adata
        Hs_transformed_array = [H.cpu().numpy() for H in Hs_transformed]
        for i, covariate in enumerate(self.covariate_keys):
            adata.obsm[covariate] = Hs_transformed_array[i].T
            adata.varm[covariate] = deepcopy(self.matrices["Ws"][i])
        adata.obsm["ALPINE_embedding"] = Hs_transformed_array[-1].T
        adata.varm["ALPINE_weights"] = deepcopy(self.matrices["Ws"][-1])

    def _compute_loss(self, m) -> List[float]:
        def kl_divergence(y, y_hat):
            y_hat = torch.clamp(y_hat, min=self.eps)
            return torch.sum(
                y * torch.log(torch.clamp(y / y_hat, min=self.eps)) - y + y_hat
            ).item()

        # compute the reconstruction loss
        W = torch.cat(m.Ws, dim=1)
        H = torch.cat(m.Hs, dim=0)
        recon_loss = torch.norm(m.X - W @ H, p="fro") ** 2
        recon_loss = recon_loss.item()

        # compute the prediction loss
        if self.loss_type == "kl-divergence":
            pred_loss = [
                kl_divergence(m.Ys[i], m.Bs[i] @ m.Hs[i]) for i in range(len(m.Ys))
            ]
        else:
            pred_loss = [
                (torch.norm(m.Ys[i] - m.Bs[i] @ m.Hs[i], p="fro") ** 2).item()
                for i in range(len(m.Ys))
            ]

        total_loss = recon_loss + sum(
            [self.lam[i] * pl for i, pl in enumerate(pred_loss)]
        )
        return [total_loss, recon_loss] + pred_loss

    def _compute_best_iter(self, train_loss) -> int:
        # compute the best iteration based on the warm-up run
        # using the Kneedle algorithm to find the elbow point
        kneedle = KneeLocator(
            np.arange(0, len(train_loss)),
            np.log10(train_loss),
            curve="convex",
            direction="decreasing",
            interp_method="polynomial",
            polynomial_degree=2
        )
        if kneedle.elbow is not None:
            return int(kneedle.elbow)
        else:
            warnings.warn("Kneedle elbow not found, using default max_iter=200")
            return 200

    def _scale_matrices(self, m: AlpineMatrices) -> None:
        # normalize each signature in W to sum to 1
        # the H and B are scaled based on the w_scaler
        for i in range(len(m.Ws)):
            w_scaler: torch.Tensor = m.Ws[i].sum(dim=0)
            m.Ws[i] = m.Ws[i] / w_scaler
            m.Hs[i] = m.Hs[i] * w_scaler.unsqueeze(1)

            if i < len(self.n_covariate_components):
                m.Bs[i] = m.Bs[i] / w_scaler
```
