## Supplemental Code for "Interpretable phenotype decoding from multi-condition sequencing data with ALPINE": optimization.py

```
import pickle
import scanpy as sc
import numpy as np
import pandas as pd
import anndata as ad

from copy import copy
from typing import Tuple, List, Optional
from sklearn.metrics.cluster import adjusted_rand_score, homogeneity_score
from hyperopt import fmin, tpe, hp, Trials, STATUS_OK, STATUS_FAIL
from .main import ALPINE
from sklearn.model_selection import StratifiedKFold


class ComponentOptimizer:
    def __init__(
        self,
        adata: ad.AnnData,
        covariate_keys: list[str],
        use_als: bool = False,
        loss_type: str = "kl-divergence",
        max_iter: int | None = None,
        batch_size: int | None = None,
        sampling_method: str = "random",
        device: str = "cuda",
        random_state: int = 42,
    ):
        self._validate_init_args(
            adata, covariate_keys, loss_type, max_iter, batch_size, device, random_state
        )

        self.adata: ad.Anndata = adata.copy()
        self.covariate_keys: List[str] = covariate_keys
        self.use_als: bool = use_als
        self.loss_type: str = loss_type
        self.max_iter: Optional[int] = max_iter  # type: ignore
        self.batch_size: Optional[int] = batch_size
        self.sampling_method: str = sampling_method
        self.device: str = device
        self.random_state: int = random_state
        self.best_param: dict = {}

        if self.max_iter is None:
            print(
                f"Owing to max_iter being None, it will be determine by the average of the first n_splits iterations."
            )
            self.max_iter_detect = True
        else:
            self.max_iter_detect = False

    def search_hyperparams(
        self,
        n_total_components_range: Tuple[int, int] = (10, 100),
        lam_range: Tuple[float, float] = (1.0, 1e4),
        orth_W_range: Tuple[float, float] = (0.0, 1.0),
        alpha_W_range: Tuple[float, float] = (0.0, 100.0),
        l1_ratio_W_range: Tuple[float, float] = (0.0, 1.0),
        min_covariate_components: Optional[List[int]] = None,
        n_splits: int = 3,
        max_evals: int = 100,
        trials_filename: Optional[str] = None,
    ):
        self._validate_search_args(
            n_total_components_range,
            lam_range,
            orth_W_range,
            alpha_W_range,
            l1_ratio_W_range,
            n_splits,
            max_evals,
        )

        self.iter_records: List = []
        self.n_splits: int = n_splits

        # Load previous trials if specified
        if trials_filename is not None:
            self.load_trials(trials_filename)
        else:
            self.trials = Trials()  # Create a new trials object if not loading

        # check min_covariate_components
        if min_covariate_components is None:
            self.min_covariate_components = [self.adata.obs[key].nunique() for key in self.covariate_keys]
        else:
            if isinstance(min_covariate_components, list):
                if len(min_covariate_components) != len(self.covariate_keys):
                    raise ValueError("min_covariate_components should have the same length as the number of covariates.")
        
            if any(comp < 2 for comp in min_covariate_components):
                raise ValueError("min_covariate_components should be greater than or equal to 2.")
            self.min_covariate_components = min_covariate_components
        
        # Define the search space for Bayesian Optimization
        self.space = {
            # unguided component size
            "n_total_components": hp.quniform(
                "n_total_components",
                n_total_components_range[0],
                n_total_components_range[1],
                1,
            ),
            # orthogonal
            "orth_W": hp.uniform("orth_W", orth_W_range[0], orth_W_range[1]),
            # alpha
            "alpha_W": hp.uniform("alpha_W", alpha_W_range[0], alpha_W_range[1]),
            # l1
            "l1_ratio_W": hp.uniform(
                "l1_ratio_W", l1_ratio_W_range[0], l1_ratio_W_range[1]
            ),
            "splits": [
                hp.uniform(f"split_{i}", 0, 1) for i in range(len(self.covariate_keys) + 1)
            ],
        }

        # Distribute the remaining space across covariate components
        for i in range(len(self.covariate_keys)):
            self.space[f"lam_{i}"] = hp.qloguniform(
                f"lam_{i}", np.log(lam_range[0]), np.log(lam_range[1]), 1
            )

        # Run the optimization using TPE
        best = fmin(
            self.objective,
            self.space,
            algo=tpe.suggest,
            max_evals=max_evals + len(self.trials.trials),
            trials=self.trials,
            rstate=np.random.default_rng(self.random_state)
        )

        if best is None:
            raise RuntimeError("Hyperparameter optimization did not return any result.")
        
        component_params = {
            "n_total_components": best["n_total_components"],
            "splits": [best[f"split_{i}"] for i in range(len(self.covariate_keys) + 1)]
        }
        n_components, n_covariate_components = self._distribute_components(component_params)

        self.best_param["n_components"] = n_components
        self.best_param["n_covariate_components"] = n_covariate_components
        self.best_param["lam"] = [
            float(best[f"lam_{i}"]) for i in range(len(self.covariate_keys))
        ]
        self.best_param["alpha_W"] = best["alpha_W"]
        self.best_param["orth_W"] = best["orth_W"]
        self.best_param["l1_ratio_W"] = best["l1_ratio_W"]
        self.best_param["random_state"] = self.random_state

        return self.best_param

    def _distribute_components(self, space):
        total_components = int(space["n_total_components"])
        split_ratios = space["splits"]

        # ensure split_ratios is a list of floats of length num_covariates
        splits = [float(s) for s in split_ratios]

        # normalize splits to sum to 1
        normalized_splits = np.array(splits) / np.sum(splits)

        # allocate components
        n_components = int(total_components / 2)
        rest_components = total_components - n_components

        n_covariate_components = [
            int(round(rest_components * ratio)) for ratio in normalized_splits[:-1]
        ]

        n_covariate_components = [max(self.min_covariate_components[i], n) for i, n in enumerate(n_covariate_components)]
        # last entry is for unguided components, rest are for guided components
        total_covariate_components = sum(n_covariate_components)
        n_components = total_components - total_covariate_components

        return n_components, n_covariate_components

    def objective(self, space):
        # Handle multiple covariate components separately
        lam = [space[f"lam_{i}"] for i in range(len(self.covariate_keys))]

        n_components, n_covariate_components = self._distribute_components(space)

        # Check if the distribution is valid
        cond_1 = sum(n_covariate_components) <= n_components
        cond_2 = all(n >= 2 for n in n_covariate_components)

        if cond_1 and cond_2:
            args = {
                "n_components": n_components,
                "n_covariate_components": n_covariate_components,
                "lam": lam,
                "orth_W": space["orth_W"],
                "alpha_W": space["alpha_W"],
                "l1_ratio_W": space["l1_ratio_W"],
            }

            score = self.calc_score(args)

            trial_history = {
                "n_components": n_components,
                "n_covariate_components": n_covariate_components,
                "lam": [cov_lam for cov_lam in lam],
                "orth_W": space["orth_W"],
                "alpha_W": space["alpha_W"],
                "l1_ratio_W": space["l1_ratio_W"],
                "max_iter": self.iter_records[-1] if self.max_iter_detect else self.max_iter,
                "score": score,
            }

            if self.max_iter_detect:
                if len(self.iter_records) >= self.n_splits:
                    self.max_iter: int = int(
                        sum(self.iter_records) / len(self.iter_records)
                    )
            return {"loss": score, "status": STATUS_OK, "params": trial_history}
        else:
            return {"loss": np.inf, "status": STATUS_FAIL}

    def calc_score(self, args):
        n_covariate_components = args["n_covariate_components"]
        n_components = args["n_components"]
        lam = args["lam"]
        orth_W = args["orth_W"]
        alpha_W = args["alpha_W"]
        l1_ratio_W = args["l1_ratio_W"]

        # Create joint labels for stratification
        if len(self.covariate_keys) == 1:
            joint_labels = self.adata.obs[self.covariate_keys[0]].astype(str)
        else:
            joint_labels = self.adata.obs[self.covariate_keys[0]].astype(str)
            for key in self.covariate_keys[1:]:
                joint_labels = joint_labels + "_" + self.adata.obs[key].astype(str)

        scores = []

        skf = StratifiedKFold(
            n_splits=self.n_splits, shuffle=True, random_state=self.random_state
        )
        for train_idx, val_idx in skf.split(self.adata.X, joint_labels):
            train_adata = self.adata[train_idx].copy()
            val_adata = self.adata[val_idx].copy()

            model = ALPINE(
                n_covariate_components=n_covariate_components,
                n_components=n_components,
                lam=lam,
                orth_W=orth_W,
                alpha_W=alpha_W,
                l1_ratio_W=l1_ratio_W,
                use_als=self.use_als,
                random_state=self.random_state,
                loss_type=self.loss_type,
                device=self.device,
            )

            model.fit(
                adata=train_adata,
                covariate_keys=self.covariate_keys,
                max_iter=self.max_iter,
                batch_size=self.batch_size,
                sampling_method=self.sampling_method,
                verbose=False,
            )

            model.store_embeddings(train_adata)
            _ = model.transform(val_adata)

            # embedding score
            sc.pp.neighbors(val_adata, use_rep='ALPINE_embedding')
            sc.tl.leiden(val_adata, flavor="igraph", resolution=1)
            embedding_score = 0
            for key in self.covariate_keys:
                selected_idx = ~val_adata.obs[key].isna()
                embedding_score += adjusted_rand_score(val_adata.obs[key][selected_idx], val_adata.obs["leiden"][selected_idx])
                embedding_score += homogeneity_score(val_adata.obs[key][selected_idx], val_adata.obs["leiden"][selected_idx])
            embedding_score /= len(self.covariate_keys)

            scores.append(embedding_score)

            if self.max_iter_detect:
                self.iter_records.append(model.max_iter)

        score = np.mean(scores)

        return score

    def extend_training(self, extra_evals=50):
        """
        Continue the Bayesian optimization process with more evaluations.

        Parameters:
            extra_evals: int
                The number of additional evaluations to perform during the Bayesian search.

        Returns:
            dict
                The updated best parameters after the additional evaluations.
        """
        if not hasattr(self, "trials"):
            raise RuntimeError(
                "Please run bayesian_search() before extending training."
            )

        # Update the max_evals for continuing the optimization
        best = fmin(
            fn=self.objective,
            space=self.space,
            algo=tpe.suggest,
            max_evals=len(self.trials.trials)
            + extra_evals,  # Continue from the current trial count
            trials=self.trials,
            rstate=np.random.default_rng(self.random_state)
        )

        if best is None:
            raise RuntimeError("Hyperparameter optimization did not return any result.")

        n_components, n_covariate_components = self._distribute_components(best)

        # Assign the best parameters from the latest search
        self.best_param["n_components"] = n_components
        self.best_param["n_covariate_components"] = n_covariate_components
        self.best_param["lam"] = [
            float(best[f"lam_{i}"]) for i in range(len(self.covariate_keys))
        ]
        self.best_param["orth_W"] = best["orth_W"]
        self.best_param["alpha_W"] = best["alpha_W"]
        self.best_param["l1_ratio_W"] = best["l1_ratio_W"]
        self.best_param["random_state"] = self.random_state

        return copy(self.best_param)

    def save_trials(self, filename: str):
        """
        Save the current trials to a file.

        Parameters:
            filename: str
                Path to the file where the trials will be saved.
        """
        with open(filename, "wb") as f:
            pickle.dump(self.trials, f)
        print(f"Trials saved to {filename}")

    def load_trials(self, filename: str):
        """
        Load the trials from a file.

        Parameters:
            filename: str
                Path to the file from which to load the trials.
        """
        with open(filename, "rb") as f:
            self.trials = pickle.load(f)
        print(f"Trials loaded from {filename}")

    def get_hyperparameter(self, idx):
        """
        Retrieve the hyperparameters of a specific trial based on its index in the sorted training history.
        Args:
            idx (int): The index of the trial in the sorted training history.
        Returns:
            dict: A dictionary containing the hyperparameters of the specified trial.
        Raises:
            IndexError: If the provided index is out of bounds for the training history.
            KeyError: If the expected keys ('tid' or 'params') are missing in the trial data.
        Notes:
            - The training history is assumed to be sorted prior to fetching the trial.
            - The method matches the trial in `self.trials` using the trial ID ('tid') from the history.
        """
        # Fetch the sorted trial history
        history_df = self.get_train_history()

        # Retrieve the corresponding trial from sorted index
        trial = history_df.iloc[idx]

        # Find the corresponding trial in self.trials using the 'tid' (trial ID)
        trial_tid = trial["tid"]

        # Extract the trial's hyperparameters using the 'tid'
        for trial in self.trials.trials:
            if trial["tid"] == trial_tid:
                return trial["result"]["params"]

    def get_train_history(self):
        """
        Retrieves and processes the training history from the optimization trials.

        This method extracts trial information, processes it into a structured format,
        and returns a pandas DataFrame containing details about each trial, including
        the parameters, scores, and derived metrics.

        Returns:
            pd.DataFrame: A DataFrame containing the following columns:
                - n_components: The number of components used in the trial.
                - n_covariate_components_X: Expanded columns for each covariate component (X is the index).
                - n_total_components: The total number of components, calculated as the sum of
                  `n_components` and all `n_covariate_components`.
                - score: The score (loss) of the trial.
                - tid: The trial ID for tracking.
                - Other columns corresponding to the trial parameters.

        Notes:
            - The DataFrame is sorted by the `score` column in descending order.
            - The index of the DataFrame is reset after sorting.
            - The columns are reordered to place `n_components` and related columns at the beginning.
        """
        history = []

        # Extract trial information
        for trial in self.trials.trials:
            if "result" in trial and trial["result"]["status"] == STATUS_OK:
                trial_info = trial["result"]["params"]
                trial_info["score"] = trial["result"]["loss"]
                trial_info["tid"] = trial["tid"]  # Add trial ID for tracking
                history.append(trial_info)

        # Convert to pandas DataFrame
        history_df = pd.DataFrame(history)

        # Expand the list columns into separate columns
        n_covariate_df = pd.DataFrame(
            history_df["n_covariate_components"].tolist(),
            columns=[
                f"n_covariate_components_{i}"
                for i in range(len(history_df["n_covariate_components"].iloc[0]))
            ],
        )
        lam_df = pd.DataFrame(
            history_df["lam"].tolist(),
            columns=[f"lam_{i}" for i in range(len(history_df["lam"].iloc[0]))],
        )

        # Concatenate the expanded columns back to the original DataFrame
        history_df = pd.concat(
            [
                history_df.drop(columns=["n_covariate_components", "lam"]),
                n_covariate_df,
                lam_df,
            ],
            axis=1,
        )

        # Add n_total_components as the sum of n_components and all n_covariate_components
        history_df["n_total_components"] = history_df["n_components"] + history_df[
            [f"n_covariate_components_{i}" for i in range(len(n_covariate_df.columns))]
        ].sum(axis=1)

        # Reorder columns to have n_components related columns at the beginning
        columns_order = (
            ["n_components"]
            + [
                f"n_covariate_components_{i}"
                for i in range(len(n_covariate_df.columns))
            ]
            + ["n_total_components"]
            + [
                col
                for col in history_df.columns
                if col
                not in ["n_components", "n_total_components"]
                + [
                    f"n_covariate_components_{i}"
                    for i in range(len(n_covariate_df.columns))
                ]
            ]
        )
        history_df = history_df[columns_order]

        # Sort by score in descending order and reset the index
        history_df = history_df.sort_values(by="score", ascending=False).reset_index(
            drop=True
        )

        return history_df

    def fit_the_best_param(self):
        """
        Fits the ALPINE model using the best parameters found through Bayesian search.
        This method initializes an ALPINE model with the best parameters stored in `self.best_param`
        and fits it using the provided data and configuration.
        Returns:
            ALPINE: An instance of the ALPINE model fitted with the best parameters.
        Raises:
            RuntimeError: If the `best_param` attribute is not set. Ensure that `bayesian_search()`
                          is run prior to calling this method to determine the best parameters.
        """

        if not hasattr(self, "best_param"):
            raise RuntimeError(
                "Please run bayesian_search() to find the best parameters first."
            )

        model = ALPINE(
            **self.best_param,
            use_als=self.use_als,
            random_state=self.random_state,
            loss_type=self.loss_type,
            device=self.device,
        )
        model.fit(
            adata=self.adata,
            covariate_keys=self.covariate_keys,
            max_iter=self.max_iter,
            batch_size=self.batch_size,
            verbose=False,
        )
        return model

    def _validate_init_args(
        self,
        adata,
        covariate_keys,
        loss_type,
        max_iter,
        batch_size,
        device,
        random_state,
    ) -> None:
        # validate the input type
        if not isinstance(adata, ad.AnnData):
            raise TypeError("adata must be an instance of AnnData")

        # validate the covariate_keys
        if not isinstance(covariate_keys, list):
            raise TypeError("covariate_keys must be a list")

        if not all(isinstance(key, str) for key in covariate_keys):
            raise TypeError("All covariate_keys must be strings")

        if not all(key in adata.obs.columns for key in covariate_keys):
            raise ValueError("All covariate_keys must be present in adata.obs")

        # validate the loss type
        if loss_type not in ["kl-divergence", "frobenius"]:
            raise ValueError("loss_type must be either 'kl-divergence' or 'frobenius'")

        # validate other parameters
        if max_iter is not None:
            if not isinstance(max_iter, int) or max_iter < 0:
                raise ValueError("max_iter must be a non-negative integer")

        if batch_size is not None:
            if not isinstance(batch_size, int) or batch_size < 0:
                raise ValueError("batch_size must be a non-negative integer")

        if not isinstance(random_state, int):
            raise TypeError("random_state must be an integer")

    def _validate_search_args(
        self,
        n_total_components_range: Tuple[int, int],
        lam_range: Tuple[float, float],
        orth_W_range: Tuple[float, float],
        alpha_W_range: Tuple[float, float],
        l1_ratio_W_range: Tuple[float, float],
        n_splits: int,
        max_evals: int,
    ) -> None:
        # validate the n_total_components_range
        if (
            not isinstance(n_total_components_range, tuple)
            or len(n_total_components_range) != 2
        ):
            raise TypeError("n_total_components_range must be a tuple of two integers")
        else:
            if n_total_components_range[0] >= n_total_components_range[1]:
                raise ValueError(
                    "n_total_components_range must be a tuple with the first element less than the second"
                )
            if n_total_components_range[0] < 2:
                raise ValueError(
                    "n_total_components_range must be a tuple with the first element greater than or equal to 2"
                )

        def _validate_tuple_range(arg, name, dtype=float):
            if not isinstance(arg, tuple) or len(arg) != 2:
                raise TypeError(f"{name} must be a tuple of two {dtype.__name__}s")
            if not all(isinstance(x, dtype) for x in arg):
                raise TypeError(f"All elements of {name} must be {dtype.__name__}s")
            if arg[0] >= arg[1]:
                raise ValueError(
                    f"{name} must be a tuple with the first element less than the second"
                )

        _validate_tuple_range(lam_range, "lam_range", float)
        _validate_tuple_range(orth_W_range, "orth_W_range", float)
        _validate_tuple_range(alpha_W_range, "alpha_W_range", float)

        _validate_tuple_range(l1_ratio_W_range, "l1_ratio_W_range", float)
        if l1_ratio_W_range[1] > 1.0:
            raise ValueError(
                "l1_ratio_W_range's second element must be less than or equal to 1.0"
            )

        if not isinstance(n_splits, int):
            raise TypeError("n_splits must be an integer")
        if n_splits < 2:
            raise ValueError("n_splits must be greater than or equal to 2")

        if not isinstance(max_evals, int) or max_evals <= 0:
            raise ValueError("max_evals must be a positive integer")
```
