## Supplemental Code for "Interpretable phenotype decoding from multi-condition sequencing data with ALPINE": encoder.py

```
import numpy as np
import pandas as pd
import numpy.typing as npt

from sklearn.preprocessing import OneHotEncoder
from typing import List, Dict

Float32Array = npt.NDArray[np.float32]


class FeatureEncoders:
    def __init__(self, covariate_keys: List[str]):
        self.covariate_keys: List[str] = covariate_keys
        self.encoders: Dict[str, OneHotEncoder] = {}
        self.encoded_labels: Dict[str, List[str]] = {}

    def fit_transform(self, df: pd.DataFrame) -> List[Float32Array]:
        if not isinstance(df, pd.DataFrame):
            raise TypeError("adata.obs must be a pandas DataFrame.")

        transformed_matrices: List[Float32Array] = []
        for key in self.covariate_keys:
            encoder: OneHotEncoder = OneHotEncoder(
                sparse_output=False, handle_unknown="ignore"
            )
            col = df[[key]]
            na_mask = col[key].isna()
            non_na_col = col[~na_mask]
            # Fit and transform only non-NA rows
            transformed_non_na = encoder.fit_transform(non_na_col).astype(np.float32)
            # Prepare a zero matrix for all rows
            transformed = np.zeros((len(col), transformed_non_na.shape[1]), dtype=np.float32)
            # Fill in the transformed values for non-NA rows
            transformed[~na_mask.to_numpy(), :] = transformed_non_na
            self.encoders[key] = encoder
            self.encoded_labels[key] = encoder.get_feature_names_out().tolist()
            transformed_matrices.append(transformed)
        return transformed_matrices

    def transform(self, df: pd.DataFrame) -> List[Float32Array]:
        if not isinstance(df, pd.DataFrame):
            raise TypeError("adata.obs must be a pandas DataFrame.")

        transformed_matrices: List[Float32Array] = []
        # mask_matrices: List[Float32Array] = []

        for key in self.covariate_keys:
            if key in self.encoders:
                col = df[[key]]
                na_mask = col[key].isna()
                non_na_col = col[~na_mask]
                encoder = self.encoders[key]
                # Transform only non-NA rows
                transformed_non_na = encoder.transform(non_na_col).astype(np.float32)  # type: ignore
                # Prepare a zero matrix for all rows
                transformed = np.zeros((len(col), transformed_non_na.shape[1]), dtype=np.float32)
                # Fill in the transformed values for non-NA rows
                transformed[~na_mask.to_numpy(), :] = transformed_non_na
                transformed_matrices.append(transformed)
        return transformed_matrices
```
