## Supplemental Code for "Interpretable phenotype decoding from multi-condition sequencing data with ALPINE": sampling.py

```
import torch
from torch.utils.data import WeightedRandomSampler
from sklearn.utils.class_weight import compute_sample_weight
from typing import List

def generate_epoch_indices(
    joint_labels: List[str], sampling_method: str, device: torch.device, **kwargs
) -> torch.Tensor:
    total_samples: int = len(joint_labels)

    if sampling_method == "weighted":
        return _get_balanced_epoch_indices(joint_labels, total_samples, device)
    elif sampling_method == "random":
        return torch.randperm(total_samples, device=device)
    else:
        raise ValueError(f"Unknown sampling method: {sampling_method}. Only 'weighted', and 'random' are supported.")

def _get_balanced_epoch_indices(
    joint_labels: List[str], total_samples: int, device: torch.device
) -> torch.Tensor:
    
    # Compute sample weights
    sample_weights = compute_sample_weight(class_weight="balanced", y=joint_labels)

    # Create weighted sampler
    sampler = WeightedRandomSampler(
        weights=sample_weights,  # type: ignore
        num_samples=total_samples,  # Sample exactly total_samples for full epoch
        replacement=True,  # avoid rare labels; if replacement is false might raise error
    )

    # Convert sampler to indices tensor
    return torch.tensor(list(sampler), device=device, dtype=torch.long)


def create_joint_labels_from_dummy_matrices(
    Ys: List[torch.Tensor]
) -> List[str]:
    joint_labels = []
    total_samples = Ys[0].shape[1]
    for sample_idx in range(total_samples):
        sample_labels = []

        for target_idx, dummy_matrix in enumerate(Ys):
            # Get the column for this sample from the dummy matrix
            sample_col = dummy_matrix[:, sample_idx]  # Shape: [num_categories]

            # Find which category is active (argmax of one-hot)
            active_category = torch.argmax(sample_col).item()
            sample_labels.append(f"cov{target_idx}_label{active_category}")

        # Join all target labels for this sample
        joint_labels.append("+".join(sample_labels))

    return joint_labels


def get_batch_indices(
    epoch_indices: torch.Tensor, batch_num: int, batch_size: int
) -> torch.Tensor:
    batch_start = batch_num * batch_size
    batch_end = min(batch_start + batch_size, len(epoch_indices))

    if batch_start >= len(epoch_indices):
        return torch.empty(0, device=epoch_indices.device, dtype=torch.long)

    return epoch_indices[batch_start:batch_end]


def get_num_batches(total_samples: int, batch_size: int) -> int:
    return (total_samples + batch_size - 1) // batch_size  # ceiling division
```
