## Supplemental Code for "Interpretable phenotype decoding from multi-condition sequencing data with ALPINE": LICENSE

BSD 3-Clause License

Copyright (c) 2024, ylaboratory

Redistribution and use in source and binary forms, with or without
modification, are permitted provided that the following conditions are met:

1. Redistributions of source code must retain the above copyright notice, this
   list of conditions and the following disclaimer.

2. Redistributions in binary form must reproduce the above copyright notice,
   this list of conditions and the following disclaimer in the documentation
   and/or other materials provided with the distribution.

3. Neither the name of the copyright holder nor the names of its
   contributors may be used to endorse or promote products derived from
   this software without specific prior written permission.

THIS SOFTWARE IS PROVIDED BY THE COPYRIGHT HOLDERS AND CONTRIBUTORS "AS IS"
AND ANY EXPRESS OR IMPLIED WARRANTIES, INCLUDING, BUT NOT LIMITED TO, THE
IMPLIED WARRANTIES OF MERCHANTABILITY AND FITNESS FOR A PARTICULAR PURPOSE ARE
DISCLAIMED. IN NO EVENT SHALL THE COPYRIGHT HOLDER OR CONTRIBUTORS BE LIABLE
FOR ANY DIRECT, INDIRECT, INCIDENTAL, SPECIAL, EXEMPLARY, OR CONSEQUENTIAL
DAMAGES (INCLUDING, BUT NOT LIMITED TO, PROCUREMENT OF SUBSTITUTE GOODS OR
SERVICES; LOSS OF USE, DATA, OR PROFITS; OR BUSINESS INTERRUPTION) HOWEVER
CAUSED AND ON ANY THEORY OF LIABILITY, WHETHER IN CONTRACT, STRICT LIABILITY,
OR TORT (INCLUDING NEGLIGENCE OR OTHERWISE) ARISING IN ANY WAY OUT OF THE USE
OF THIS SOFTWARE, EVEN IF ADVISED OF THE POSSIBILITY OF SUCH DAMAGE.
