## Supplemental Code for "Interpretable phenotype decoding from multi-condition sequencing data with ALPINE": pyproject.toml

```
[project]
name = "alpine"
version = "0.2.0"
description = "ALPINE is a semi-supervised non-negative matrix factorization (NMF) framework designed to effectively distinguish between multiple phenotypic conditions based on shared biological factors, while also providing direct interpretability of condition-associated genes."
authors = [
    { name = "Wei-Hao Lee", email = "" },
    { name = "Wei-Hao Lee", email = "" },
    { name = "Lechuan Li", email = "" },
    { name = "Ruth Dannenfelser", email = "" },
    { name = "Vicky Yao", email = "" },
]
requires-python = "~=3.10"
readme = "README.md"
dependencies = [
    "scikit-learn>=1.5.2,<2",
    "tqdm>=4.66.5,<5",
    "hyperopt>=0.2.7,<0.3",
    "kneed>=0.8.5,<0.9",
    "torch>=2.5.1,<3",
    "numpy>=2.0.0",
    "pandas>=2.3.1",
    "scanpy>=1.11.4",
    "anndata>=0.11.4",
    "scipy>=1.14.1",
]

[build-system]
requires = ["hatchling"]
build-backend = "hatchling.build"

[dependency-groups]
dev = [
    "ruff>=0.12.8",
]
```
