## Supplemental Code for "Interpretable phenotype decoding from multi-condition sequencing data with ALPINE": README.md

<img src="assests/alpine_icon.png" alt="ALPINE icon" width="200" height="200"/>

# ALPINE

**ALPINE: Adaptive Layering of Phenotypic and Integrative Noise Extraction**

`ALPINE` is a semi-supervised non-negative matrix factorization (NMF) framework designed to effectively distinguish between multiple phenotypic conditions based on shared biological factors, while also providing direct interpretability of condition-associated genes. The entire package is developed in Python and supports GPU usage, significantly enhancing computational speed.

ALPINE can be useful for:

- Identifying condition-associated genes and cells.
- Studying the biological functions of condition-related genes.
- Removing batch effects from the data.

The ALPINE preprint is now available; please review the article at [link](https://www.biorxiv.org/content/10.1101/2025.02.15.638471v1).

> [!NOTE]
> 1. The `get_normalized_expression` function now saves the normalized counts in `adata.layers["normalized_expression"]` instead of `adata.obsm["normalized_expression"]`.
> 2. The `bayesian_search` method in the `ComponentOptimizer` class has been renamed to `search_hyperparams`.
> 3. Rename the `get_conditional_gene_scores()` to `get_covariate_gene_scores()` and it also supports the anndata as input then automatically save into the given anndata.
> 4. The `gpu` argument in both `ALPINE` and `ComponentOptimizer` has been replaced with a `device` argument. Users can now manually specify the device to use (e.g., `"cpu"`, `"cuda"`, or `"mps"`). macOS users with Apple Silicon can try `"mps"` to leverage the GPU on M-series chips.

**Contents**:

- [ALPINE](#alpine)
  - [Installation](#installation)
  - [Usage](#usage)
    - [1. Optimization](#1-optimization)
    - [2. Multi-condition disentangle using ALPINE](#2-multi-condition-disentangle-using-alpine)
      - [a. Training the model](#a-training-the-model)
      - [b. Get the decomposed matrices and counts](#b-get-the-decomposed-matrices-and-counts)
  - [Easy transfer to Seurat object](#easy-transfer-to-seurat-object)
  - [More usage, and analysis](#more-usage-and-analysis)
  - [Citation](#citation)

## Installation

Currently, `ALPINE` is not yet available on the PyPI repository. However, users can clone the entire repository and install the package in their environment by running:

```shell
# download the repo
git clone https://github.com/ylaboratory/ALPINE.git
cd ALPINE

# you can install it in your python environment via your favorite package manager
conda activate your_env_name
# mamba activate your_env_name

# in the alpine folder
pip install .
```

**Important:**
ALPINE is implemented in [PyTorch](https://pytorch.org/), the `pyproject.toml` has included the installation of it. However, if user has issued with the `Pytorch` version. Users will need to install PyTorch separately to ensure compatibility with their specific CUDA or CPU version.

## Usage

The input data type for `ALPINE` is the `AnnData` format. Users should note that ALPINE's model is based on an NMF structure, which supports only **non-negative** values. ALPINE can be trained using either the entire gene list or a selection of highly variable genes.

ALPINE consists of two primary components: optimization and training. The associated classes can be imported as follows:

```python
from alpine import ALPINE, ComponentOptimizer
```

### 1. Optimization

ALPINE integrates a Bayesian optimizer to efficiently search for all necessary hyperparameters, allowing users to easily apply the optimized parameters directly within ALPINE.

```python
from alpine import ComponentOptimizer

# create optimization object with data and covariate keys
co = ComponentOptimizer(adata, covariate_keys=["cov_1", "cov_2"])

# start searching with given parameter range
params = co.search_hyperparams(
    n_total_components_range=(50, 100), 
    alpha_W_range=(0, 1),
    orth_W_range=(0, 0.5),
    l1_ratio_range=(0, 1),
)
```

- `covariate_keys` specifies the categorical columns in `adata.obs` that will be used as covariates.
- `n_total_components_range` sets the range for the total number of components, including `n_components` for unguided embeddings and `n_covariate_components` for guided embeddings.
- `lam_power_range` defines the range for lambda values, spanning from \(10^1\) to \(10^5\).
- `orth_W_range`: The range for the orthogonal weight regularization on the \( W \) matrix, designed to encourage gene signatures to capture distinct patterns.
- `l1_ratio_range`: The range for the L1 ratio, controlling the balance between L1 (LASSO) and L2 (ridge) regularization.
- `alpha_W_range`: The range for the regularization weight on the \( W \) matrix, determining the influence of LASSO and ridge regularization on \( W \).


The `ComponentOptimizer` class offers a range of convenient and practical functions to help users monitor and extend their training process. See the analysis section below for more details.

### 2. Multi-condition disentangle using ALPINE

With ALPINE, you have the flexibility to either manually define the parameters you want to use or apply the optimized parameters learned from previous steps.

#### a. Training the model

1. **Manually specified parameters:**

```python
# user can maually specify desired paramteres
alpine_model = ALPINE(
    n_components = 30,
    n_covariate_components = [5, 5] 
    alpha_W = 0,
    lam = [1e+3, 1e+3],
    device = "cuda"
)
alpine_model.fit(adata, covariate_keys=["cov_1", "cov_2"])
```

2. **Using optimized parameters from `ComponentOptimizer` (Recommend):**

```python
# if you use the ComponentOptimizer, you can simply plugin the parameteres learned from the last step
alpine_model = ALPINE(**param)
alpine_model.fit(adata, covariate_keys=["cov_1", "cov_2"])
```

Finally, users can save the trained embeddings directly into `adata` by running:

```python
alpine_model.store_embeddings(adata)

# the H embedding can be retrieved by
alpine_model.obsm["ALPINE_embedding"] # unguided embedding
alpine_model.obsm["cov_1"] # covariate embedding
alpine_model.obsm["cov_2"] # covariate embedding

# the W embedding
alpine_model.varm["ALPINE_embedding"] # unguided gene signature embedding
alpine_model.varm["cov_1"] # covariate gene signature embedding
alpine_model.varm["cov_2"] # covariate gene signature embedding

```

#### b. Get the decomposed matrices and counts

In addition to obtaining embeddings from `adata`, users can also retrieve the decomposed matrices from the `ALPINE` model by using:

```python
Ws, Hs, Bs = alpine_model.get_decomposed_matrices()
```

The order of the matrices in `Ws` and `Hs` follows the sequence of the covariate keys, with the unguided embeddings placed at the end. In contrast, the `Bs` matrices do not include the unguided portion; their order strictly adheres to the covariate keys.

To obtain the normalized counts that are free from batch effects and conditions, where:

```python
alpine_model.get_normalized_expression(adata)

# the normalized counts is in here
adata.layers["normalized_expression"]
```

There are additional applications for our model; please refer to the next section for more details.

## Easy transfer to Seurat object

In Python, we usually store the `anndata` into `h5ad` format. To easily transfer the `h5ad` to Seurat object, this requires three packages installed in R. If want to check the full tutorial, please visit our tutorial repo [here](https://github.com/ylaboratory/ALPINE-analysis/blob/master/tutorials/transfer_from_anndata_to_seurat.Rmd).

```R
if (!requireNamespace("BiocManager", quietly = TRUE))
    install.packages("BiocManager")
BiocManager::install(c("zellkonverter", "SingleCellExperiment", "Seurat"))
```

Then we can load the `h5ad` format through the following commands, and we could quickly examine through the data is intact.

```R
adata <- readH5AD("your_file.h5ad")

# check the adata
assayNames(adata)   # e.g. "X", "counts", "normalized_expression"
reducedDimNames(adata) # e.g. guided and unguided embeddings: "ALPINE_embedding", "batch", "condition"
colnames(colData(adata)) # cell metadata
rownames(rowData(adata)) # gene metadata
```

Then we can create the Seurat Object, and move the ALPINE embeddings and gene signatures into the object.

```R
# createt Seurat object
raw_counts <- assays(adata)[["counts"]]  # or "X" if that's your raw
seurat_obj <- CreateSeuratObject(counts = raw_counts, project = "ALPINE_demo")

# store the ALPINE counts
ALPINE_counts <- assays(adata)[["normalized_expression"]]
seurat_obj[["normalized_expression"]] <- CreateAssayObject(counts = ALPINE_counts)

# add cell metadata
cell_metadata <- as.data.frame(colData(adata))
seurat_obj <- AddMetaData(seurat_obj, metadata = cell_metadata)

# add the ALPINE embeddings and weights
gene_loadings <- adata@rowRanges@elementMetadata@listData$varm@listData$ALPINE_weights
seurat_obj[["ALPINE"]] <- CreateDimReducObject(
    embeddings = reducedDims(adata)$ALPINE_embedding,
    loadings = gene_loadings,
    key = "ALPINE_",
    assay = DefaultAssay(seurat_obj)
)
```

## More usage, and analysis

All analyses from the papers and case studies are stored in the ALPINE-analysis repository, where you can access a variety of resources. Additionally, the repository provides valuable tips for tuning the model.

- [ALPINE-anlaysis repository](https://github.com/ylaboratory/ALPINE-analysis) provdes following tutorials:
  - Save, load, and extend the optimization process.
  - Retrieve condition-associated gene scores.
  - Transform new, unseen data using the trained model.

## Citation

If you use our tool in your study, please cite the following paper:
> Lee WH, Li L, Dannenfelser R, Yao V. Interpretable phenotype decoding from multi-condition sequencing data with ALPINE. bioRxiv. 2025:2025-02.

<!-- ## Overview of ALPINE

![overview](./imgs/figure1.png) -->
