## Supplementary figures and images for "Interpretable phenotype decoding from multi-condition sequencing data with ALPINE"

### alpine_icon.png

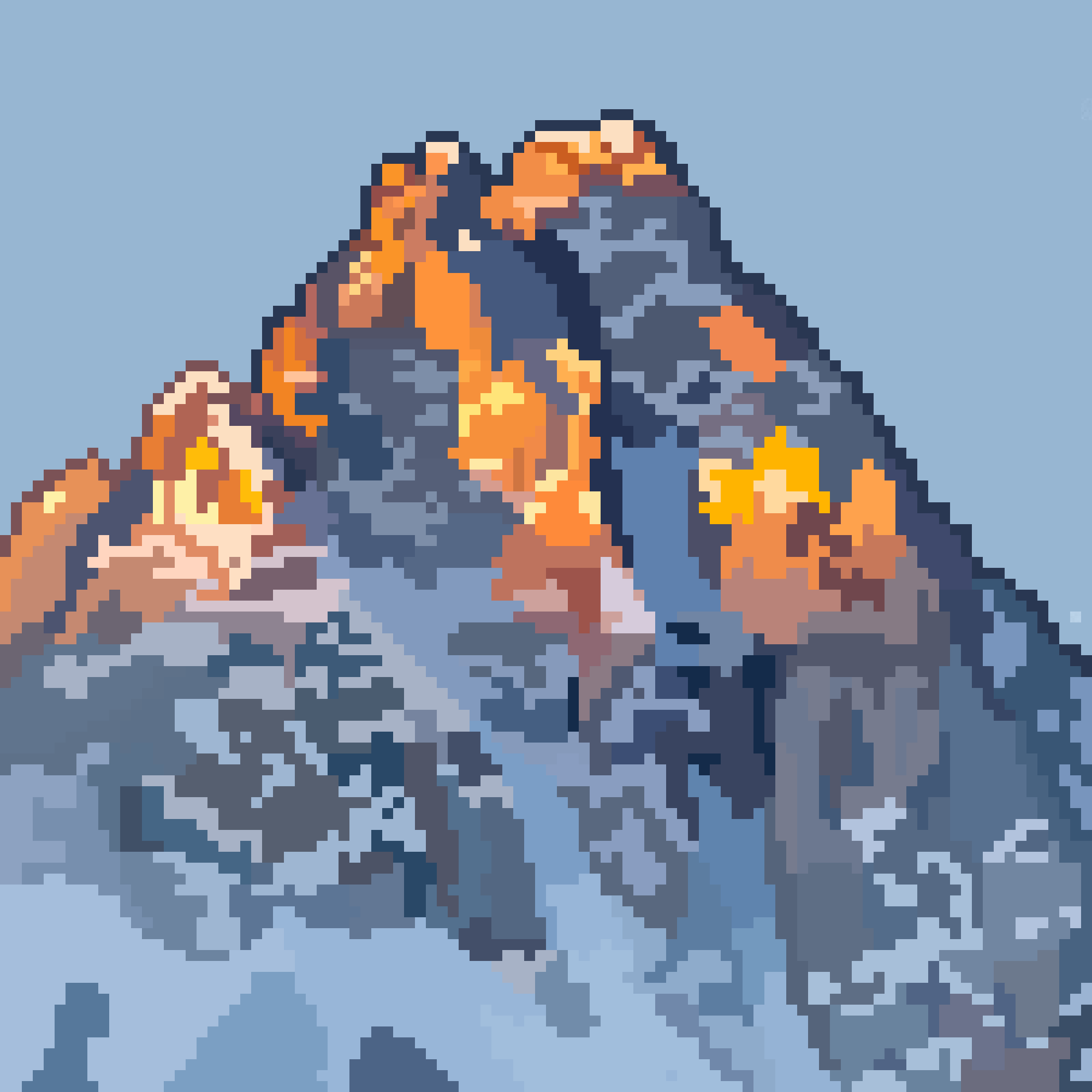

### figure1.png

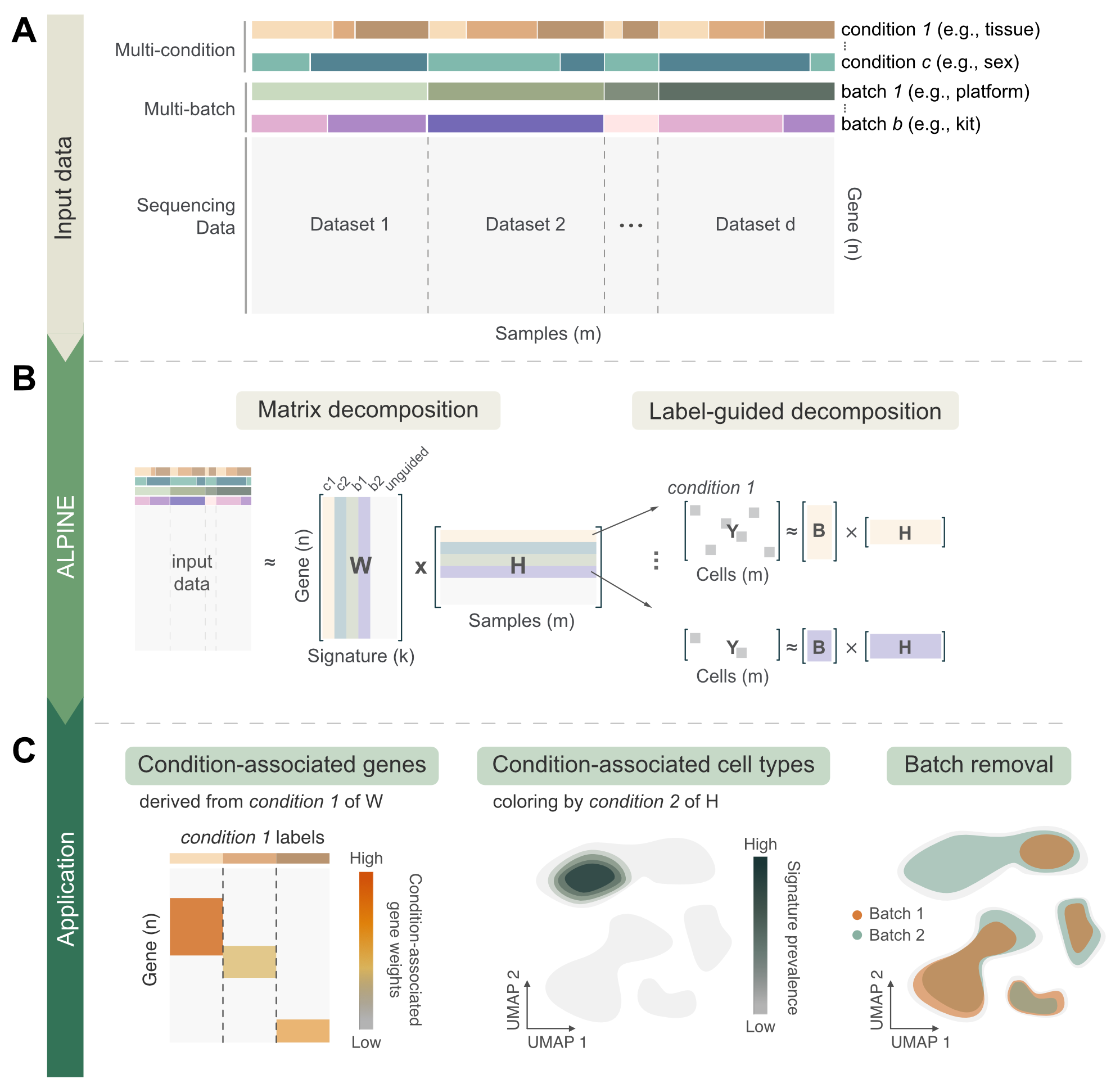
